# The parasitic lifestyle of an archaeal symbiont

**DOI:** 10.1101/2023.02.24.529834

**Authors:** Joshua N. Hamm, Yan Liao, Andriko von Kügelgen, Nina Dombrowski, Evan Landers, Christopher Brownlee, Emma M. V. Johansson, Renee M. Whan, Matthew A. B. Baker, Buzz Baum, Tanmay A. M. Bharat, Iain G. Duggin, Anja Spang, Ricardo Cavicchioli

## Abstract

DPANN Archaea are a diverse group of organisms typically characterised by small cells and reduced genomes. To date, all cultivated DPANN Archaea are ectosymbionts that require direct cell contact with an archaeal host species for proliferation. However, the dynamics of DPANN – host interactions and the impacts of these interactions on the host species are poorly understood. Here, we show that one DPANN archaeon (*Candidatus* Nanohaloarchaeum antarcticus) engages in parasitic interactions with its host (*Halorubrum lacusprofundi*) that result in host cell lysis. Our data also suggest that these interactions involve invasion of the host cell by the nanohaloarchaeon. This is the first reported instance of such a predatory-like lifestyle amongst Archaea and indicates that some DPANN Archaea may interact with host populations in a manner similar to viruses.

## Introduction

DPANN Archaea are extremely diverse and thought to comprise various ectosymbiotic Archaea^1-6^. Initially comprised of the phyla <u>D</u>iapherotrites, <u>P</u>arvarchaeota, <u>A</u>enigmarchaeota, <u>N</u>anoarchaeota, and <u>N</u>anohaloarchaeota (from which the acronym is derived), the DPANN superphylum is now assumed to include the Woesearchaeota, Huberarchaeota, Pacearchaeota, Mamarchaeota, Micrarchaeota, Altiarchaeota (predicted to be free-living), and Undinarchaeota^7,8^. DPANN Archaea have been identified in a diverse range of environments including acidic hot springs^2^, deep-sea sediment^3^, acid mine drainage^4^, microbial mats^3^, the human microbiome^6^ aquifers^9^ and marine waters^8,10^, as well as Antarctic lakes^1^ and hypersaline systems that harbour, among others^11,12^, Nanohaloarchaeota^5^ To date, all successfully cultivated DPANN Archaea are symbionts, requiring direct cell contact with another archaeon for proliferation^9^. Cultivation of DPANN has proven difficult, with only three lineages currently represented in published laboratory co-cultures: the Nanoarchaeota^2,13,14^, the Micrarchaeota^15,16^, and the Nanohaloarchaeota^1,17,18^. Thus far, the processes by which DPANN cells associate with their hosts and proliferate are poorly understood^9^.

The ectoparasitic interactions between *Ignicoccus hospitalis* and *Nanoarchaeum equitans* (the best characterised DPANN-host system) appears to involve membrane fusion between *N. equitans* and *I. hospitalis* generating a channel that connects the cytoplasms of both organisms^19^. ‘Cytoplasmic bridges’ have been observed between other DPANN and their hosts and are thought to facilitate nutrient transfer^1,20,21^. However, the proteins forming or catalysing the formation of these structures are unknown^21^. Moreover, many DPANN appear to engage in interactions without forming such structures^16,17^, and the mechanism by which they acquire nutrients is unclear. Furthermore, the process of cell proliferation is poorly understood with some DPANN (*e.g.* Nanoarchaeota) remaining predominantly associated with their hosts^2^, while others (*e.g.* Nanohaloarchaeota) produce large quantities of dissociated cells^1,21^. Thus, much remains to be learned about how DPANN attach, proliferate and detach.

Recently, the Antarctic DPANN archaeon, *Candidatus* Nanohaloarchaeum antarcticus was discovered to require the haloarchaeon *Halorubrum lacusprofundi* for growth^1^. Here we report the results of live fluorescence, cryogenic correlative light and electron microscopy, and electron cryotomography demonstrating that *Ca.* Nha. antarcticus cells interact with *Hrr. lacusprofundi* cells, appear to internalise within the host cytoplasm, and subsequently trigger lysis of the host.

## Results and Discussion

### Dynamics of *Ca.* Nha. antarcticus – *Hrr. lacusprofundi* interactions

Our enrichment culture predominantly consisting of the symbiont *Ca.* Nanohaloarchaeum antarcticus and several *Halorubrum lacusprofundi* host strains as well as a *Natrinema* species in low abundance (<1%)^1^ offers an ideal system for studying archaeal symbiosis. Specifically, it can be used to produce large numbers of nanohaloarchaeal cells (so that they make up ∼50% of total cells in a co-culture at ∼10^8^ cells mL^-1^), which can be isolated and used to infect a single host strain. To investigate *Ca*. Nha antarcticus – *Hrr. lacusprofundi* interaction dynamics, we used MitoTracker fluorescent dyes^22^ as vital cell stains to identify and track live interactions between *Ca.* Nha. antarcticus and *Hrr. lacusprofundi* strain R1S1^23^ as well as electron microscopy to investigate morphological features.

Purified *Ca*. Nha. antarcticus cells were stained with MitoTracker DeepRed (MTDeepRed) followed by incubation with MitoTracker Orange (MTOrange)-stained Hrr. lacusprofundi. Live co-cultures of labelled cells were then immobilized and cultured on an agarose gel pad or in a microfluidic flow chamber and imaged using time-lapse fluorescence microscopy, 3D laser scanning confocal microscopy, and 4D (3D time-lapse) live cell imaging. In agreement with previous work on other haloarchaeal species^22^, our data shows that these Mitotracker dyes are retained by *Hrr. lacusprofundi* cells, and do not affect cell growth rates (Fig. S4, S9 and S11).

A total of 163 MTOrange-stained *Hrr. lacusprofundi* cells were analysed in detail during two incubations of 24 hrs on agarose pads. Of these, 132 (81%) were observed with one or more MTDeepRed-stained *Ca.* Nha. antarcticus cell(s) attached at the first timepoint imaged (0 h), indicating that attachment predominantly occurred during the initial incubation period (≤ 1 h) prior to commencement of time-lapse imaging. Over time, *Ca.* Nha. antarcticus cells appeared to migrate from the exterior to the interior of *Hrr. lacusprofundi* cells (Fig. 1, S1). Confocal imaging with 3D-orthogonal projection after 10 h incubation showed *Ca.* Nha. antarcticus cells that appeared fully internalised within *Hrr. lacusprofundi* (Fig. 1b, S2). Migration of nanohaloarchaeal cells typically took several hours but the exact duration varied between different observed interactions (Fig. 1, S1).

**Fig. 1.**
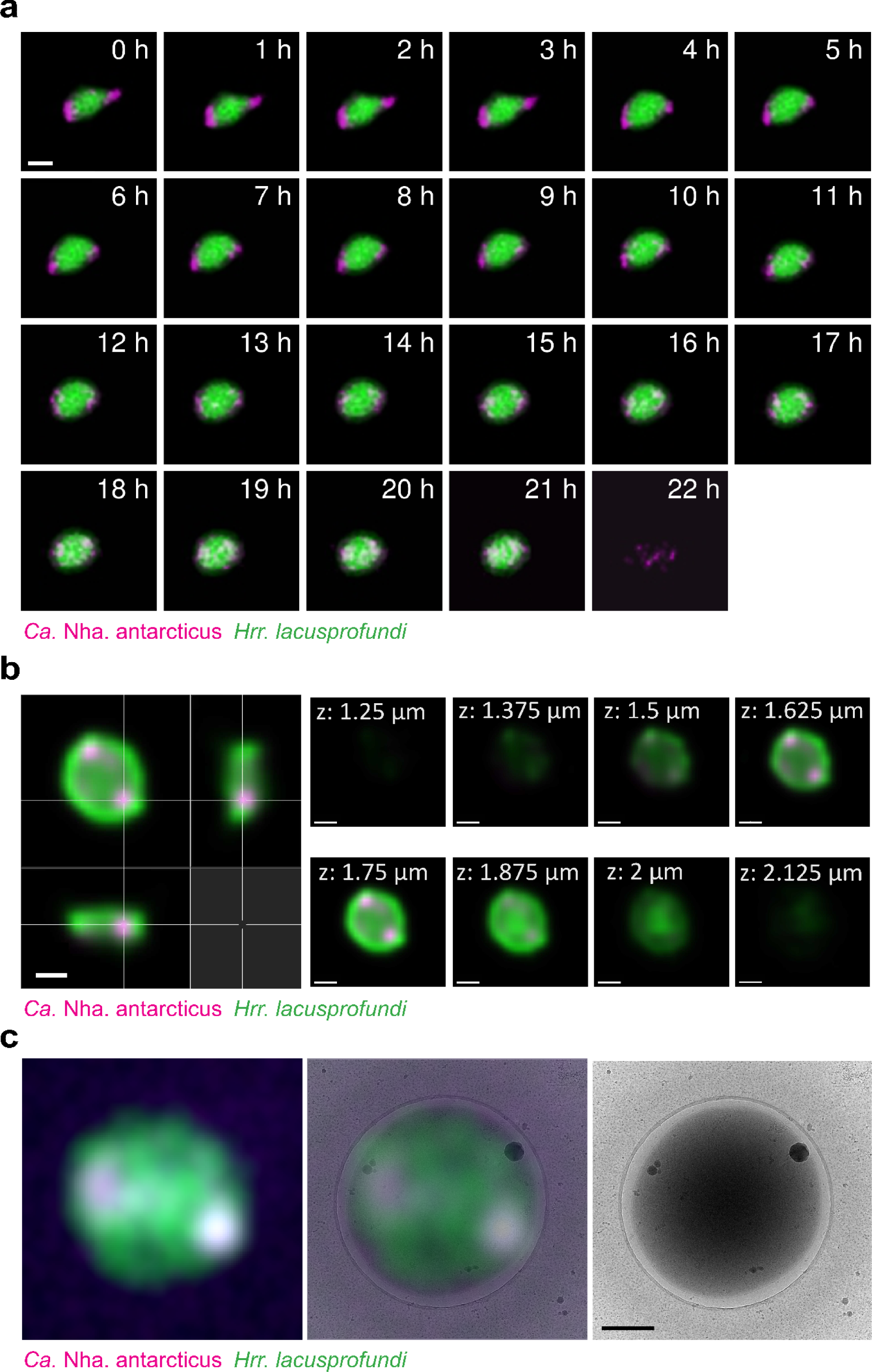
Live fluorescence demonstrates *Ca.* Nha. antarcticus enters *Hrr. lacusprofundi* cells. **a**, Representative live fluorescence time lapse series of *Ca.* Nha. antarcticus cells (MitoTracker DeepRed, coloured Magenta) attached to a host *Hrr. lacusprofundi* cell (MitoTracker Orange, coloured Green) (0 – 9 h), migrating internally (∼10 – 21 h), followed by lysis of the host (22 h). **b**, Confocal 3D-orthogonal slice maximum intensity projection showing representative images of *Ca.* Nha. antarcticus cells internalised within *Hrr. lacusprofundi* after 10 h incubation. **c,** Cryo-CLEM of internalised *Ca.* Nha. antarcticus cells (stained with MitoTracker DeepRed, represented Magenta) within a *Hrr. lacusprofundi* cell (stained with MitoTracker Orange, represented Green). Overlay of fluorescence and cryo-TEM data shows the host cell remains intact following internalisation and no sign of the *Ca.* Nha. antarcticus cells on the exterior of the *Hrr. lacusprofundi* cell. Scale bars: **a** – 1 µm, **b** and **c** – 500 nm.

Following migration, the area occupied by the *Ca.* Nha. antarcticus dye within the bounds of the *Hrr. lacusprofundi* cell expanded over time (Fig. 1a, S1, and S3). Over the course of the 24 hr incubation period, 27% (36/132) of the *Hrr. lacusprofundi* cells that were observed with attached *Ca.* Nha. antarcticus cell(s) lysed, accounting for 22% (36/163) of total *Hrr. lacusprofundi* cells in co-cultures (Fig. 1a, S1, 3). Lysis occurred relatively rapidly and was complete within the 30 min time window separating image acquisitions. No lysis was observed over periods of up to 70 h in control samples of pure host cells (Fig. 1a, S1, S3, and S4). Upon lysis, the dye used to label host cells in co-cultures dissipated completely, whereas labelled *Ca.* Nha. antarcticus cells remained apparently unaffected (e.g., Fig. 1a, compare 21 h and 22 h). These results are consistent with *Ca*. Nha. antarcticus growth and survival upon host cell lysis.

**Fig. 2.**
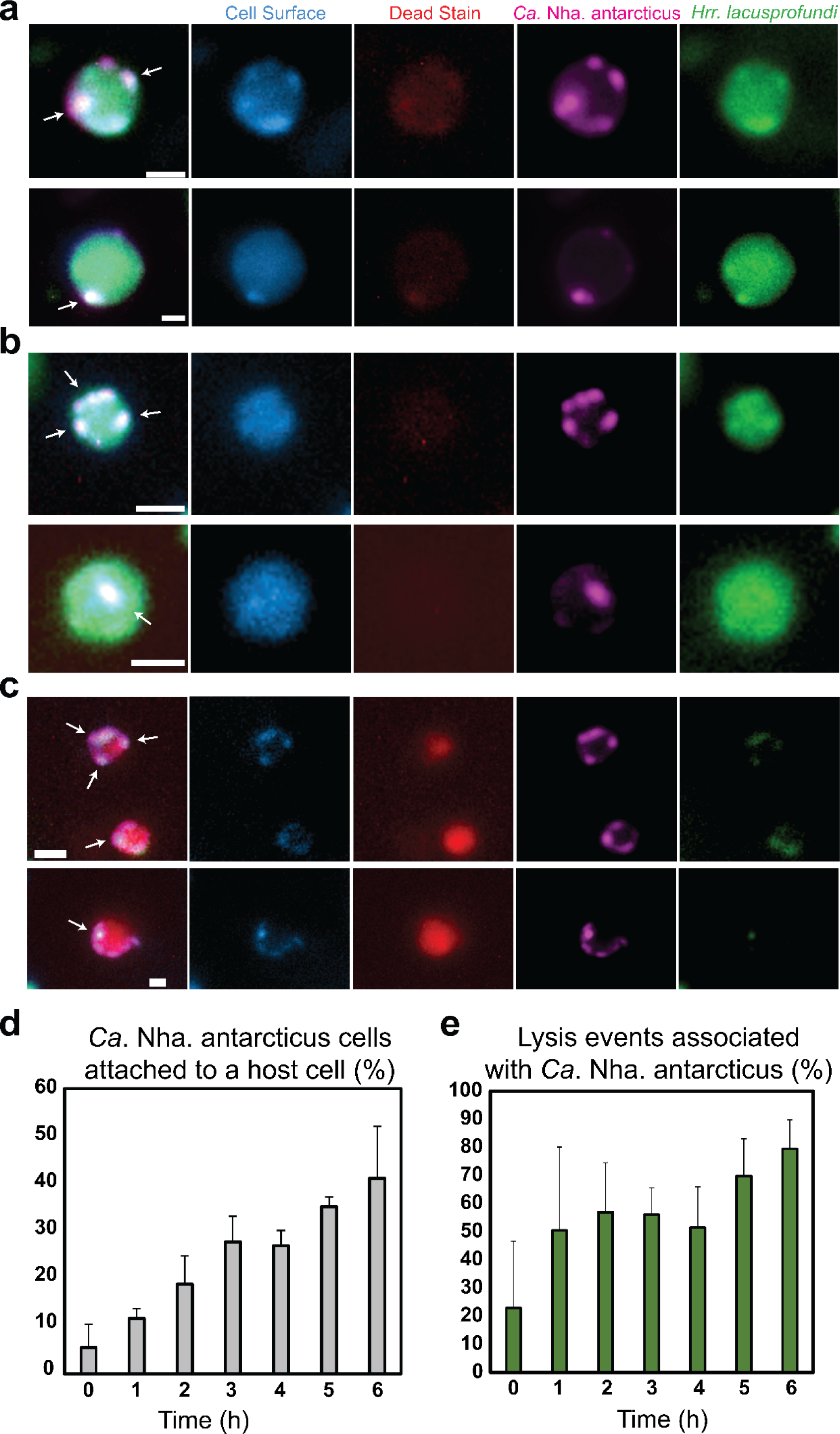
Fluorescence microscopy of *Ca.* Nha. antarcticus induced cell lysis. **a - c** Live fluorescence micrographs taken 6 h post-mixing showing *Ca.* Nha. antarcticus (MitoTracker Green, coloured Magenta) interactions with *Hrr. lacusprofundi* (MitoTracker Orange, coloured Green) including additional stains for cell surface (ConA-AF350, coloured Blue), and cell death (RedDot 2, coloured red). **a,** Representative fluorescence micrographs showing *Ca.* Nha. antarcticus cells (MitoTracker Green, coloured Magenta) attached to the surface of *Hrr. lacusprofundi* (MitoTracker Orange, coloured Green). Cell surface staining (ConA-AF350, coloured Blue) shows foci corresponding to regions where *Ca.* Nha. antarcticus was attached to the host cell. No signs of lysis were detected by a dead cell stain (RedDot 2, coloured Red). **b,** Representative live fluorescence micrographs showing *Ca.* Nha. antarcticus cells (stained with MitoTracker Green, represented Magenta) internalised within *Hrr. lacusprofundi* cells (stained with MitoTracker Orange, represented Green). Cell surface staining (Concanavalin A, represented Blue) does not show foci corresponding to *Ca.* Nha. antarcticus cells, indicating the symbiont has become internalised. No signs of lysis are evident from inclusion of a dead stain (RedDot 2, represented Red). **c,** Representative fluorescence of *Hrr. lacusprofundi* (MitoTracker Orange, coloured Green) lysis events associated with *Ca.* Nha. antarcticus (MitoTracker Green, coloured Magenta). Lysis is indicated by positive fluorescence for RedDot 2 (coloured Red) and is associated with loss of MitoTracker Orange signal from the host cell while the *Ca.* Nha. antarcticus cells remain intact and positive for both MitoTracker Green and the cell surface stain (Con-AF350A, coloured Blue). **d** and **e,** Quantitative data for **d**) lysis and **e**) attachment events over short-term incubations. Data show **d**) percentage of lysis events associated with a *Ca.* Nha. antarcticus cell and **e**) the percentage of *Ca.* Nha. antarcticus cells attached to host cells over the course of a time series (0 – 6 h). Data show average number of events across triplicate experiments, and error bars represent standard deviation as summarised in Supplementary Table 2. Arrows: examples of *Ca.* Nha. antarcticus cells; Scale bars: 500 nm.

**Fig. 3.**
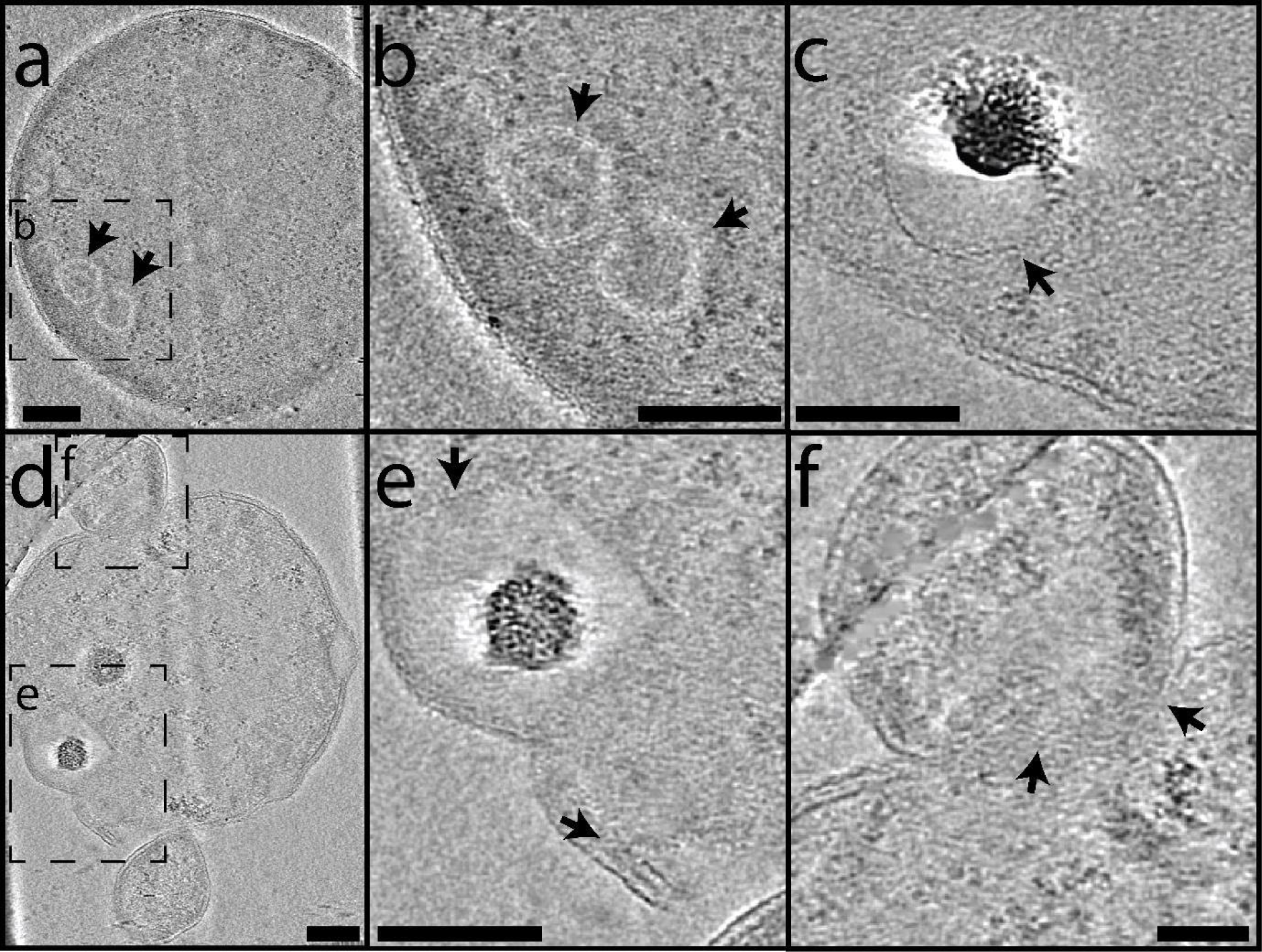
Cryo-electron tomography of *Ca.* Nha. antarcticus – *Hrr. lacusprofundi* co-cultures show internal membrane bound structures. **a – f**, Z-slices from tomograms (**a** and **b:** Supplementary Video 2, **c:** Supplementary Video 3, **d** - **f:** Supplementary Video 4) showing (**a** – **d**) internal membrane bound structures within *Hrr. lacusprofundi* cells consistent with *Ca.* Nha. antarcticus that have internalised. **a – b,** A *Hrr. lacusprofundi* cell with an intact cell membrane and multiple internal membrane bound structures (arrows). **c,** A *Hrr. lacusprofundi* cell with an intact cell membrane and an internal membrane bound structure (arrow). (**d** – **f**) A *Hrr. lacusprofundi* cell with internal structures consistent with internalised *Ca*. Nha. antarcticus cells (**e**, arrows) and a *Ca*. Nha. antarcticus cell that appears to be in the process of internalising (**f**, arrows indicate region where *Ca.* Nha. antarcticus cell appears to be partially within the *Hrr. lacusprofundi* cell). Images acquired using a Titan Krios at 300 kV. Scale bars: 100 nm

To further investigate whether migration of the *Ca.* Nha. antarcticus cells into the *Hrr. lacusprofundi* cells corresponded to complete internalisation, a lectin cell surface stain (Concanavalin A conjugated with Alexa Fluor 350 (ConA-AF350)) and a live-cell-impermeable stain (RedDot 2) were used to label surface bound nanohaloarchaeal cells and to assess loss of host cell membrane integrity, respectively. As expected, ConA-AF350 added to co-cultures labelled *Ca.* Nha. antarcticus cells that were attached to the surface of *Hrr. lacusprofundi* (Fig. 2a, S5). By contrast, when ConA-AF350 was added to co-cultures at later time-points (6h), many *Ca.* Nha. antarcticus cells did not appear positive for ConA-AF350 (Fig. 2b, S6), consistent with host cell invasion. The absence of RedDot 2 staining indicated that these host cells had remained intact during the internalisation process (Fig. 2b, S6). As expected, host cells that were inferred to have lysed via the loss of MitoTracker Orange signal stained positive for RedDot 2 (Fig. 2c, S6). Over the course of 0 – 6 h incubations, the proportion of *Hrr. lacusprofundi* lysis events associated with *Ca.* Nha. antarcticus cells increased from ∼23% to ∼80% while the proportion of nanohaloarchaeal cells attached to a host cell increased from ∼6% to ∼41% (Fig. 2d and e, Supplementary Table 2). Throughout, *Ca.* Nha. antarcticus cells associated with lysed *Hrr. lacusprofundi* cells labelled positive for both MitoTracker and Concanavalin A stains (Fig. 2c, S7).

To complement this analysis, similar experiments were performed using continuous liquid flow culture (in a microfluidics system) to assess the interactions of immobilised, MTOrange stained *Hrr. lacusprofundi* R1S1 cells (0.7 – 1.1 μm trap height) with MTDeepRed stained, FACS-purified, *Ca.* Nha. antarcticus cells (Fig. S3, Supplementary Video 1). As with the agarose pad experiments, *Ca.* Nha. antarcticus cell(s) were observed attaching to *Hrr. lacusprofundi* cells at the time of first imaging. Again, over a 2 – 23 h time-period, decreased signal intensity and increased area were observed for the internalised *Ca.* Nha. antarcticus cells that were consistent with growth of *Ca.* Nha. antarcticus within their host (Fig. S3). During the time course, 360 *Hrr. lacusprofundi* cells lysed (56%), whereas no lysis occurred in the uninfected and unstained control (407 cells) while 2 lysis events occurred in the uninfected and stained control (654 cells) (Fig. S3 and S9, Supplementary Table 1). Taken together, both the agarose pad and microfluidic experiments demonstrate that *Ca.* Nha. antarcticus cells induce lysis of their hosts (22 – 56% of total *Hrr. lacusprofundi* cells were lysed versus ∼0% in the control, Supplementary Table 1).

Experiments were also performed to investigate the effect of *Ca.* Nha. antarcticus on the morphology of *Hrr. lacusprofundi*, which includes rods and coccoid cells (Fig. S9, Supplementary Table 1 and ref. ^24,25^). After co-incubation with MTDeepRed-stained *Ca*. Nha. antarcticus, 34% rod-shaped MTOrange-stained *Hrr. lacusprofundi* cells (on agarose pad) had changed to a more rounded shape (Fig. S1 and S3) compared to stable morphology of control *Hrr. lacusprofundi* cells (Fig. S4, Table S1). A higher proportion of morphological changes of co-cultured *Hrr. lacusprofundi* compared to pure culture was also seen with the microfluidics experiments (Fig. S9, Supplementary Table 1).

These results suggest that the association of the two species impacts the structure of the host cell envelope. To investigate this, Cryo-CLEM (cryo-correlated light and electron microscopy) was performed to obtain high-resolution images of the fluorescently labelled cells (Fig. 1c, S12). Cells were fluorescently labelled as for live fluorescence imaging, and the labelled *Ca.* Nha. antarcticus and *Hrr. lacusprofundi* cells incubated together for 20 h to enable attachment and invasion prior to vitrification by plunge-freezing and imaging. *Hrr. lacusprofundi* cells were observed that also appeared co-labelled for the MTDeepRed used to stain *Ca.* Nha. antarcticus. This included examples where MTDeepRed fluorescence was confined to discrete regions of the host cell as well as ones in which the fluorescent signal was present throughout the host cell (Fig. 1c, S12). In all observed examples where *Hrr. lacusprofundi* cells were co-stained for the *Ca.* Nha. antarcticus dye, the host cell appeared intact with no obvious disruptions to its cell membrane. This suggests that the initial association, while leading to a change in shape and internalisation, does so without yet causing host cell lysis.

### Structural features of the *Ca.* Nha. antarcticus symbiosis

To investigate structural features of the interactions and apparent internalisation in greater detail, higher quality three-dimensional images of *Hrr. lacusprofundi* and *Ca.* Nha. antarcticus cells were acquired for both pure samples and co-cultures using electron cryotomography (cryo-ET). In pure *Ca.* Nha. antarcticus samples, bulges within the membrane and cytoplasmic structures were observed in *Ca.* Nha. antarcticus cells (Fig. S13). The appearance of the cytoplasmic structures in *Ca.* Nha. antarcticus, with a surface monolayer surrounding a higher electron density core and uniform texture resemble polyhydroxyalkanoate-like (PHA-like) granules which have previously been identified in *Hrr. lacusprofundi*^26^ and may be acquired from the host during interactions. The bulges within the *Ca.* Nha. antarcticus membrane resemble lipid droplets^27^ and were also observed in the membranes of *Hrr. lacusprofundi* cells interacting with *Ca.* Nha. antarcticus cells (Fig. 3d). Interestingly, these were not observed in the membranes of uninfected *Hrr. lacusprofundi* cells (Fig. S15), implying that the Nanohaloarchaeota play a role in inducing the formation of these structures. This is potentially significant as *Ca.* Nha. antarcticus lacks identifiable genes for both lipid biosynthesis and metabolism^1^ and therefore must acquire lipids from the host to survive.

Strikingly, internal membrane-bound structures (∼80 - 250 nm diameter) were observed in several *Hrr. lacusprofundi* cells incubated with *Ca.* Nha. antarcticus and imaged by cryo-ET (Fig. 3, S16). In some cases, these appeared in intact cells (Fig. 3a – c, S16c – d) – consistent with the idea that a population of *Ca.* Nha. enters *Hrr. lacusprofundi* cells without inducing their lysis. In addition, internal structures were seen in cells that appeared damaged (Fig. S16a – b) or to have a disrupted bounding membrane (Fig. 3d – f). In general, the internalised structures were highly radiation sensitive (Fig. S16), limiting the signal to noise ratio of the tomograms that could be reconstructed. However, in many instances, the membrane-bound structures seen in co-cultures had a surface that exhibited a repeating pattern^21,28^ characteristic of an S-layer (Fig. 3b). Archaea and Bacteria are known to possess several mechanisms to prevent S-layer proteins assembling in the cytoplasm^29-31^. In turn, these intracellular structures surrounded by a putative S-layer, are consistent with *Ca*. Nha antarcticus cells that have internalised. While these structures were only seen in co-cultures, high-contrast granules that resemble calcium-phosphate granules^32^ were observed in *Hrr. lacusprofundi* cells from both pure cultures and co-cultures (Fig 3, S14 – 16).

We also used Cryo-ET to examine the contact sites between *Ca.* Nha. antarcticus and *Hrr. lacusprofundi* cells prior to invasion (Fig. S14). In some cases, these images suggested fusion of the two cells membranes and the opening of a cytoplasmic channel – similar to the structure of the interaction interface previously reported for *N. equitans* and *I. hospitalis* cells^19^. In these cases, the S-layers of both organisms appear to be discontinuous at the interaction site (Fig. S14). In all cases, the interaction site was limited to a region of ∼15 – 20 nm in width, despite the close physical association of the two cells. By Cryo-TEM we also observed a single case of a cytoplasmic bridge extending outwards from a *Ca.* Nha. antarcticus cell (Fig. S17), resembling the structure previously described connecting *Ca.* Nha. antarcticus and *Hrr. lacusprofundi* cells based upon FISH and TEM^1^. This cytoplasmic bridge structure shows no discernible barrier between the cytoplasm of *Ca.* Nha. antarcticus and the bridge contents, suggesting that the structure may enable direct cytoplasmic transfer, as has been suggested for other DPANN^20^. However, the host cell to which the bridge extends was not clearly visible and the ultrastructure of the host associated region of the bridge has not been elucidated.

### *Ca.* Nha. antarcticus is a parasitic archaeon

Our data demonstrates that the relationship between *Ca.* Nha. antarcticus and *Hrr. lacusprofundi* is parasitic, with interactions between the two organisms leading to lysis of large numbers of host cells in pure co-cultures which may explain why such pure co-cultures cannot be maintained stably^1^. Fluorescence microscopy approaches demonstrated that initial attachment of nanohaloarchaeal cells is followed by migration of the nanohaloarchaeal signal, with the nanohaloarchaeal cell eventually appearing fully encapsulated by the host cell (Fig. 1, S1). Addition of a cell surface stain revealed that the surfaces of nanohaloarchaeal cells encapsulated by host cells are inaccessible, in contrast to externally attached nanohaloarchaeal cells (Fig. 2, S5). Finally, cryo-ET data revealed the presence of internal, membrane bound structures within *Hrr. lacusprofundi* cells in co-cultures that appeared to be bound by an S-layer and were not observed in pure *Hrr. lacusprofundi* cultures (Fig. 3, S16).

Our data is suggestive of at least a subset of Nha cells invading the host cytoplasm during infection, acquiring nutrients, and being released via lysis, though the molecular mechanisms responsible for this process remain enigmatic. Considering that the size of the internal structures observed with cryo-ET are at the lower limit of external nanohaloarchaeal cells, it may be speculated that these represent vesicles rather than symbiont cells. However, we believe this explanation is unlikely for several reasons; 1) the most common form of intracellular vesicle structure in haloarchaea are gas vesicles^33^ which *Hrr. lacusprofundi* has not been shown to produce and for which it lacks the necessary genes^34^, 2) the host strain used in these experiments is known to produce vesicles^35^ but no similar internal structures were observed in pure host controls, 3) current models for production of other vesicle types in Archaea involve budding of vesicles from the cell surface rather than production of vesicles within the cell^35,36^, 4) multiple internal structures have intact S-layers on the outside of their lipid bilayer despite Archaea possessing several mechanisms to avoid intracellular S-layer assembly^29-31^, and 5) vesicle formation via intracellular budding should lead to either no S-layer, or an inverted bi-layer – S-layer structure. Thus, while we cannot completely discount the possibility of these structures being vesicles, the data seems most compatible with them being nanohaloarchaeal cells.

The theoretical lower limit for cell size is commonly cited as 200 nm diameter to provide sufficient volume for essential cellular functions^37^ but the size of putative *Ca.* Nha. antarcticus cells observed in *Hrr. lacusprofundi* is often lower than this. However, this limit assumes the cell in question is free-living and must perform all essential cellular functions independently^37^, which is not the case for *Ca.* Nha. antarcticus cells during infection of *Hrr. lacusprofundi.* This theoretical limit also assumes that the cells exist at this size for the majority of the cell cycle^37^, whilst in our model for the *Ca.* Nha. antarcticus lifestyle this state would be transient. Notably, recent work on Saccharibacteria TM7i (a parasitic CPR bacterium) has shown that cell division is asymmetric with planktonic or newly divided cells having calculated areas as low as 0.02 µm^2^ (which corresponds to a diameter of ∼160 nm), which was significantly smaller than adhered and pre-divisional cells^38^. The dynamics of cell division in DPANN Archaea are currently unknown but a similar asymmetric process could explain the apparent size of putative *Ca.* Nha. antarcticus cells observed here with cryo-ET.

Our proposed lifestyle for *Ca.* Nha. antarcticus shows similarities to the recently reported lifestyle for *Ca.* Vampirococcus lugosii, a CPR bacterium recently reported to prey on a gammproteobacterium^39^. However, *Ca*. V. lugosii does not appear to invade its host cells but remains in an ectosymbiotic state^39^. Lifestyles involving invasion and proliferation within a host cell are characteristic of viruses and some bacterial predators, most notably *Bdellovibrio bacteriovorus^40^*. Archaea are not known to internalise within other archaeal cells or to be hosts of intracellular symbionts. In fact, the only known examples of archaeal endosymbionts are certain methanogenic Archaea residing within protists^41,42^. As a result, the herein observed parasitic lifestyle of *Ca.* Nha. antarcticus in co-culture with *Hrr. lacusprofundi* is thus far unique among Archaea. In contrast to *B. bacteriovorus,* which invades the periplasm^40^, we provide support for *Ca.* Nha. antarcticus entering the host cytoplasm. Due to differences in host membrane structure both *Ca.* Nha. antarcticus and *B. bacteriovorus* appear to accomplish internalization by crossing a single lipid bilayer without destabilising the host cell. A simplified putative life cycle for *Ca.* Nha. antarcticus is depicted in Fig. 4 but further *in-situ* studies will be needed to confirm this.

**Fig. 4.**
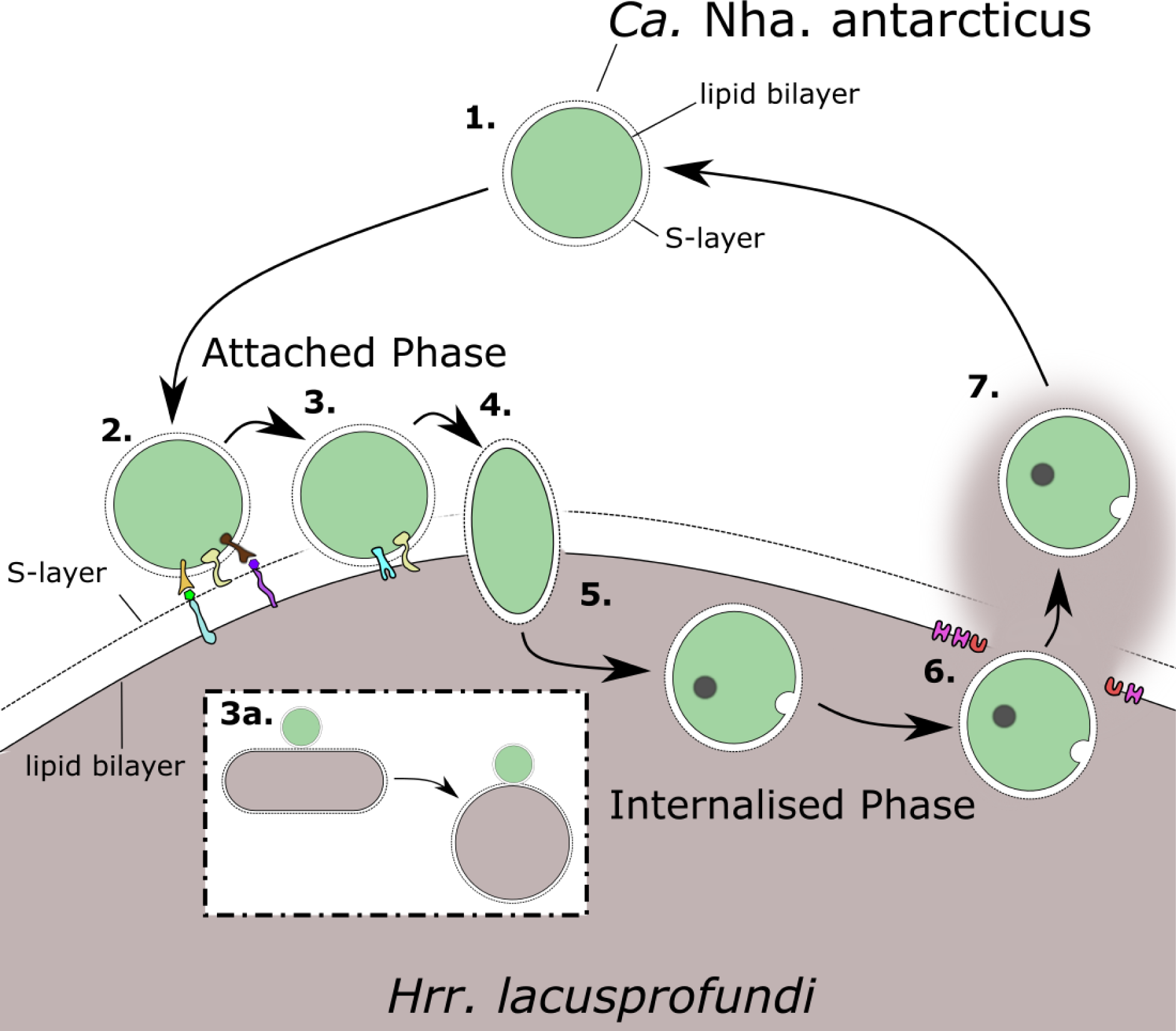
Proposed lifestyle of *Ca.* Nha. antarcticus. 1) Unattached Stage: *Ca.* Nha. antarcticus cells utilise nutrient stores accumulated during interactions with *Hrr. lacusprofundi* to enable survival until a new host cell is encountered and successfully invaded. **2)** Host Recognition Stage: Initial attachment involves host recognition and adhesion potentially mediated by *Ca.* Nha. antarcticus proteins with domains including cell adhesion domains including Ig-folds (e.g. SPEARE protein^1^; yellow), lectin/glucanase domains (brown), and pectin lyase domains (orange), which likely bind to sugar moieties on host glycolipids and glycoproteins. **3)** Pre-Invasion Stage: Localised degradation of the host S-layer by proteases including potentially the protease domain on the SPEARE protein (yellow) enables *Ca.* Nha. antarcticus access to the *Hrr. lacusprofundi* membrane. **3a)** The localised degradation of the host S-layer can destabilise host cell morphology and cause some rod-shaped host cells to become coccoidal. The timing of morphological change varies and is likely dependent on a range of factors such as the site of invasion and size of the cells. **4)** Invasion Stage: The *Ca.* Nha. antarcticus cell passes into the host cell. **5)** Internalised Stage: while internalised, *Ca.* Nha. antarcticus cells acquire nutrients from *Hrr. lacusprofundi* for proliferation and generate stores of nutrients including PHA-like granules (grey) and membrane embedded lipid droplets. **6)** Host Lysis Stage: The *Hrr. lacusprofundi* membrane is destabilised and many host cells lyse. **7)** Release Stage: *Ca.* Nha. antarcticus cells are released into the environment, enabling them to re-enter the Unattached Stage.

### Genetic loci with potential roles in DPANN – Host interactions

Type IV pili are believed to play an important role in the lifestyles of *B. bacteriovorus*^40^, *Ca.* V. lugosii^39^, Saccharibacteria TM7i^38^, and may also facilitate interactions between *Ca.* Nha. antarcticus and *Hrr. lacusprofundi*. Analysis of a set of 569 representative archaeal genomes revealed the presence of conserved loci encoding Type-IV pilus homologues across multiple cluster 2 DPANN^8^ lineages (Nanohaloarchaeota, Woesearchaeota, Pacearchaeota, Nanoarchaeota, and Aenigmarchaeota) as well as Undinarchaeota. Two distinct gene clusters were identified within these lineages which, in addition to the Type-IV pilus genes, encoded proteins with coiled-coil domains predicted to structurally resemble viral attachment proteins (sigma-1 protein: PDB_ID 6GAO, Fig. 5). The association of proteins structurally similar to viral proteins is conspicuous considering the apparent invasion of the host cell observed in microscopy data. However, these loci are also present in a cultivated species belonging to the Nanoarchaeota (Ca. Nanoclepta minutus^13^) that has not been observed to internalise and as such seem unlikely to be directly responsible for internalisation. Previously generated proteomics data^1^ confirmed several proteins within the *Ca.* Nha. antarcticus loci are actively expressed including the coiled-coil domain containing proteins. In addition to the type-IV pilus-like loci, comparisons of the genetic content between *Ca.* Nha. antarcticus and *Ca.* Nanohalobium constans^17^ (a cultivated nanohaloarchaeon that was not reported to invade host cells) revealed that *Ca*. Nha. antarcticus encodes proteins that structurally resemble autolysins, bacteriocins, and phage cell-puncturing proteins (Supplementary Discussion) which are absent from the *Ca*. Nanohalobium constans genome and may support predation of the host.

**Fig. 5.**
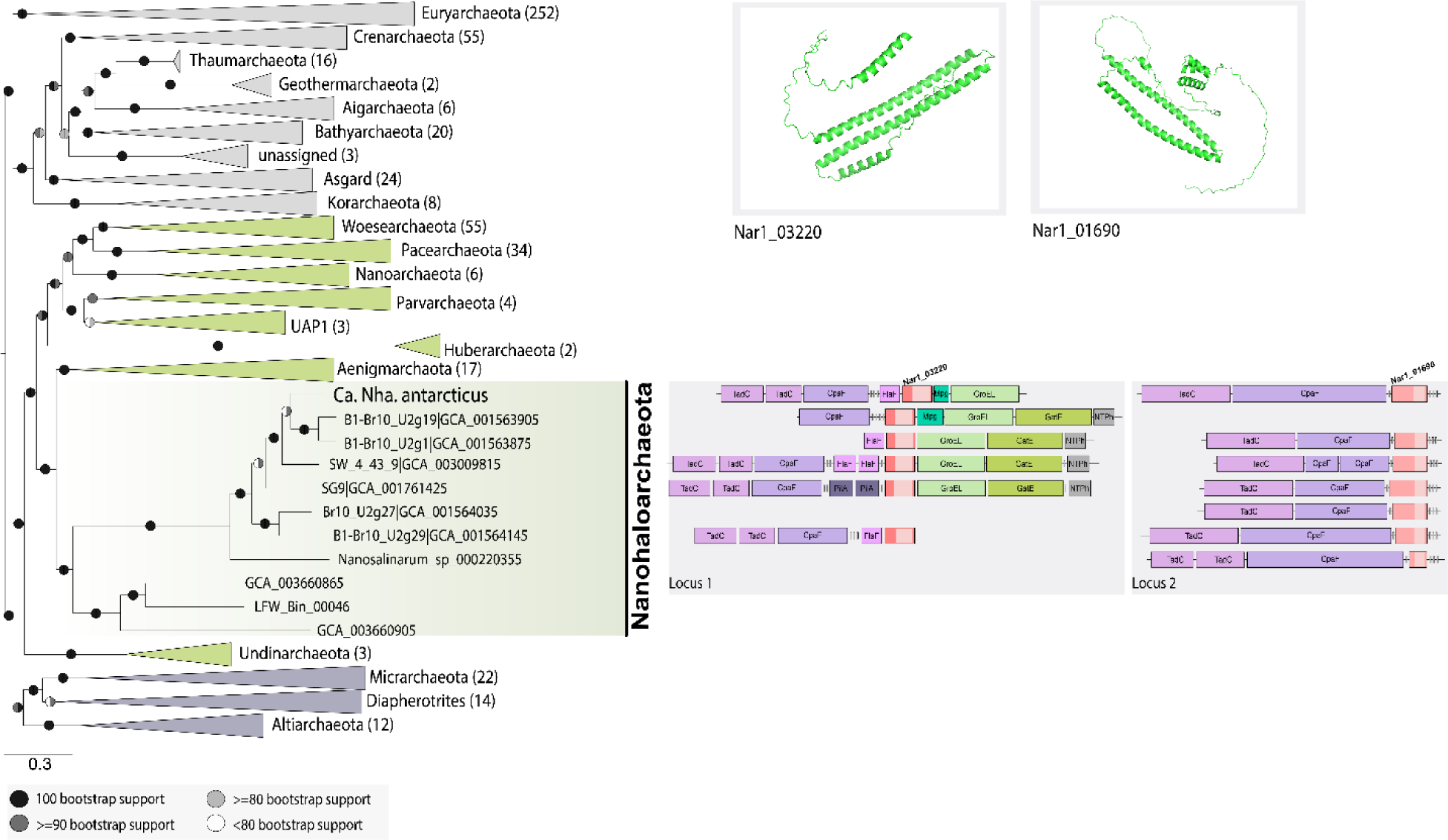
Conservation of loci encoding CCP genes in Nanohaloarchaeota. The maximum-likelihood phylogenetic tree was based on 51 marker proteins and 569 archaeal species. The alignment was trimmed with BMGE^56^ (alignment length, 11399 aa). A phylogenetic tree was inferred in IQ-TREE^76^ with the LG+C20+F+R model with an ultrafast bootstrap approximation (left half of bootstrap symbol) and SH-like approximate likelihood test (right half of bootstrap symbol), each run with 1000 replicates (see key for shading indicating bootstrap support). The tree was artificially rooted between DPANN Archaea (cluster 1 DPANN in dark purple, cluster 2 DPANN in green) and all other Archaea (shaded in grey). The number of species represented in each clade is shown in parentheses after the taxonomic name of the clade. Scale bar: average number of substitutions per site. The two, Cluster 2 DPANN loci are shown aligned to the Nanohaloarchaeota sequences in the phylogenetic tree. The OmegaFold predicted coiled-coil structure of both the *Ca.* Nha. antarcticus locus 2 CCPs (NAR1_03220 and NAR1_01690) is depicted above the locus diagram. *Ca.* Nha. antarcticus proteins identified in proteomic data are highlighted (bold outline). The type-IV filament proteins encoded in each locus (CpaF, pilus assembly ATPase; TadC, membrane assembly platform) or just Locus 1 (FlaF and PilA, filament proteins) are shown. Other proteins encoded in Locus 1 are Mpg (3-methyladenine DNA glycosylase), GroEL (chaperonin), GatE (Archaeal Glu-tRNA (Gln) amidotransferase subunit E) and NTPhyd (P-loop containing nucleoside triphosphate hydrolase). The gene-locus images were manually generated and loci were only included if they had putative flagella or pili genes up- or downstream of the CCP genes.

### Conclusions

Here we describe the lifestyle of a symbiotic DPANN archaeon leading to host cell lysis which appears to involve an intracellular stage. It is unclear how widespread such parasitic lifestyles may be amongst DPANN Archaea as the majority of DPANN species are uncultivated^9^ and the environmental factors that influence growth of DPANN remain enigmatic. However, the characteristics we describe herein illustrate the potential capacity of certain DPANN Archaea to contribute to nutrient cycling through lysis of host cells, similar to viral predation in the top-down control of the food web in Antarctic aquatic systems^43^. In this way, *Ca.* Nha antarcticus is likely to contribute to the recycling of nutrients in the three haloarchaeal-dominated, hypersaline lakes that it is known to colonise^1^. Considering that it has been suggested that DPANN Archaea may associate not only with other archaea but also with bacteria^7,9,44^, the capacity of some representatives of the DPANN Archaea to behave in a predatory manner similar to viruses could have implications for microbial food web dynamics across the globe and may necessitate a re-evaluation of their functional importance and ecological roles.

## Methods

### Production of nanohaloarchaeal cells

Purified *Ca.* Nha. antarcticus cells were sourced from an enrichment culture (Nha-CHl) grown at 18 °C by FACS as previously described^1^ or through filtration. To acquire *Ca.* Nha. antarcticus through filtration 10 mL of the Nha-CHl culture was first filtered three times through a 0.8 µm pore size cellulose acetate syringe filter and then subsequently filtered five times through a 0.22 µm pore size cellulose acetate syringe filter. The resulting filtrate was centrifuged at 20,000 *g* for 10 min and the cell pellet was resuspended in 1 mL of fresh DBCM2^1^. To confirm purity of filtered cells, aliquots were spot plated on DBCM2 agar and incubated for 2 months at 30 °C. Absence of growth indicated filtration had successfully removed *Hrr. lacusprofundi* cells from the sample. *Hrr. lacusprofundi* strain R1S1^35^ cells were grown as previously described for strain ACAM34^1^, and after two weeks growth, incubated with FACS-purified *Ca.* Nha. antarcticus cells.

### Live fluorescence microscopy

MitoTracker dye (1:1000 dilution; 1 μM final centration) was added to 500 μL of *Ca. Nha. antarcticus* sorted cells (∼ 2 x 10^7^ mL^-1^; MitoTracker Deep Red FM) or *Hrr. lacusprofundi* cells (∼ 3 x 10^8^ mL^-1^; MitoTracker Orange CMTMRos)^22^. Cells were maintained at 30°C with static incubation for 1 h. The dye was washed out three times with fresh DBCM2 media^1^ via centrifugation after staining and resuspended in 50 μL (*Ca.* Nha. antarcticus cells) or 250 μL (*Hrr. lacusprofundi*) of DBCM2 media. Resuspended *Hrr. lacusprofundi* (2 μl) and *Ca.* Nha. antarcticus (4 μL) cells were mixed prior to use. For live-cell fluorescent microscopy imaging, 3 μL of mixed cells was placed on a ∼1 mm thick agarose pad (0.3% w/v agarose and containing the full media requirements for DBCM2 media), that had been prepared on a 8 mm diameter #1.5 circular glass coverslip (World Precision Instruments, Inc). The coverslip-pad-cell sample assembly was placed inverted onto the base of a 35 mm #1.5 FluoroDish (WPI)^45^. The pre-warmed (30°C) liquid DBCM2 medium (4 mL) was gently applied to cover the pad assembly in the FluoroDish. The lid was applied to avoid evaporation and the dish was incubated on the microscope stage (at 30°C) for imaging. The initial stages of microscope setup for obtaining images of multiple, individual cells took ∼ 1 h, meaning that cells had the opportunity to interact prior to the initiation (t_0_) of the time course. Time-lapse fluorescence imaging was performed at 30°C on a Nikon Ti-E-Perfect Focus microscope with DS-Qi2 camera and a × 100 Oil Plan NA 1.45 objective using a TRITC filter (Ex: 535/36 nm; Em: 590/34 nm) for the MitoTracker Orange fluorescence signal, and a Cy5 filter (Ex: 645/30 nm; Em: 660/40 nm) for the MitoTracker Deep Red fluorescence signal. Z-stack imaging was performed on a confocal laser scanning Nikon A1 microscope with A1-DUG GaAsP Multi Detector Unit (hybrid 4-channel detector: 2 GaAsP PMTs + 2 normal PMTs) at 30°C using a Plan Fluor 100 × Oil objective (z-axis step 0.125 μm) with the TRITC filter (Ex: GaAsP 561 nm; Em: 595/50 nm) and Alx647 channel (Ex:PMT, 637.4 nm; Em 700/75 nm), or on a DeltaVision Elite microscope at 30°C using a 100 × Oil NA 1.4 objective (Z-axis step 0.2 μm or 0.5 μm) with the TRITC and Cy5 filters. The imaging data were processed for deconvolution and bleach correction as stated in the figure legend. The processed Z-stack data were re-constructed for 3D ‘orthogonal’ slice projection analyses using the Imaris software package (Bitplane AG, Zurich, Switzerland).

To determine whether migration of the *Ca.* Nha. antarcticus stain into *Hrr. lacusprofundi* R1S1 cells corresponded to internalisation or invagination, cells were stained with either MitoTracker Orange (*Hrr. lacusprofundi* R1S1) or MitoTracker Green (*Ca.* Nha. antarcticus) as described above (1 μM final concentration). Cells were then mixed and incubated at 30°C. Samples (10 μL) were taken hourly and additionally stained with Concanavalin A (Alexa Fluor 350 conjugated, 200 μg/mL) and RedDot 2 (200X solution diluted to 1X final concentration). Cells were mounted onto glass slides and imaged on a Carl Zeiss Imager M.2 microscope at room temperature with a 100× Neofluor objective using a Carl Zeiss filter sets 02 (Ex: G 365 nm; Em: LP 420 nm), 38 (Ex: BP 470/40 nm; Em: BP 525/50 nm), 00 (Ex: BP 530-585 nm; Em: LP 615 nm), and 50 (Ex: BP 640/30 nm, Em: BP 690/50 nm). To assess the effects of MitoTracker dye on cell growth, *Hrr. lacusprofundi* R1S1 cells were stained with MitoTracker Orange (1 μM final concentration) as described above. MitoTracker-stained and unstained control cells were each inoculated into 5 mL fresh DBCM2 medium in 50 mL conical tubes (three biological replicates) to an optical density (OD_595_) of ∼0.05, cultures incubated with shaking (150 RPM) at 30 °C, duplicate aliquots dispensed daily into microtitre plates, and OD_595_ monitored using a SpectraMax 190 Microplate Reader (Molecular Devices LLC) with fresh DBCM2 medium as a blank. To assess the effects of MitoTracker dye reversal on the interactions between *Ca.* Nha. antarcticus and *Hrr. lacusprofundi*, FACS-purified *Ca.* Nha. antarcticus cells were stained with MitoTracker Orange CMTMRos (1 μM final centration), *Hrr. lacusprofundi* R1S1 cells were stained with MitoTracker Deep Red FM (1 μM final concentration), and cell mixtures were imaged using fluorescence time-lapse microscopy as described above. Reversing the labelling with dyes yielded analogous results to prior experiments (Fig. S8). To assess the effects of cell fixation on interactions between *Ca.* Nha. antarcticus and *Hrr. lacusprofundi*, FACS-purified *Ca.* Nha. antarcticus cells (500 μL, ∼ 2 x 10^7^ mL^-1^) were pelleted (5 min, 19,745 *g*) and gently resuspended in 1 mL 18% buffered salt water^46^ containing 4 % (v/v) paraformaldehyde (PFA) and cells fixed by shaking (250 RPM) at room temperature overnight. The fixed cells were washed twice by centrifugation (5 min, 19,745 *g*), and the cell pellet resuspended in 500 μL DBCM2 medium. The fixed *Ca.* Nha. antarcticus cells were stained with MitoTracker Deep Red FM and incubated with *Hrr. lacusprofundi* R1S1 cells stained with MitoTracker Orange CMTMRos, and the cells imaged as described above. Pre-treatment of *Ca.* Nha. antarcticus cells with paraformaldehyde led to a reduced number of *Hrr. lacusprofundi* cells with attached *Ca.* Nha. antarcticus cells (106 of 186 imaged *Hrr. lacusprofundi* cells; 57 %) and subsequently fewer lysed *Hrr. lacusprofundi* cells (12 cells; 6.5%) (Fig. S8, Supplementary Table 1).

Pre-treatment of the *Hrr. lacusprofundi* cells with paraformaldehyde also resulted in a substantial reduction in the frequency of both attachment (31 of 265 imaged *Hrr. lacusprofundi* cells; 11.7%) and lysis events (no cells: 0%) (Fig. S8, Supplementary Table 1). Agarose pad time-course experiments were performed by staining *Hrr. lacusprofundi* R1S1 cells with MitoTracker Orange CMTMRos and FACS-purified *Ca.* Nha. antarcticus cells with MitoTracker Deep Red FM, as described above. The mixed cultures were sampled at different time-points (0, 3, 6, 9, 12 and 24 h) and placed on a 1 % (w/v) agarose pad containing DBCM2 basal salts on a glass slide with a #1.5 glass coverslip placed on top, and cells imaged as described above.

Microfluidic time-course interactions between *Ca.* Nha. antarcticus and *Hrr. lacusprofundi* were performed using a CellASIC ONIX2 microfluidics system to immobilize and record live cells that were exposed to a constant flow of liquid. CellASIC B04A plates (EMD Millipore) were equilibrated with 1 mg mL^-1^ Bovine Serum Albumin in phosphate-buffered saline followed by DBCM2 basal salts at a constant flow pressure of 5 psi. The mixed cell culture (*Hrr. lacusprofundi* R1S1 stained with MitoTracker Orange CMTMRos, and FACS-purified *Ca.* Nha. antarcticus cells stained with MitoTracker Deep Red FM) were loaded into the microfluidics chamber and perfused with DBCM2 medium at 0.25 psi for up to 48 h. Cells were imaged at 30 °C every hour or 30 min using a Nikon TiE2 inverted microscope fitted with a 100× oil-immersion phase-contrast NA 1.45 objective.

For display purposes, time-lapse images were prepared by using OMERO and where needed adjusted for enhancing brightness with same setting applied to the whole image series. The quantitative analysis for the attachment, lysis, morphological change events and cell area were performed by combining automated detection (in FIJI 1.52P^47^ and Microbe J 5.13I^48^) and manual curation. Cell outlines were detected in MicrobeJ by phase-contrast image using the Local Default method and manually corrected where needed. Fluorescence signals were detected by ‘Maxima’ in Microbe J using the Foci and Basic modes (*Hrr. lacusprofundi* fluorescence: tolerance 1000, Z score 20, area > 0.5 intensity > 800; *Ca.* Nha. antarcticus fluorescence: tolerance 1000, Z-score 6, area > 0.05, intensity > 200). For quantification of interactions in experiments using MitoTracker Green and Orange, Concanavalin A, and Reddot 2, channels were subjected to auto thresholding (Moments dark stack: MitoTracker Green, MitoTracker Orange, and Reddot 2; MaxEntropy dark stack: Concanavalin A). Channels were then converted to binary masks and particles counted (“Analyze Particles…”, “size=0.1-Infinity summarize in_situ”). Interactions between *Ca.* Nha. antarcticus and *Hrr. lacusprofundi* were quantified by taking overlaps between MitoTracker Green and MitoTracker Orange (ImageCalculator(“AND create”)) and counting particles (“Analyze Particles…”, “summarize in_situ”). Association of *Ca.* Nha. antarcticus with lysis events was quantified by taking overlaps between MitoTracker Green and Reddot 2 (ImageCalculator(“AND create”)) and counting particles (“Analyze Particles…”, “summarize in_situ”).

### Cryo electron microscopy and tomography

For cryo-CLEM, purified *Ca.* Nha. antarcticus and *Hrr. lacusprofundi* cells were stained as described (see **Live fluorescence microscopy**), mixed at a cell-to-cell ratio of 1:3 and incubated for a period of 20 h. Samples were screened using an Olympus BX61 microscope using cellSens v2.2 software (Olympus Corporation) in order to assess staining. Once screened, the sample was loaded onto Quantifoil carbon coated grids (R 2/2; Quantifoil Micro Tools, Jena, Germany) and cryo-fixed by plunge freezing with a Leica GP (Leica Microsystems) into liquid ethane, as previously described^49^. Grids were assembled into autogrids and imaged using a Zeiss LSM 900 upright confocal with Airyscan 2 fluorescence microscope on a Linkam CMS196 Cryo-correlative microscopy stage with a 100x NA 0.75 objective using a TRITC filter at an excitation of 561 nm for MitoTracker Orange and a Cy5 filter at an excitation of 647 nm for MitoTracker DeepRed. Images were acquired using an Axiocam 506 mono camera (Carl Zeiss AG). Autogrids were transferred for imaging on a Talos Arctica Cryo-electron microscope at 200 kV. For cryo-ET of fluorescently labelled and internalised *Ca.* Nha. antarcticus cells, tilt series were acquired at binning 2 on a Falcon 3 camera under the controls of Tomography software (ThermoFisher). For cryo-ET of the enrichment culture, biomass was loaded onto Quantifoil carbon coated grids (Quantifoil Micro Tools, Jena, Germany) and imaged on a Titan Krios (Thermo Fisher Scientific, Waltham, Massachusetts) at 300 kV. Defocus for all images was -10 µm. For cryo-ET of *Hrr. lacusprofundi* – *Ca.* Nha. antarcticus co-cultures cells were mixed as described above for fluorescence microscopy and incubated at 30°C for 17 h. Cells were then loaded onto Quantifoil holey carbon coated grids (Cu/Rh 3.5/1 200 mesh for *Hrr. lacusprofundi* cells and co-cultures and Cu/Rh 2/2 200 mesh for the pure *Ca.* Nha. antarcticus cells). Samples were cryo-fixed by plunge-freezing in liquid ethane using a Vitrobot Mark IV and stored under liquid nitrogen until imaging, as previously described^50^. Imaging was performed on a Titan Krios at 300 kV using a Bioquantum energy filter and the K3 detector (Gatan Inc.). Tilt series was collected at 2° increments between ±60°, defocus was varied from -6 to -12 µm depending on the tilt series (specified in figure legends), and a total dose of 80 e-/Å^2^ was applied over the series. Tomographic reconstructions were performed using IMOD (http://bio3d.colorado.edu/imod/)51 and tomo3D^52^.

### Bioinformatic analyses

For analysing groups of orthologous proteins, a list of all archaeal genomes was downloaded from NCBI. Genomes were filtered on the basis of stage of assembly (scaffold) and number of scaffolds (<100) in order to produce a reduced list of moderate quality genomes for analysis (607 total). Nucleotide and amino acid fasta files were downloaded for those genomes using a custom python script through the NCBI ftp site. Protein sequences from the genomes were run through Orthofinder 2.3.1_53_ using the diamond blast option (-t 16 -S diamond) in order to identify orthologous proteins shared amongst the genomes. Once orthogroups had been identified they were filtered using a custom python script to identify groups unique to DPANN. These orthogroups were then subjected to preliminary domain annotation using InterProScan version 5.25-64.0^54^.

For identification of proteins potentially involved in cell-cell interactions, initial analyses of DPANN-specific protein clusters revealed that many DPANN seemingly encoded one or two proteins with putative nucleopore domains that possessed predicted coiled-coil structures. To investigate these proteins in more detail and determine their distribution across Archaea, HMM profiles were generated from these coiled-coil protein (CCP) amino acid sequences in DPANN and the profiles were used as queries against an archaeal reference database (569 species, Supplementary Table 12). Specifically, the protein sequences from the relevant orthogroups were aligned using MAFFT L-INS-I v7.407^55^ and trimmed using BMGE v1.12 (settings: -t AA -m BLOSUM30 -h 0.55)^56^. Subsequently, protein domains were predicted using HHpred within the hh-suite^57^ with the following two steps: hhblits v3.3.0 was run (settings: -i trimmed_alignment -E 1E-01 -d uniclust30_2018_08) and provided the input a3m file for hhsearch v3.1b2 (settings: -i a3m_file -d pdb70 -p 20 -Z 250 -loc -z 1 - b 1 -B 250 -ssm 2 -sc 1 -seq 1 -dbstrlen 10000 -norealign -maxres 32000 -contxt context_data.crf - blasttab). The top 50 hits were manually inspected for a match to a potential nucleopore domain. The exact positions of the domain in the respective proteins were extracted from the full protein alignment using bedtools v2.26.0^58^ and used to build HMM profiles with hmmbuild. The two profiles were used to search for potential nucleopore domain proteins across our archaeal reference database using a custom script (hmmsearchTable) which implements the hmmsearch algorithm (available in 3_Scripts.tar.gz at https://zenodo.org/record/3839790#.Xywn7btR23U). In order to annotate the identified proteins and verify the presence of nucleopore domains, positive hits were extracted and analysed with HHpred. The potential secondary and tertiary structures of the positive hits were also examined by investigating the protein sequences using the Phyre2 webserver^59^. Additionally, the secondary domain structure of these proteins from *Ca.* Nha. antarcticus was investigated using JPred4^60^. The protein sequences of the ten genes up- and downstream surrounding the CCP genes were examined and all annotations (see below) are provided (Supplementary Table 13), including the top hits of the HHphred and Phyre2 results for the CCPs. To complement Phyre2 predictions OmegaFold^61^ structural predictions were produced for all predicted coding sequences in the *Ca.* Nanohaloarchaeum antarcticus genome. Structurally similar proteins were identified using FoldSeek^62^ and results summarised in Supplementary Table 5.

In order to ensure consistency, all genomes were annotated using the same settings and databases. Gene calling was performed using Prokka^63^ (v1.14, settings: --kingdom Archaea -- addgenes --increment 10 --compliant --centre UU --norrna --notrna). For further functional annotation, the generated protein files were compared against several databases, including the arCOGs (version from 2014)^64^, the KO profiles from the KEGG Automatic Annotation Server (KAAS; downloaded April 2019)^65^, the Pfam database (Release 31.0)^66^, the TIGRFAM database (Release 15.0)^67^, the Carbohydrate-Active enZymes (CAZy) database (downloaded from dbCAN2 in September 2019)^68^, the MEROPs database (Release 12.0)^69^, the Transporter Classification Database (TCDB; downloaded in November 2018)^70^, the hydrogenase database (HydDB; downloaded in November 2018)^71^ and NCBI_nr (downloaded in November 2018). Additionally, all proteins were scanned for protein domains using InterProScan (v5.29-68.0; settings: --iprlookup --goterms)^54^. ArCOGs were assigned using PSI-BLAST v2.7.1+ (settings: -evalue 1e-4 -show_gis -outfmt 6 -max_target_seqs 1000 -dbsize 100000000 -comp_based_stats F -seg no)^72^. KOs as well as PFAMs, TIGRFAMs and CAZymes were identified in all archaeal genomes using hmmsearch v3.1b298 (settings: -E 1e-4)^73^. The Merops database was searched using BLASTp v2.7.1 (settings: -outfmt 6, -evalue 1e-20)^72^. For all database searches, the best hit for each protein was selected based on the highest e-value and bitscore. For InterProScan, multiple hits corresponding to the individual domains of a protein were reported using a custom script (parse_IPRdomains_vs2_GO_2.py).

In order to identify genes that may be involved in the internalisation process, an all-vs-all BLAST (BLASTp v2.7.1 settings: -outfmt 6, -evalue 1e-30)^72^ was performed on the *Ca.* Nha. antarcticus genome against the *Ca.* Nanohalobium genome (Supplementary Table 14). Hits that aligned to <30% of the reference sequence were discarded. *Ca.* Nha antarcticus genes that did not have an identified homolog in *Ca.* Nanohalobium were then subjected to structural prediction through the Phyre2 server (Supplementary Table 4)^59^. Structural predictions were reviewed alongside functional annotations (Supplementary Table 3 - 5) in order to assess likelihood of involvement in the internalisation process.

### Phylogenetic Analyses

Maximum likelihood phylogenetic reconstructions of an archaeal species tree were performed using a combination of the GDTB^74^, ribosomal^7^ and phylosift^75^ marker sets. Briefly, using hmmsearch, a modified TIGRFAM database was queried against a protein database generated from proteins called from 569 archaeal species^8^ (custom scripts are available at https://zenodo.org/record/3839790#.Xywn7btR23U). An initial set of 151 marker protein trees was manually investigated for resolved monophyletic clades of well-established archaeal phylum- or order-level lineages resulting in 51 marker proteins (Supplementary Table 15) used for further analyses. In particular, the 151 single gene trees were generated by individually aligning marker proteins using MAFFT L-INS-i v7.407 (settings: --reorder)^55^, trimming using BMGE v1.12 (settings: -t AA -m BLOSUM30 -h 0.55)^56^ and inferring phylogenetic trees using IQ-TREE (v1.6.7, settings: -m LG+G -wbtl -bb 1000 -bnni)^76^. After selecting the final marker set, the 51 non-redundant marker proteins of interest were extracted from the larger database and individually aligned using MAFFT L-INS-i v7.407 (settings: --reorder)^55^ and trimmed using BMGE v1.12 (settings: -t AA -m BLOSUM30 -h 0.55)^56^. The single proteins were concatenated using catfasta2phyml.pl (https://github.com/nylander/catfasta2phyml) and a phylogenetic tree was generated using IQ-TREE (v1.6.7, settings: -m LG+C20+F+R -bb 1000 -alrt 1000)^76^, visualised using FigTree (v1.4.4) and annotated with Inkscape and Adobe Illustrator.

Single protein trees for relevant pili genes were generated as follows: Pili proteins were identified and extracted based on their arCOG identifiers from the archaeal reference set and a bacterial reference database (3022 species, Supplementary Table 17). arCOGs belonging to the same COG were combined (see Supplementary Table 20). Single gene trees were generated by individually aligning marker proteins using MAFFT L-INS-i v7.407 (settings: --reorder)^55^, trimming using BMGE v1.12 (settings: -t AA -m BLOSUM30 -h 0.55)^56^ and inferring phylogenetic trees using IQ-TREE (v1.6.7, settings: -m LG+C10+F+R -nt 5 -wbtl -bb 1000 -bnni)^76^.

## Acknowledgements

We thank Sarah Brazendale, Alice M. Hancock and expedition personnel for obtaining Antarctic samples. The live fluorescence microscopy results were obtained at UTS Microbial Imaging Facility, and we thank Chris Evenhuis, Michael Johnson, and Louise Cole for technical support. The authors acknowledge the use of the Cryo Electron Microscopy Facility through the Victor Chang Cardiac Research Institute Innovation Centre, funded by the NSW government. Electron microscopy results were obtained at the Electron Microscope Unit, within the Mark Wainwright Analytical Centre of the University of New South Wales and at the Structural Studies Division, MRC Laboratory of Molecular Biology. Cryogenic Light microscopy was performed at the Biomedical Imaging Facility within the Mark Wainwright Analytical Centre of the University of New South Wales. Fluorescence Activated Cell Sorting was performed at the Flow Cytometry Facility within the Mark Wainwright Analytical Centre of the University of New South Wales. We thank Nicholas Ariotti, Joanna Biazik, Hariprasad Venugopal, Gediminas Gervinskas, and Georg Ramm for assistance developing electron microscopy approaches. DNA sequencing of PCR products was performed at the Ramaciotti Centre for Genomics (UNSW Sydney), and computational analyses were performed on the computational cluster Katana, supported by the Faculty of Science (UNSW Sydney). This work was supported by the Australian Research Council (DP150100244 to R. C.; FT160100010 to I. G. D.), the Australian Antarctic Science program (project 4031 to R. C.), the UTS Chancellor’s Postdoctoral Research Fellowship (PRO21-13161 to Y.L), the Swedish Research Council (VR starting grant 2016-03559 to A. S. and supporting J. H.), the NWO-I foundation of the Netherlands Organisation for Scientific Research (WISE fellowship to A. S.), the European Research Council (ERC) under the European Union’s Horizon 2020 research and innovation programme (grant agreement ERC Starting grant No. 947317 to A. S.) as well as by the Moore–Simons Project on the Origin of the Eukaryotic Cell, Simons Foundation 735929LPI, https://doi.org/10.46714/735929LPI. (Award Number: 811944 to A. S. and supporting J. H.). T. A. M. B. would like to thank UKRI MRC (Programme MC_UP_1201/31), the Human Frontier Science Programme (Grant RGY0074/2021), the Vallee Research Foundation, the European Molecular Biology Organization, the Leverhulme Trust and the Lister Institute for Preventative Medicine for support.

## Author Contributions

J. N. H., Y.L., and R. C. conceived the study. J. N. H., Y. L., E. L., C. B., and E. M. V. J. performed cultivation and purification of *Ca.* Nha. antarcticus cells. Y. L., I. G. D., J. N. H, and A. S. conducted live fluorescence microscopy. J. N. H., A. v. K, T. A. M. B, E. L, M. A. B. B., and R. M. W performed electron microscopy. J. N. H, Y. L., A. v. K, T. A. M. B, N. D., E. L., A. S., and R. C. conducted data interpretation. J. N. H, Y. L., A. v. K, T. A. M. B, N. D., E. L., A. S., and R. C. wrote the manuscript with input from all other co-authors. All authors have read and approved the manuscript submission.

## Data and Materials Availability

Supplementary data are available in our repository on figshare [https://doi.org/10.6084/m9.figshare.12957092]

## Supplementary Materials

Extended Data Fig. 1 – 26

Supplementary Tables 1 - 20

Supplementary Videos 1 – 15

## References

1 Hamm, J. N. et al. Unexpected host dependency of Antarctic Nanohaloarchaeota. Proc Natl Acad Sci U S A 116, 14661–14670, (2019).

2 Huber, H. et al. A new phylum of Archaea represented by a nanosized hyperthermophilic symbiont. Nature 417, 63–67, (2002).

3 Liu, X. et al. Insights into the ecology, evolution, and metabolism of the widespread Woesearchaeotal lineages. Microbiome 6, 102, (2018).

4 Baker, B. J. et al. Enigmatic, ultrasmall, uncultivated Archaea. Proc Natl Acad Sci U S A 107, 8806–8811, (2010).

5 Narasingarao, P. et al. De novo metagenomic assembly reveals abundant novel major lineage of Archaea in hypersaline microbial communities. ISME J 6, 81–93, (2012).

6 Koskinen, K. et al. First Insights into the Diverse Human Archaeome: Specific Detection of Archaea in the Gastrointestinal Tract, Lung, and Nose and on Skin. Mbio 8, (2017).

7 Castelle, C. J. & Banfield, J. F. Major New Microbial Groups Expand Diversity and Alter our Understanding of the Tree of Life. Cell 172, 1181–1197, (2018).

8 Dombrowski, N. et al. Undinarchaeota illuminate DPANN phylogeny and the impact of gene transfer on archaeal evolution. Nat Commun 11, 3939, (2020).

9 Dombrowski, N., Lee, J. H., Williams, T. A., Offre, P. & Spang, A. Genomic diversity, lifestyles and evolutionary origins of DPANN archaea. FEMS Microbiol Lett 366, (2019).

10 Tully, B. J., Graham, E. D. & Heidelberg, J. F. The reconstruction of 2,631 draft metagenome-assembled genomes from the global oceans. Sci Data 5, 170203, (2018).

11 Vavourakis, C. D. et al. A metagenomics roadmap to the uncultured genome diversity in hypersaline soda lake sediments. Microbiome 6, 168, (2018).

12 Xie, Y. G. et al. Functional differentiation determines the molecular basis of the symbiotic lifestyle of Ca. Nanohaloarchaeota. Microbiome 10, 172, (2022).

13 St John, E., et al. A new symbiotic nanoarchaeote (Candidatus Nanoclepta minutus) and its host (Zestosphaera tikiterensis gen. nov., sp. nov.) from a New Zealand hot spring. Syst Appl Microbiol 42, 94–106, (2019).

14 Wurch, L. et al. Genomics-informed isolation and characterization of a symbiotic Nanoarchaeota system from a terrestrial geothermal environment. Nat Commun 7, 12115, (2016).

15 Krause, S., Bremges, A., Munch, P. C., McHardy, A. C. & Gescher, J. Characterisation of a stable laboratory co-culture of acidophilic nanoorganisms. Sci Rep 7, 3289, (2017).

16 Golyshina, O. V. et al. ’ARMAN’ archaea depend on association with euryarchaeal host in culture and in situ. Nat Commun 8, 60, (2017).

17 La Cono, V. et al. Symbiosis between nanohaloarchaeon and haloarchaeon is based on utilization of different polysaccharides. Proc Natl Acad Sci U S A 117, 20223–20234, (2020).

18 Reva, O. et al. Functional diversity of nanohaloarchaea within xylan-degrading consortia. Frontiers in Microbiology 14, (2023).

19 Heimerl, T. et al. A Complex Endomembrane System in the Archaeon Ignicoccus hospitalis Tapped by Nanoarchaeum equitans. Front Microbiol 8, 1072, (2017).

20 Comolli, L. R. & Banfield, J. F. Inter-species interconnections in acid mine drainage microbial communities. Front Microbiol 5, 367, (2014).

21 Krause, S. et al. The importance of biofilm formation for cultivation of a Micrarchaeon and its interactions with its Thermoplasmatales host. Nat Commun 13, 1735, (2022).

22 Maslov, I. et al. Efficient non-cytotoxic fluorescent staining of halophiles. Sci Rep 8, 2549, (2018).

23 Tschitschko, B. et al. Genomic variation and biogeography of Antarctic haloarchaea. Microbiome 6, (2018).

24 Franzmann, P. D. et al. Halobacterium lacusprofundi sp. nov., a Halophilic Bacterium Isolated from Deep Lake, Antarctica. Systematic and Applied Microbiology 11, 20–27, (1988).

25 Liao, Y. et al. Morphological and proteomic analysis of biofilms from the Antarctic archaeon, Halorubrum lacusprofundi. Sci Rep 6, 37454, (2016).

26 Williams, T. J. et al. Cold adaptation of the Antarctic haloarchaea Halohasta litchfieldiae and Halorubrum lacusprofundi. Environ Microbiol 19, 2210–2227, (2017).

27 Murphy, D. J. The dynamic roles of intracellular lipid droplets: from archaea to mammals. Protoplasma 249, 541–585, (2012).

28 von Kugelgen, A., Alva, V. & Bharat, T. A. M. Complete atomic structure of a native archaeal cell surface. Cell Rep 37, 110052, (2021).

29 Bharat, T. A. M., von Kugelgen, A. & Alva, V. Molecular Logic of Prokaryotic Surface Layer Structures. Trends Microbiol 29, 405–415, (2021).

30 Herdman, M. et al. High-resolution mapping of metal ions reveals principles of surface layer assembly in Caulobacter crescentus cells. Structure 30, 215–228 e215, (2022).

31 von Kugelgen, A. et al. In Situ Structure of an Intact Lipopolysaccharide-Bound Bacterial Surface Layer. Cell 180, 348–358 e315, (2020).

32 Strubbe-Rivera, J. O. et al. The mitochondrial permeability transition phenomenon elucidated by cryo-EM reveals the genuine impact of calcium overload on mitochondrial structure and function. Sci Rep 11, 1037, (2021).

33 Pfeifer, F. Haloarchaea and the formation of gas vesicles. Life (Basel) 5, 385–402, (2015).

34 Williams, T. J. et al. Microbial ecology of an Antarctic hypersaline lake: genomic assessment of ecophysiology among dominant haloarchaea. ISME J 8, 1645–1658, (2014).

35 Erdmann, S., Tschitschko, B., Zhong, L., Raftery, M. J. & Cavicchioli, R. A plasmid from an Antarctic haloarchaeon uses specialized membrane vesicles to disseminate and infect plasmid-free cells. Nat Microbiol 2, 1446–1455, (2017).

36 Liu, J. et al. Extracellular membrane vesicles and nanotubes in Archaea. microLife 2, (2021).

37 National Research Council Steering Group for the Workshop on Size Limits of Very Small Microorganisms. Size Limits of Very Small Microorganisms: Proceedings of a Workshop. (National Academies Press (US) 1999).

38 Xie, B. et al. Type IV pili trigger episymbiotic association of Saccharibacteria with its bacterial host. Proc Natl Acad Sci U S A 119, e2215990119, (2022).

39 Moreira, D., Zivanovic, Y., Lopez-Archilla, A. I., Iniesto, M. & Lopez-Garcia, P. Reductive evolution and unique predatory mode in the CPR bacterium Vampirococcus lugosii. Nat Commun 12, 2454, (2021).

40 Chanyi, R. M. & Koval, S. F. Role of type IV pili in predation by Bdellovibrio bacteriovorus. PLoS One 9, e113404, (2014).

41 Kellner, S. et al. Genome size evolution in the Archaea. Emerg Top Life Sci 2, 595–605, (2018).

42 Lind, A. E. et al. Genomes of two archaeal endosymbionts show convergent adaptations to an intracellular lifestyle. ISME J 12, 2655–2667, (2018).

43 Cavicchioli, R. Microbial ecology of Antarctic aquatic systems. Nat Rev Microbiol 13, 691–706, (2015).

44 Reysenbach, A. L. et al. Complex subsurface hydrothermal fluid mixing at a submarine arc volcano supports distinct and highly diverse microbial communities. Proc Natl Acad Sci U S A 117, 32627–32638, (2020).

45 Duggin, I. G. et al. CetZ tubulin-like proteins control archaeal cell shape. Nature 519, 362–365, (2015).

46 Dyall-Smith, M. The Halohandbook v7.3. (2015).

47 Schindelin, J., et al. Fiji: an open-source platform for biological-image analysis. Nat Methods 9, 676-682, (2012).

48 Ducret, A., Quardokus, E. M. & Brun, Y. V. MicrobeJ, a tool for high throughput bacterial cell detection and quantitative analysis. Nat Microbiol 1, 16077, (2016).

49 Dubochet, J. & McDowall, A. W. Vitrification of pure water for electron microscopy. Journal of Microscopy 124, (1981).

50 Sulkowski, N. I., Hardy, G. G., Brun, Y. V. & Bharat, T. A. M. A Multiprotein Complex Anchors Adhesive Holdfast at the Outer Membrane of Caulobacter crescentus. Journal of Bacteriology 201, e00112–00119, (2019).

51 Mastronarde, D. N. & Held, S. R. Automated tilt series alignment and tomographic reconstruction in IMOD. J Struct Biol 197, 102–113, (2017).

52 Agulleiro, J. I. & Fernandez, J. J. Tomo3D 2.0--exploitation of advanced vector extensions (AVX) for 3D reconstruction. J Struct Biol 189, 147–152, (2015).

53 Emms, D. M. & Kelly, S. OrthoFinder: solving fundamental biases in whole genome comparisons dramatically improves orthogroup inference accuracy. Genome Biol 16, 157, (2015).

54 Jones, P. et al. InterProScan 5: genome-scale protein function classification. Bioinformatics 30, 1236–1240, (2014).

55 Katoh, K. & Standley, D. M. MAFFT multiple sequence alignment software version 7: improvements in performance and usability. Mol Biol Evol 30, 772–780, (2013).

56 Criscuolo, A. & Gribaldo, S. BMGE (Block Mapping and Gathering with Entropy): a new software for selection of phylogenetic informative regions from multiple sequence alignments. BMC Evol Biol 10, 210, (2010).

57 Steinegger, M. et al. HH-suite3 for fast remote homology detection and deep protein annotation. BMC Bioinformatics 20, 473, (2019).

58 Quinlan, A. R. & Hall, I. M. BEDTools: a flexible suite of utilities for comparing genomic features. Bioinformatics 26, 841–842, (2010).

59 Kelley, L. A., Mezulis, S., Yates, C. M., Wass, M. N. & Sternberg, M. J. The Phyre2 web portal for protein modeling, prediction and analysis. Nat Protoc 10, 845–858, (2015).

60 Drozdetskiy, A., Cole, C., Procter, J. & Barton, G. J. JPred4: a protein secondary structure prediction server. Nucleic Acids Res 43, W389–394, (2015).

61 Wu, R., et al. High-resolution de novo structure prediction from primary sequence. bioRxiv, 2022.2007.2021.500999, (2022).

62 van Kempen, M. et al. Fast and accurate protein structure search with Foldseek. Nat Biotechnol, (2023).

63 Seemann, T. Prokka: rapid prokaryotic genome annotation. Bioinformatics 30, 2068–2069, (2014).

64 Makarova, K. S., Wolf, Y. I. & Koonin, E. V. Archaeal Clusters of Orthologous Genes (arCOGs): An Update and Application for Analysis of Shared Features between Thermococcales, Methanococcales, and Methanobacteriales. Life (Basel) 5, 818–840, (2015).

65 Aramaki, T. et al. KofamKOALA: KEGG Ortholog assignment based on profile HMM and adaptive score threshold. Bioinformatics 36, 2251–2252, (2020).

66 Bateman, A. et al. The Pfam protein families database. Nucleic Acids Res 32, D138–141, (2004).

67 Haft, D. H., Selengut, J. D. & White, O. The TIGRFAMs database of protein families. Nucleic Acids Res 31, 371–373, (2003).

68 Yin, Y. et al. dbCAN: a web resource for automated carbohydrate-active enzyme annotation. Nucleic Acids Res 40, W445–451, (2012).

69 Rawlings, N. D. Peptidase specificity from the substrate cleavage collection in the MEROPS database and a tool to measure cleavage site conservation. Biochimie 122, 5–30, (2016).

70 Saier, M. H., Jr., Tran, C. V. & Barabote, R. D. TCDB: the Transporter Classification Database for membrane transport protein analyses and information. Nucleic Acids Res 34, D181–186, (2006).

71 Sondergaard, D., Pedersen, C. N. & Greening, C. HydDB: A web tool for hydrogenase classification and analysis. Sci Rep 6, 34212, (2016).

72 Altschul, S. F. et al. Gapped BLAST and PSI-BLAST: a new generation of protein database search programs. Nucleic Acids Res 25, 3389–3402, (1997).

73 Finn, R. D., Clements, J. & Eddy, S. R. HMMER web server: interactive sequence similarity searching. Nucleic Acids Res 39, W29–37, (2011).

74 Chaumeil, P. A., Mussig, A. J., Hugenholtz, P. & Parks, D. H. GTDB-Tk: a toolkit to classify genomes with the Genome Taxonomy Database. Bioinformatics 36, 1925–1927, (2019).

75 Darling, A. E. et al. PhyloSift: phylogenetic analysis of genomes and metagenomes. PeerJ 2, e243, (2014).

76 Nguyen, L. T., Schmidt, H. A., von Haeseler, A. & Minh, B. Q. IQ-TREE: a fast and effective stochastic algorithm for estimating maximum-likelihood phylogenies. Mol Biol Evol 32, 268–274, (2015).

