## Supplementary Information for "The parasitic lifestyle of an archaeal symbiont"

### Supplementary Discussion

#### *Ca.* Nha. antarcticus compared to *Ca.* Nanohalobium

##### Genes unique to *Ca.* Nha. antarcticus

The observed lifestyle of *Ca.* Nha. antarcticus contrasts with the reported lifestyle of *Ca.* Nanohalobium. In order to investigate the differences between the two Nanohaloarchaeota an all-vs-all BLAST was carried out and *Ca.* Nha. antarcticus proteins with no identifiable homolog in the *Ca.* Nanohalobium metagenome assembled genome MAG extracted (438 genes, Supplementary Table 14). Functional annotations of these proteins were reviewed, and structural predictions carried out using Phyre2[1] to assess the potential role of these proteins in the internalisation lifecycle of *Ca.* Nha. antarcticus (Supplementary Table 4). Complete structures for all *Ca.* Nha. antarcticus proteins were also predicted with OmegaFold[2] and a structural similarity search conducted using FoldSeek[3]. For many of these candidates structural homology to known proteins was not identified (Supplementary Table 5). Two proteins (Halo_Nar1_07490 and Halo_Nar1_00600) were predicted by both Phyre2 and Omegafold to structurally resemble a n-acetylmuramoyl-l-alanine amidase, an autolysin involved in cell wall degradation[4]. Six proteases, including one predicted by Phyre2 to structurally resemble karilysin protease (Halo_Nar1_05310) associated with pathogenicity of the bacterium, *Tannerella forsythia*[5], which OmegaFold also predicted to be structurally similar to proteases, were also identified within the *Ca.* Nha. antarcticus MAG but was absent from the *Ca.* Nanohalobium MAG (Supplementary Tables 4 and 5). Two additional proteins (Halo_Nar1_00290 and Halo_Nar1_01640) were predicted by Phyre2 to structurally resemble the colicin Ia bacteriocin, which has been found to be involved in the killing of host cells by forming a voltage-dependent channel in the target cell's membrane[6] but structural homologues of the OmegaFold structure were not identified through FoldSeek. This was also true for a protein predicted to resemble envelope glycoproteins involved in fusion of viral membranes and host cells (Halo_Nar1_11390) by Phyre2. The majority of proteins (234) were not annotated with any functional domains and a further 66 were annotated with domains of unknown function or general function prediction only (Supplementary Table 3).

#### Orthogroup Analysis

##### Orthogroups potentially involved in DPANN-host interactions

Whilst the above-mentioned proteins may be involved in the observed behaviour of *Ca.* Nha. antarcticus, we cannot exclude the possibility that proteins shared between *Ca.* Nanohalobium and *Ca.* Nha. antarcticus mediate interactions in *Ca.* N. constans - *Halomicrobium* but fulfil different functions or are not actively synthesized under the conditions that have been used for cultivating *Ca.* Nanohalobium. To address this, we performed orthogroup analysis of 607 archaeal genomes to identify DPANN specific protein families that may be involved in mediating interactions. Functional predictions of orthogroups unique to DPANN *Archaea* revealed that many possessed functional domains which may facilitate DPANN-host interactions. These include immunoglobulin-folds (166 protein families), concanavalins (18 protein families), pectin lyases (22 protein families), fibronectins (13 protein families), β-propellers (6 protein families), von Willebrand factors (5 protein families), and polycystic kidney disease domains (7 protein families) (Supplementary Table 6 and 7). Included in this list were several protein families that initially appeared to encode domains homologous to eukaryotic nucleoporin and golgin proteins (Supplementary Tables 8 – 11). Further analysis indicated that these proteins are not related to eukaryotic proteins but share similar structural features (see **Gene context and potential function of CCPs**). Typically, these protein families contained coiled-coil domains, commonly found in proteins involved in tethering and attachment and may therefore be involved in interactions between DPANN and their hosts.

#### Coiled-coil proteins (CCPs)

##### Distribution of genes encoding CCPs

The identification of CCP genes resembling nucleoporin and golgin genes among the DPANN orthogroups prompted us to analyse their distribution and function across Archaea. Particularly, to determine whether the CCP orthologous groups (OGs) discovered in *Ca.* Nha. antarcticus are indeed unique to DPANN, their distribution was analysed across a reference set of 569 archaeal species using sensitive homology searches (see Methods for details) as well as by investigating their genomic location (see **Gene context and potential function of CCPs**). The two Hidden Markov models (HMM)-profiles built for each OG alignment recovered 96 and 186 putative homologs across *Archaea* encoded within two distinct gene clusters (Locus 1 and Locus 2, Fig. 4, Extended Data Fig. 18, Supplementary Tables 13 and 18). Of these, the majority of OG alignments (71 and 112) were found in DPANN *Archaea*. While homologs were not detected in DPANN cluster 1 lineages[7] (Altiarchaeota, Micrarchaeota and Diapherotrites), many members of the Nanoarchaeota, Nanohaloarchaeota, Parvarchaeota, Pacearchaeota and Woesearchaeota as well as Undinarchaeota (DPANN cluster 2 archaea[7]) contained Locus 1 and Locus 2 gene clusters.

**Gene context and potential function of CCPs**

To obtain insight into the potential function of the CCPs, the homology and structural prediction tool HHpred was used. This revealed that most Locus 1 and Locus 2 CCPs in DPANN cluster 2 *Archaea* had a protein domain which modelled to the capsid protein sigma-1 (PDB_ID 6GAO), a protein involved in viral attachment[8]. The predicted secondary structure of this specific domain (helical coiled-coil) (Extended Data Fig. 18 b,c) is similar to sigma-1 proteins[9]. This was consistent with Phyre2 results of the *Ca*. Nha. antarcticus Locus 2 CCP which modelled over 41% of its length with a confidence above 90% to a sigma-1 coiled-coil tail protein (PDB_ID 6GAJ, Fig. 5, Supplementary Tables 13 and 19). However, the *Ca.* Nha. antarcticus Locus 1 CCP was only distantly related to any known structure based on Phyre2. FoldSeek homology searches using the OmegaFold predictions of the *Ca.* Nha. antarcticus CCP genes did not yield identifiable structural homologues.

To characterize the gene context of Locus 1 and Locus 2, the ten genes up- and downstream of the CCP genes were examined across all *Archaea* that encoded at least one of the CCP genes. While all DPANN cluster 2 *Archaea* exhibited a conserved gene arrangement around the CCP genes, minimal conservation was observed in other archaeal lineages (Extended Data Fig. 18 a; Supplementary Table 13). In particular, the DPANN Locus 1 CCP gene was flanked by genes annotated as archaeal chaperonin (GroEL; arCOG01257) on one side, and pili/flagella-related genes on the other side; the latter were annotated as flagella protein FlaF (arCOG01824), the pilus assembly ATPase CpaF (arCOG01819) and the type IV pilus component TadC (arCOG01811). The Locus 2 CCP gene was generally flanked by genes encoding hypothetical proteins on one side, and CpaF and TadC on the other side. Previous work has revealed that various DPANN *Archaea* such as Nanohaloarchaeota[10], Parvarchaeota[11-13] and potentially Micrarchaeota[14] may encode pili, yet the pili gene clusters of DPANN have not been characterized in detail and seem to differ from classified archaeal pili gene clusters[15].

To further assess the diversity of putative pili-like gene clusters in DPANN, searches were performed for the presence of additional pili-related sequences (Supplementary Tables 17 and 20). Most of the proteins constituting Clade 1 pili systems[15] were identified. The secretion peptidase CpaF (arCOG01819) and TadC-like protein (arCOG01811) encoded in Locus 1 and Locus 2 are components of this system. Other Clade 1 pili sequences identified were the regulatory ATPase FleN (arCOG00589), a potential prepilin signal peptidase (arCOG02300), and several minor pilin proteins (arCOG03821, arCOG05787, arCOG05788); the sequences for these minor pilin proteins were distributed across the different DPANN phyla, with *Ca.* Nha. antarcticus possessing homologs assigned to arCOG03821. While InterProScan did not identify any major pili proteins with a class III signal peptide motif in *Ca.* Nha. antarcticus, the genome was shown to encode several proteins with pili domains (IPR013373) that could represent uncharacterized major pili. In fact, all of those pili proteins have a predicted archaeal class III signal peptide cleavage site based on FlaFind[16]. However, *Ca.* Nha. antarcticus seems to lack PilY-like adhesins, which are generally part of Clade 1 pili. It is possible that sigma-capsid-like proteins in *Ca.* Nha. antarcticus represent comparable adhesins, and that at least one of these DPANN cluster 2 *Archaea* gene clusters (Locus 1 and Locus 2) may mediate cell-cell interactions including tethering of Nanohaloarchaeota to their host.

To investigate the evolutionary history of pili proteins encoded by *Ca.* Nha. antarcticus, pilus genes were extracted from a set of reference archaeal and bacterial genomes (Supplementary Table 20) based on arCOG/COG identifiers (Supplementary Table 17). The topologies of the single gene trees, that often included distinct paralogous sub-families, indicate various instances of horizontal gene transfers, but the resolution and support for certain cladings are general low (Extended Data Fig. 18 - 26) such that it is impossible to draw definite conclusions about the history of the various pili gene families across archaea. However, it seems that putative HGT events are not necessarily consistent across multiple single gene trees suggesting that transfers may affect individual genes, rather than entire gene clusters.

#### Supplementary References


1 Kelley, L. A. *et al.* The Phyre2 web portal for protein modeling, prediction and analysis. *Nat Protoc* **10**, 845-858,(2015).

2 Wu, R. *et al.* High-resolution de novo structure prediction from primary sequence. *bioRxiv*, 2022.2007.2021.500999,(2022).

3 van Kempen, M. *et al.* Fast and accurate protein structure search with Foldseek. *Nat Biotechnol*,(2023).

4 Scheurwater, E. M. *et al.* Production and purification of the bacterial autolysin N-acetylmuramoyl-L-alanine amidase B from Pseudomonas aeruginosa. *Protein Expr Purif* **56**, 128-137,(2007).

5 Karim, A. Y. *et al.* A novel matrix metalloprotease-like enzyme (karilysin) of the periodontal pathogen Tannerella forsythia ATCC 43037. *Biol Chem* **391**, 105-117,(2010).

6 Cascales, E. *et al.* Colicin biology. *Microbiol Mol Biol Rev* **71**, 158-229,(2007).

7 Dombrowski, N. *et al.* Undinarchaeota illuminate DPANN phylogeny and the impact of gene transfer on archaeal evolution. *Nat Commun* **11**, 3939,(2020).

8 Lee, P. W. *et al.* Protein sigma 1 is the reovirus cell attachment protein. *Virology* **108**, 156-163,(1981).

9 Dietrich, M. H. *et al.* Structural and Functional Features of the Reovirus sigma1 Tail. *J Virol* **92**,(2018).

10 Hamm, J. N. *et al.* Unexpected host dependency of Antarctic Nanohaloarchaeota. *Proc Natl Acad Sci U S A* **116**, 14661-14670,(2019).

11 Baker, B. J. *et al.* Enigmatic, ultrasmall, uncultivated Archaea. *Proc Natl Acad Sci U S A* **107**, 8806-8811,(2010).

12 Comolli, L. R. *et al.* Inter-species interconnections in acid mine drainage microbial communities. *Front Microbiol* **5**, 367,(2014).

13 Comolli, L. R. *et al.* Three-dimensional analysis of the structure and ecology of a novel, ultra-small archaeon. *ISME J* **3**, 159-167,(2009).

14 Golyshina, O. V. *et al.* 'ARMAN' archaea depend on association with euryarchaeal host in culture and in situ. *Nat Commun* **8**, 60,(2017).

15 Makarova, K. S. *et al.* Diversity and Evolution of Type IV pili Systems in Archaea. *Front Microbiol* **7**, 667,(2016).

16 Szabo, Z. *et al.* Identification of diverse archaeal proteins with class III signal peptides cleaved by distinct archaeal prepilin peptidases. *J Bacteriol* **189**, 772-778,(2007).


#### Extended Data Figures

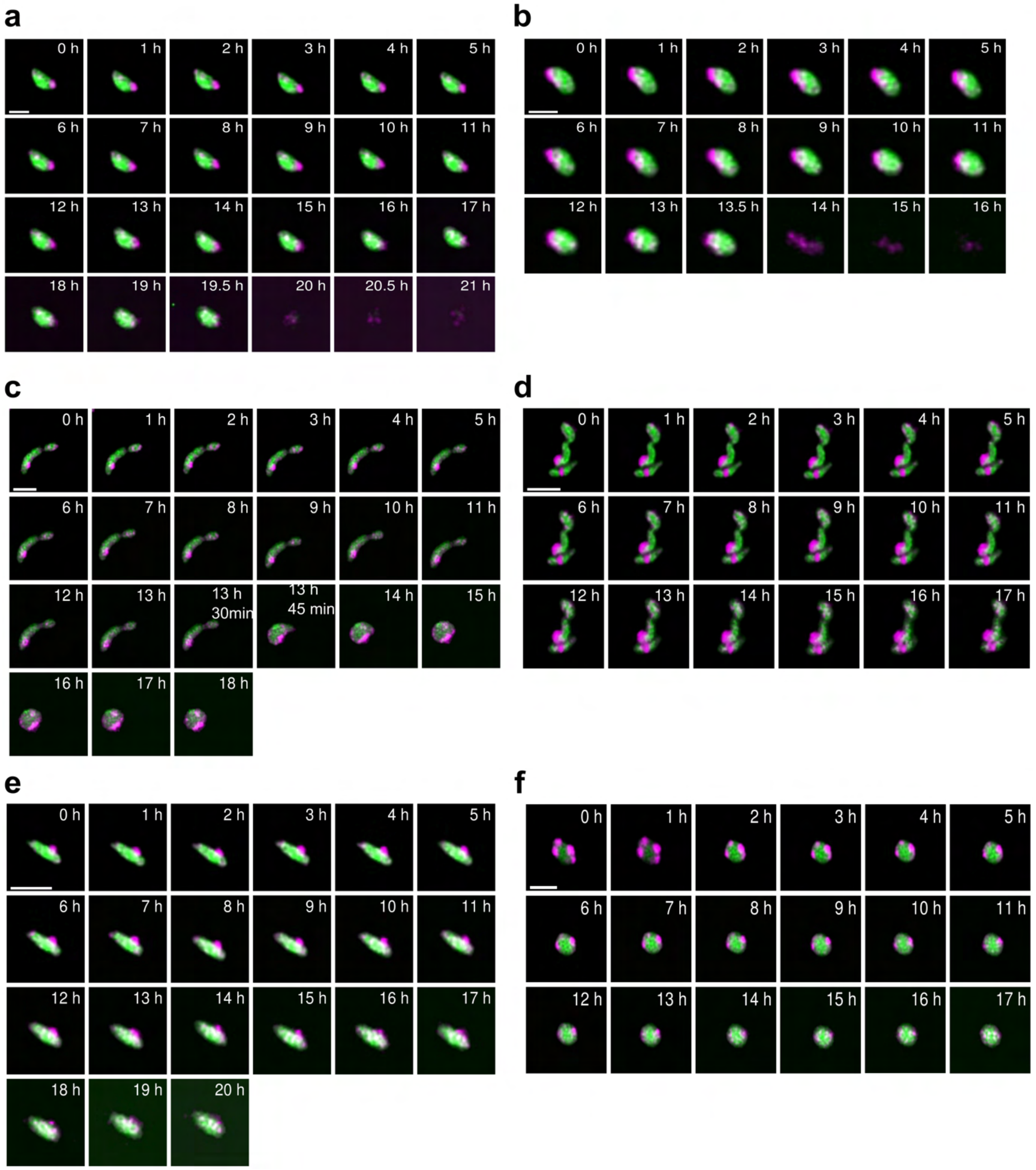


**Extended Data Fig. 1. Live fluorescence microscopy showing internalisation of *Ca.* Nha. antarcticus, lysis and morphological changes of *Hrr. lacusprofundi.*** Live fluorescence time lapse series of *Ca.* Nha. antarcticus (stained with MitoTracker DeepRed, represented magenta) and *Hrr. lacusprofundi* (stained with MitoTracker Orange, represented Green) grown on an agarose pad. **a** & **b,** During the process of *Ca.* Nha. antarcticus migration and internalisation, the short rod-shaped *Hrr. lacusprofundi* cell became rounder (**a**: 0 – 19.5 h; **b**: 0-13.5 h) before lysing (identified through loss of host signal) resulting in the retention of the internalised nanohaloarchaeal cells (**a**: 20 h; **b**: 14 h). The time taken for internalisation and lysis showed variation between different observed cells (also see Fig. 1a). **c**, The host cell abruptly transited (within a 15 min window) from a rod (0 – 13 h 30 min) to a coccus (13 h 45 min – 18 h) morphology. **d**, *Ca.* Nha. antarcticus cells appeared attached to the surface of *Hrr. lacusprofundi* cells and internalised within *Hrr. lacusprofundi* cells. Across the time-course, the fluorescent regions increased in area, consistent with *Ca*. Nha. antarcticus cells replicating. The *Hrr. lacusprofundi* cells retained their morphology, indicating that, unlike panel **a-c**, The process of *Ca.* Nha. antarcticus internalising and replicating did not destabilise *Hrr. lacusprofundi* cellular morphology. **e**, The rod morphology of the *Hrr. lacusprofundi* cell gradually broadened. **f**, Gradual internalisation of the nanohaloarchaeal signal (2 – 10 h), with subsequent diffusion of the signal throughout the host cell (10 – 17 h). The images are indicative of multiple *Ca*. Nha. antarcticus cells simultaneously attaching and internalising within the one *Hrr. lacusprofundi* cell. Scale bars: **a-b**, 1 µm; **c-f**, 2 µm. Images shown were captured on a confocal laser scanning Nikon A1 microscope with “automatic” 2D deconvolution algorithms and adjusted for bleaching using histogram normalisation in Fiji.

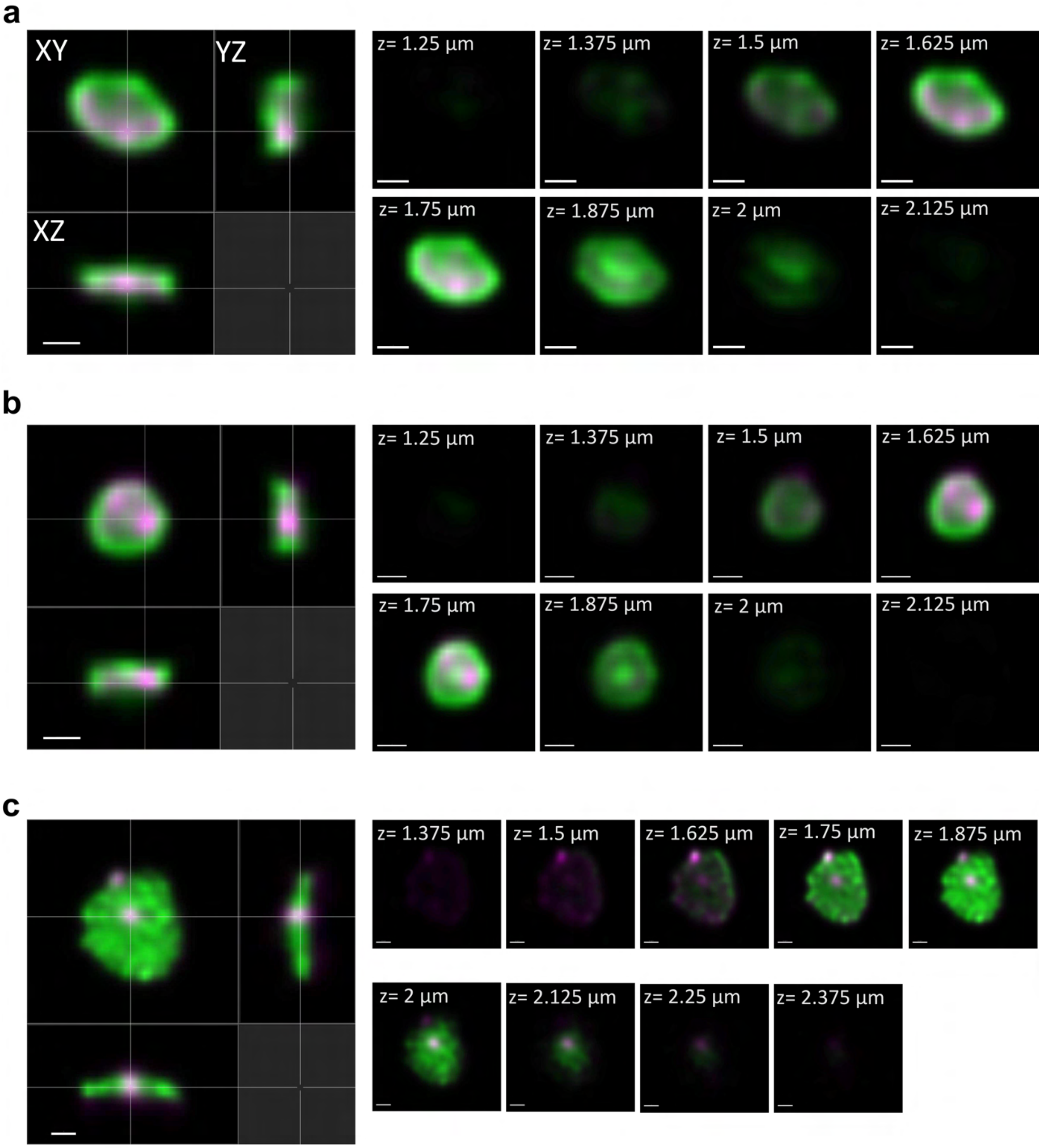


**Extended Data Fig. 2. Confocal microscopy showing internalisation of *Ca*. Nha. antarcticus. a-c**, Mixed culture cells were incubated for 10 h (stained as Extended Data Fig. 1) and then loaded on an agarose pad for imaging. **a-b**, 3D-orthogonal slice analysis (maximum intensity projection, MIP) of *Hrr. lacusprofundi* cells (green) with *Ca*. Nha. antarcticus cells (magenta) internalised. The Z-sectional images (collected at 0.125 µm intervals) showed that by increasing the stack height, the *Ca*. Nha. antarcticus signals started to appear with host cell fluorescence and then disappeared with host cell fluorescence remaining. **c**, A *Hrr. lacusprofundi* cell with *Ca.* Nha. antarcticus cells attached to the surface or located within the host cell profile. Scale bars: **a-c**, 500 nm. Images shown were captured on a confocal laser scanning Nikon A1 microscope with “automatic” 3D deconvolution algorithms.

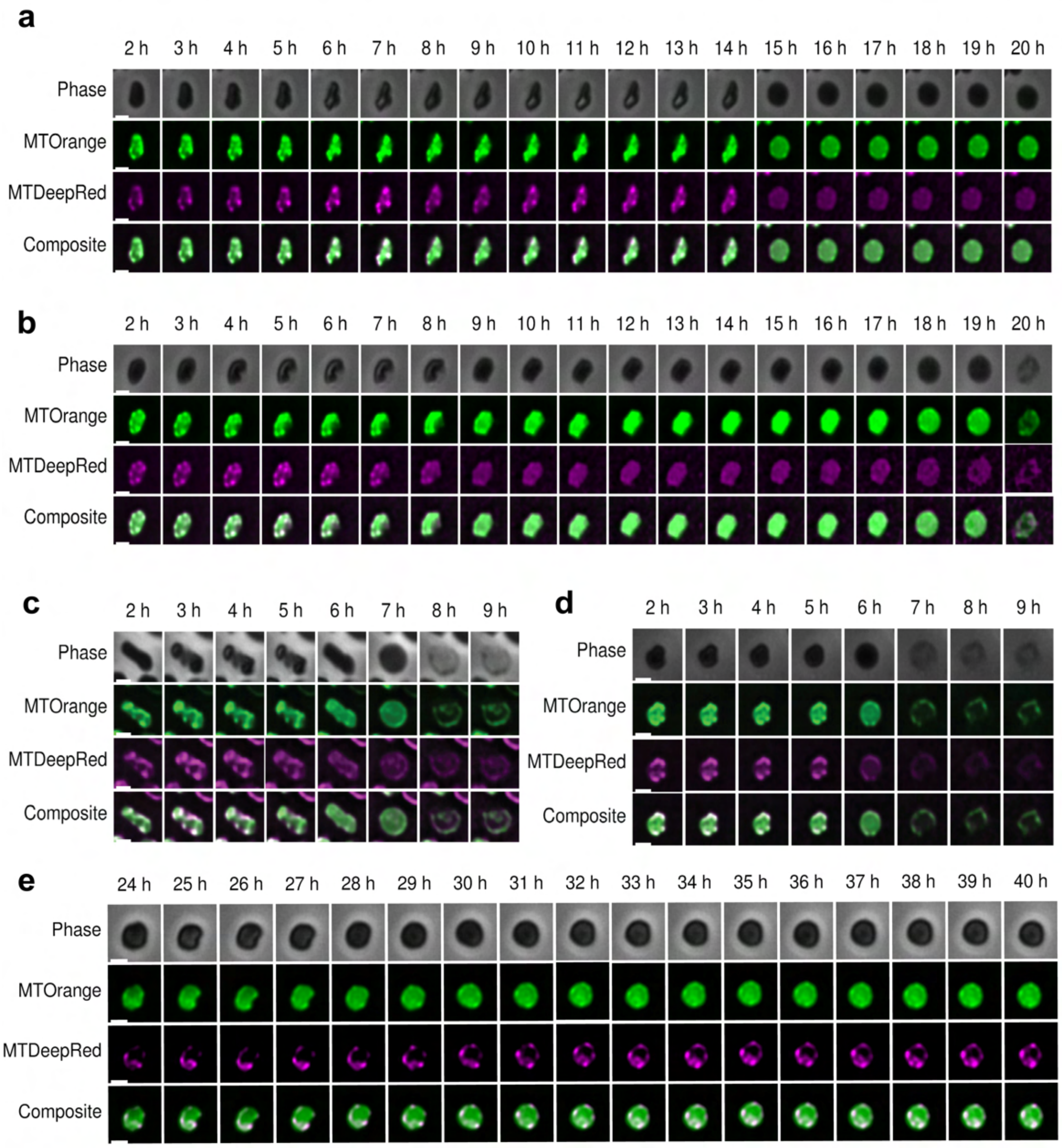


**Extended Data Fig. 3. Live fluorescence microscopy of *Ca*. Nha. antarcticus and *Hrr. lacusprofundi* within a cellASIC microfluidics system.** Cells were stained as Extended Data Fig. 1 (MTOrange: MitoTracker Orange; MTDeepRed: MitoTracker DeepRed). **a - e**, Interactions between *Ca.* Nha. antarcticus and *Hrr. lacusprofundi* within the cellASIC microfluidics system were consistent with those observed in agarose pad live fluorescence experiments. The *Ca*. Nha. antarcticus stain appeared to disperse throughout the host cell generating locally more intense areas, before decreasing in intensity (**a** & **b**) and host cell lysis occurring (**b**). **a - c**, Internalisation of *Ca*. Nha. antarcticus cells within rod shaped *Hrr. lacusprofundi* cells was associated with a shift to a more coccoidal morphology in the *Hrr. lacusprofundi* cell, consistent with observations from agarose pad live fluorescence experiments (Fig. 1a and b, S1, S2). **b - d**, Internalisation frequently led to lysis of the host cell associated with loss of host fluorescent signal. **e**, In some instances interactions were not observed to result in host cell lysis during the time series, with areas of localised dye intensity remaining, consistent with *Ca*. Nha. antarcticus cells being stably internalised within host cells. Images shown are representative of 642 *Hrr. lacusprofundi* cells analysed during 2-23 h. Of these, 360 cells lysed (~ 56%), 33 cells had a rod-shape (circularity < 0.8, Perimeter > 5 μm, ~ 5%), and 12 of the rod-shaped cells exhibited morphological change (~ 36% of the population from rod to cocci). Scale bars represent 1 μm. Images were taken on a Nikon TiE2 microscope with "automatic" 2D deconvolution algorithms in Nikon NIS-Elements AR and were adjusted for bleaching using histogram normalisation in Fiji.

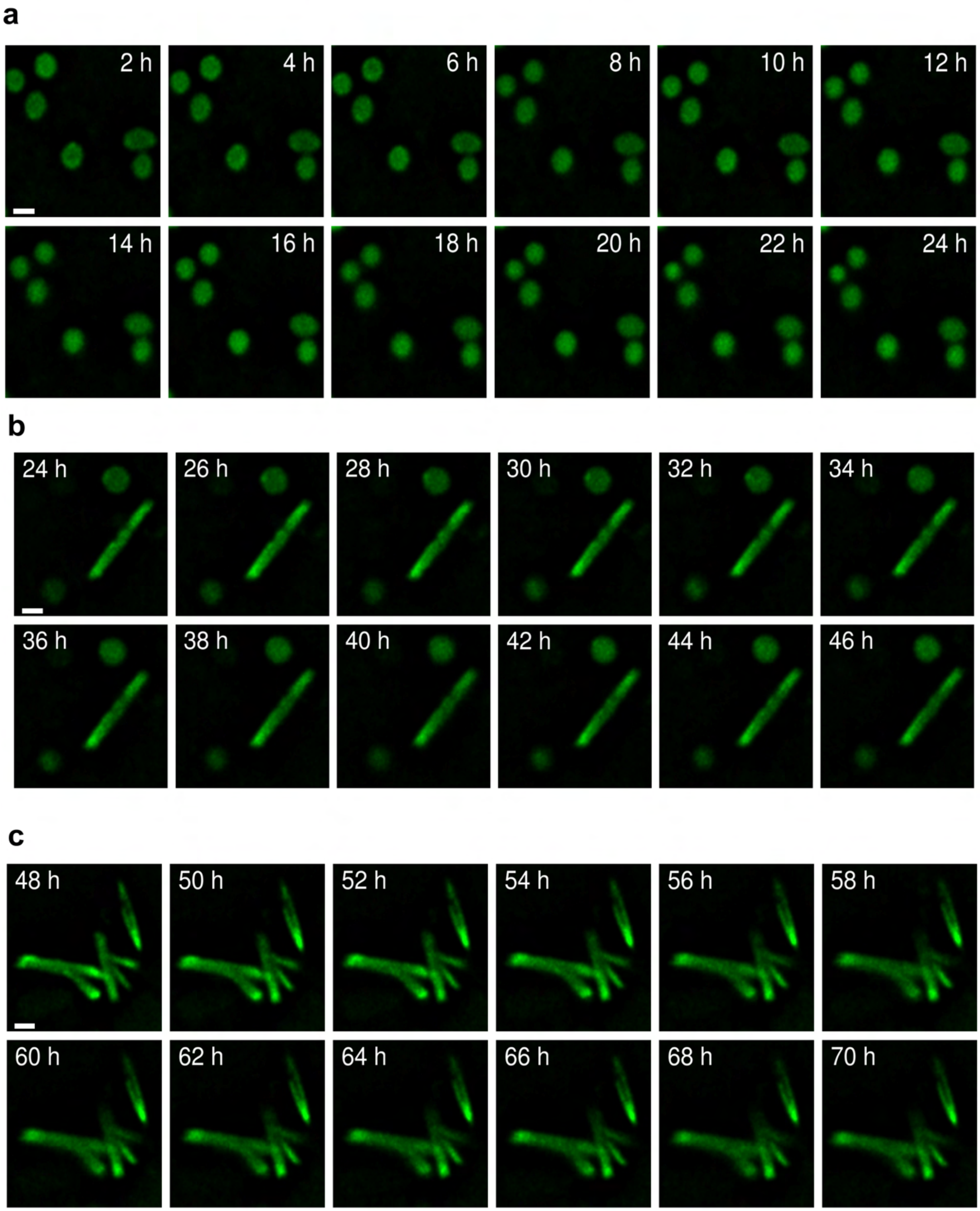


**Extended Data Fig. 4. Live fluorescence microscopy of *Hrr. lacusprofundi.* a - c**, Live fluorescence time lapse series (showing a single focal plane) of *Hrr. lacusprofundi* (stained with MitoTracker Orange, represented green). Across a 70 h period, images of three different sections of the imaging slide were recorded: **a**, 2 h – 24 h; **b**, 24 h – 46 h; **c**, 48 h – 70 h. In this control experiment, without incubation with *Ca.* Nha. antarcticus, the *Hrr. lacusprofundi* cells retained their morphology and were not observed to lyse within 70 h. Scale bars: **a** – **c**, 1 µm. Images were taken on a Delta Vision Elite microscope and then deconvolved and corrected bleaching by softWoRx (enhanced Ratio algorithm with 10 number of cycles, medium noise filtering and applied correction).

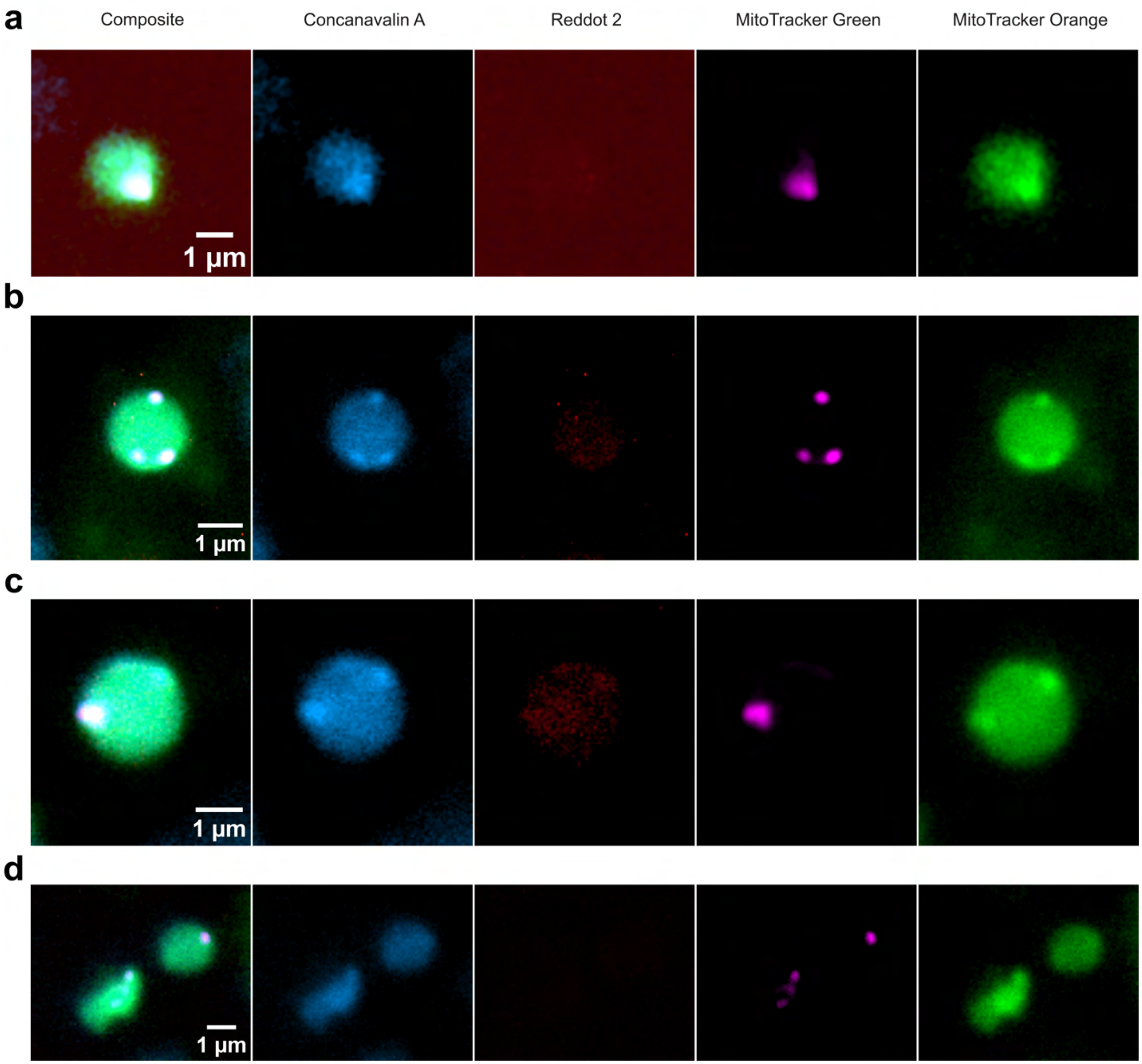


**Extended Data Fig. 5. Live fluorescence microscopy of pre-internalisation *Ca.* Nha. antarcticus - *Hrr. lacusprofundi* interactions. a - d**, Incorporation of a cell surface stain (Concanavalin A, Blue) shows externally attached *Ca.* Nha. antarcticus cells and a dead stain (RedDot 2, Red) shows both cells are viable at this stage of the interaction. *Ca.* Nha. antarcticus cells are visible with both Concanavalin A stain and MitoTracker Green (Magenta) indicating they have not yet internalised within the *Hrr. lacusprofundi* cells. *Hrr. lacusprofundi* cells are visible in Concanavalin A and MitoTracker Orange (Green). No cells are visible in the dead stain indicating they are viable. Scale bars: **a - d,** 1 µm.

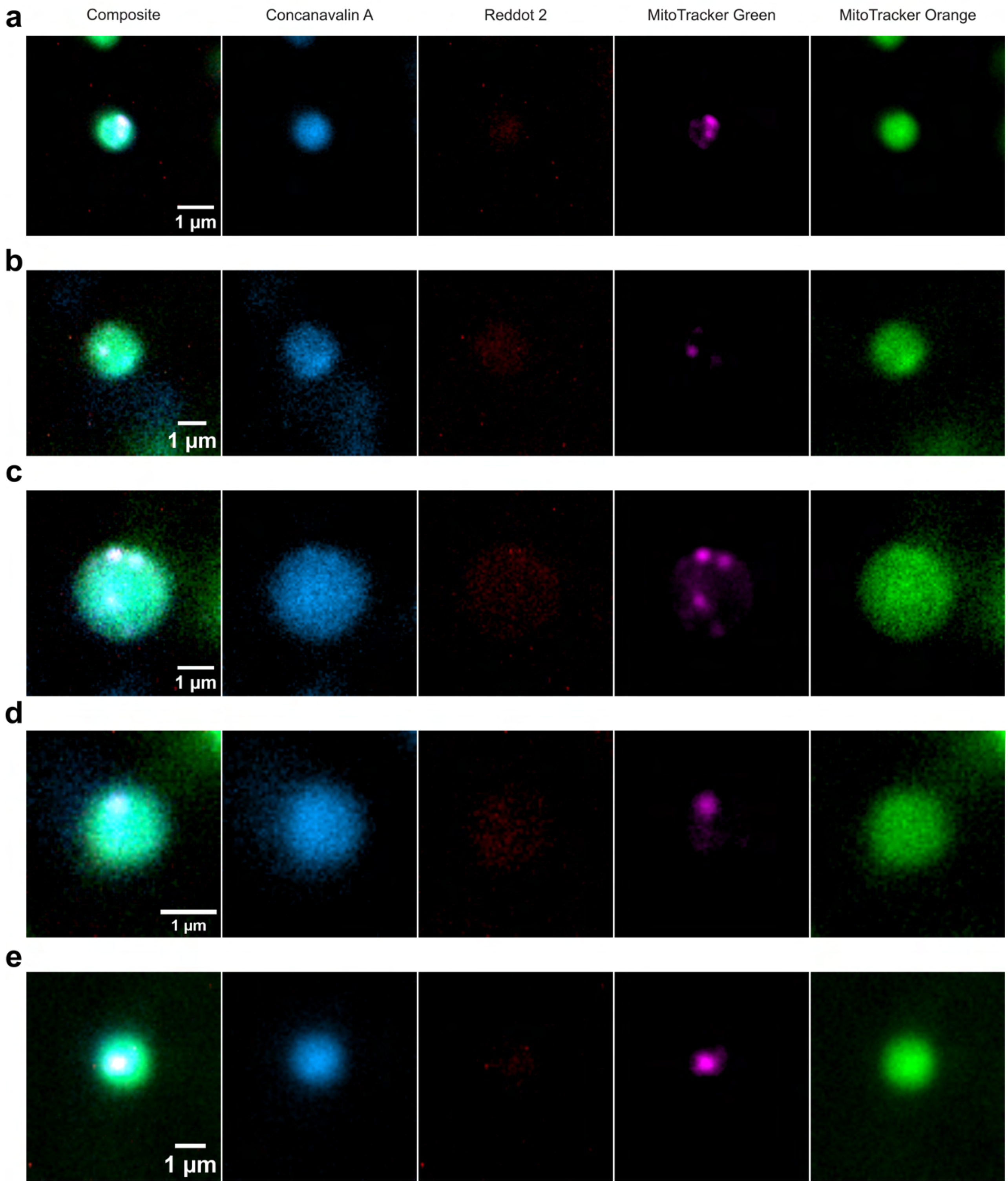


**Extended Data Fig. 6. Live fluorescence microscopy of internalised *Ca.* Nha. antarcticus - *Hrr. lacusprofundi* interactions. a - e**, Cell surface stain (Concanavalin A, Blue) shows internalised *Ca.* Nha. antarcticus cells and a dead stain (RedDot 2, Red) shows both cells are viable at this stage of the interaction. *Ca.* Nha. antarcticus cells are not visible with Concanavalin A stain but are visible with MitoTracker Green (Magenta) indicating they have become internalised within the *Hrr. lacusprofundi* cells. *Hrr. lacusprofundi* cells are visible in Concanavalin A and MitoTracker Orange (Green). No cells are visible in the dead stain indicating they are viable. Scale bars: **a - e,** 1 μm.

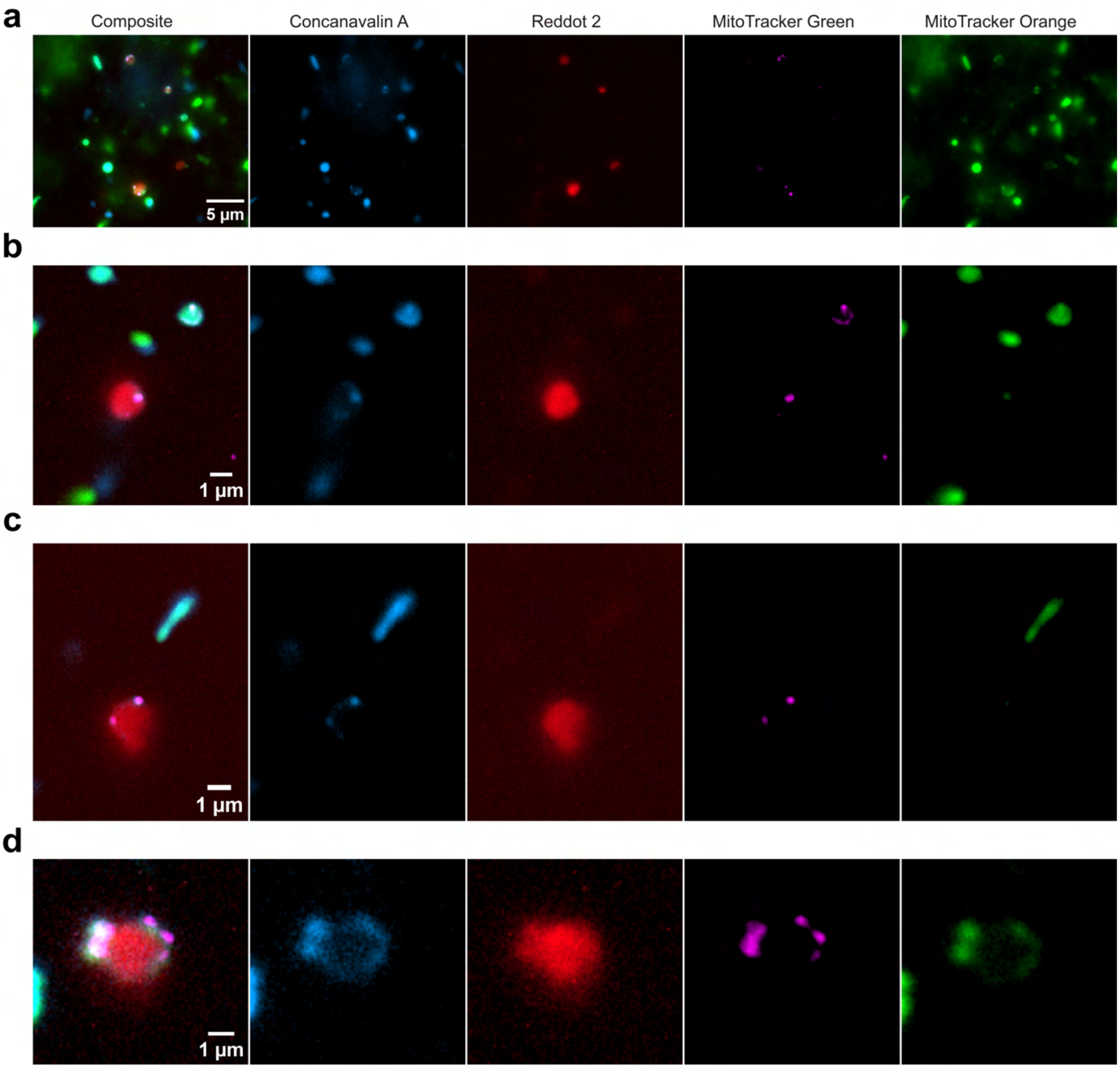


**Extended Data Fig. 7. Live fluorescence microscopy of lysis events during *Ca.* Nha. antarcticus - *Hrr. lacusprofundi* interactions. a - d**, *Hrr. lacusprofundi* cell that showed strong fluorescence for RedDot 2, indicative of loss of membrane integrity and therefore lysis. These lysis events are associated with the presence of *Ca.* Nha. antarcticus cells. Lysis also corresponds to loss of both MitoTracker Orange (Green) and Concanavalin A (Blue) signals. Retention of MitoTracker Green (Magenta) and Concanavalin A signals by associated *Ca.* Nha. antarcticus cells indicates that the nanohaloarchaeal cells remain viable during host lysis. Scale bars: **a - d,** 1 µm.

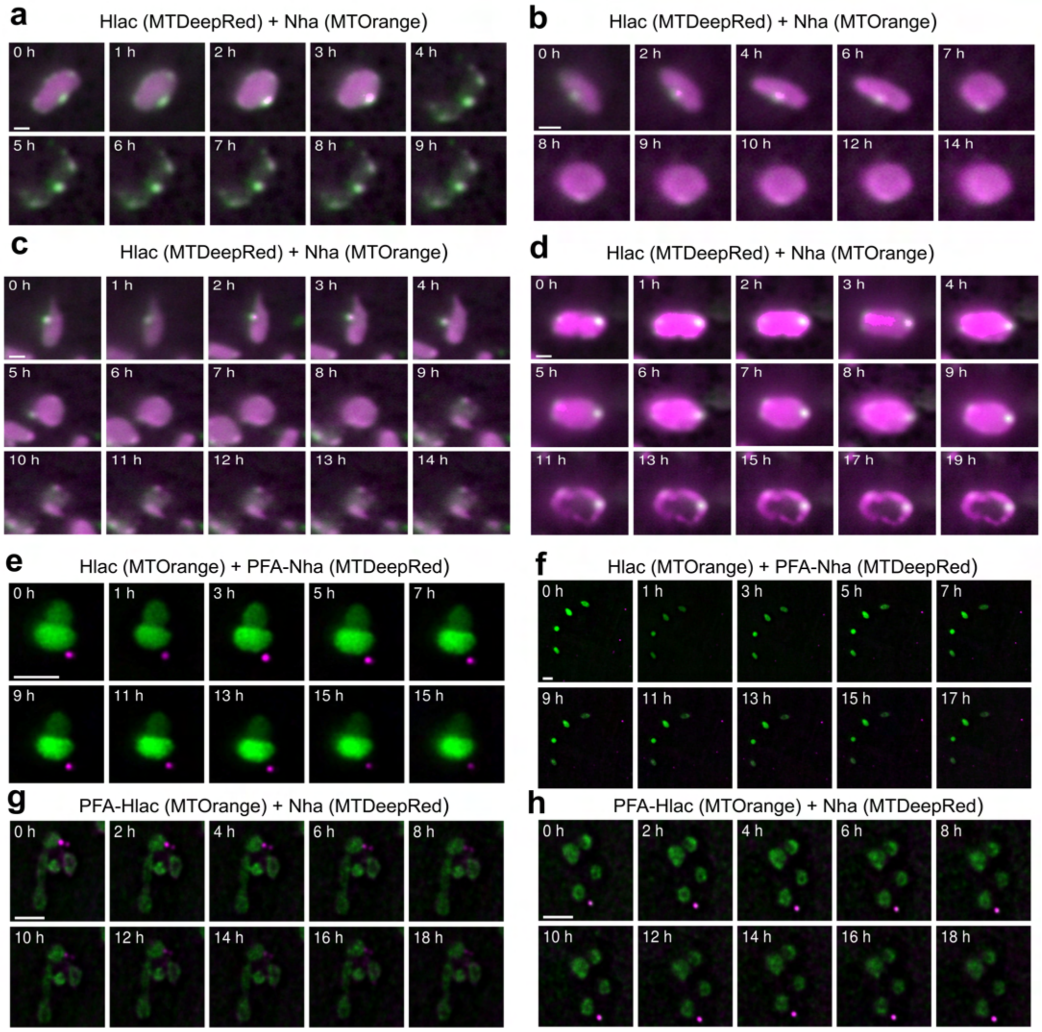


**Extended Data Fig. 8. Live fluorescence microscopy control experiments. a** – **d**, Live fluorescence time lapse series of *Ca*. Nha. antarcticus (stained with MitoTracker Orange, represented green) and *Hrr*. *lacusprofundi* (stained with MitoTracker DeepRed, represented magenta). Images reveal interactions consistent with those observed in previous experiments with the dyes reversed. From 174 *Hrr*. *lacusprofundi* cells analysed during 25 h, 163 cells lysed (~ 94%), 40 cells had a rod-shape (circularity < 0.5, Perimeter > 5 µm, ~ 23%), and 31 of the rod-shaped cells exhibited morphological change (~ 78% of the population from rod to cocci). **a**, Between 3 and 4 h, *Hrr*. *lacusprofundi* fluorescence disappears while *Ca*. Nha. antarcticus fluorescence remains, consistent with host cells lysing leaving *Ca*. Nha. antarcticus cells intact. **b**, *Hrr*. *lacusprofundi* cell morphology changing, most notably from a rod morphology at 6 h to a more coccoid form at 7 h and beyond. **c**, The *Ca*. Nha. antarcticus foci moved along the cell surface of the rod-shaped *Hrr*. *lacusprofundi* cell that changed shape to coccoid form between 4 and 5 h and finally lysed. **d**, Hrr. lacusprofundi cell with *Ca*. Nha. antarcticus cells appeared to lyse between 9 and 10 h. Rather than complete loss of cell integrity, the image is consistent with a disrupted host cell remaining until at least 18 h. **e** – **f**, Live fluorescence time lapse series of paraformaldehyde (PFA) treated *Ca*. Nha. antarcticus cells (stained with MitoTracker DeepRed, represented magenta) and *Hrr*. *lacusprofundi* (stained with MitoTracker Orange, represented green). PFA treated *Ca*. Nha. antarcticus cells displayed reduced capacity to attach to and internalise within *Hrr*. *lacusprofundi* cells. From 186 *Hrr*. *lacusprofundi* cells analysed, 12 cells lysed (~ 6.5%), and no cells showed morphological change. **g** - **h**, Live fluorescence time lapse series of PFA treated *Hrr*. *lacusprofundi* cells (stained with MitoTracker Orange, represented green) and *Ca*. Nha. antarcticus cells (stained with MitoTracker DeepRed, represented magenta). Fixed *Hrr*. *lacusprofundi* cells reduced the capacity for *Ca*. Nha. antarcticus to attach to and internalise within *Hrr*. *lacusprofundi* cells. From 265 *Hrr*. *lacusprofundi* cells analysed, no cells showed morphological changes or lysis.

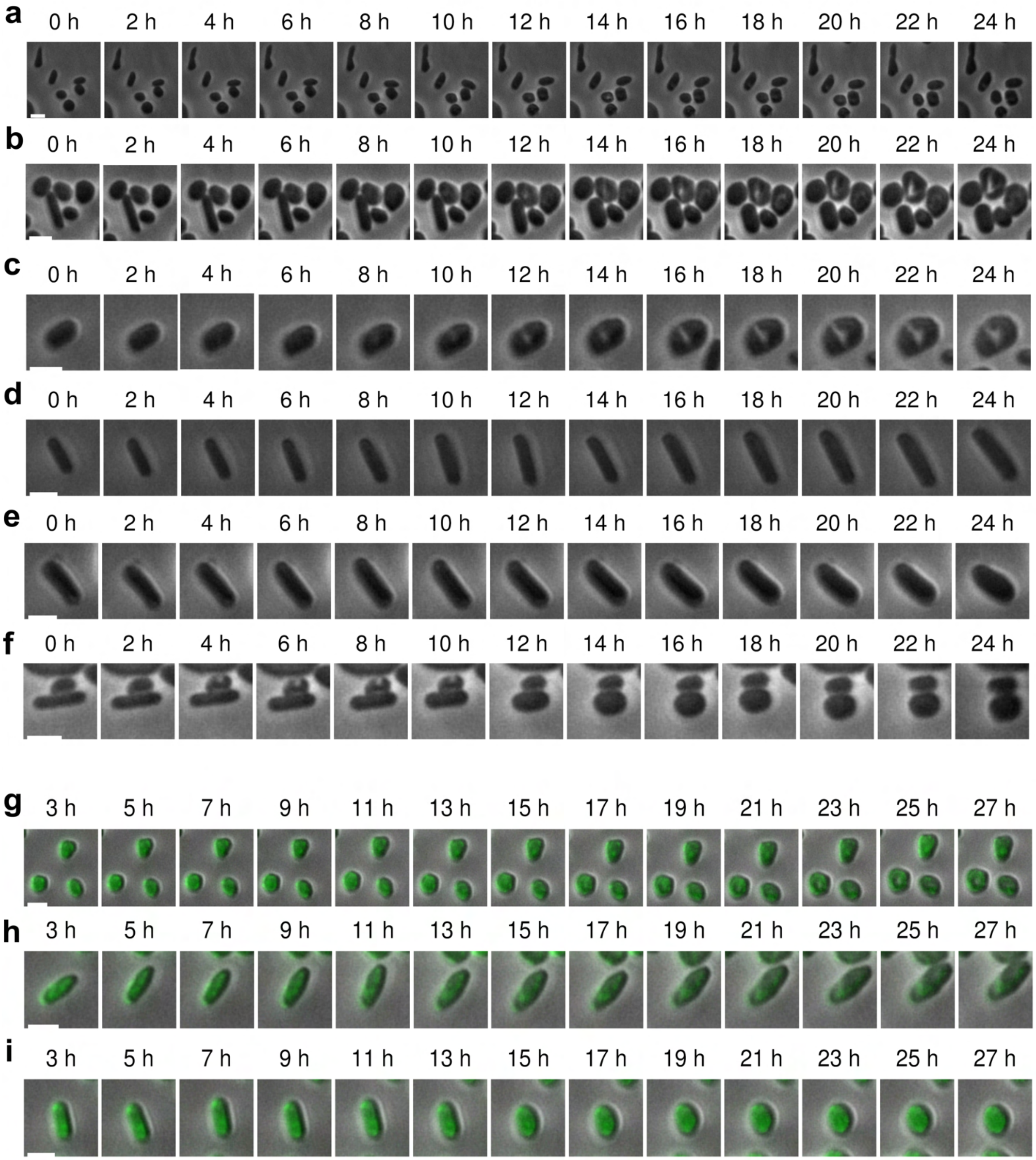


**Extended Data Fig. 9. CellASIC Live fluorescence microscopy control experiments. a** - **f**, Unstained pure *Hrr. lacusprofundi* cells. During 0 - 24 h, no lysis was observed for 407 *Hrr. lacusprofundi* cells; 59 cells exhibited rod morphology (circularity < 0.8, perimeter > 5 µm, ~ 15%) of which only one (~1.7% of rod-shaped population) underwent morphological changes from rod to cocci. **g** - **i**, MitoTracker Orange (represented green) stained pure *Hrr. lacusprofundi* cells. From 654 *Hrr. lacusprofundi* cells analyzed, 2 cells lysed (~ 0.3%) during 3-27 h; 21 cells had rod morphology (circularity < 0.8, perimeter > 5 µm), and 4 of them exhibited morphological changes (~ 19% of rod-shaped population, from rod to cocci). The proportion of control *Hrr. lacusprofundi* cells that lysed (~ 0-0.3%) or underwent morphological change (~ 1.7-19% of rod-shaped cells) was markedly lower than the number of *Hrr. lacusprofundi* cells that lysed (~ 56%) or underwent morphological change (~ 36% of rod-shaped cells) after they were incubated with *Ca*. Nha. antarcticus cells (Extended Data Fig. 3). The cell area of pure unstained or stained *Hrr. lacusprofundi* cells gradually increased. Also see cell size characteristics in Extended Data Fig. 10. Scale bars: **a** - **i,** 2 µm. Images were taken on a Nikon TiE2 microscope and were adjusted for bleaching using histogram normalisation in Fiji (**g** - **i**).

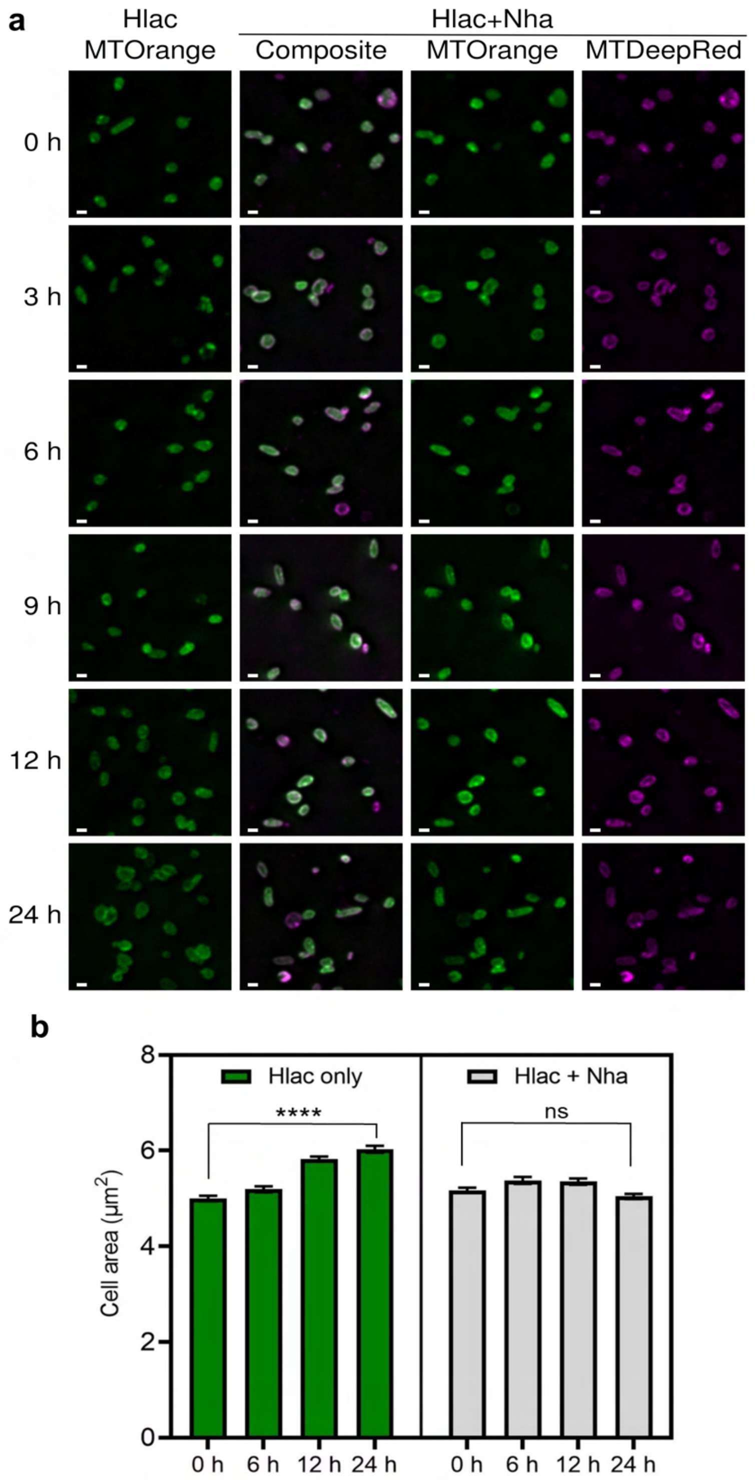


**Extended Data Fig. 10. Cell size and shape characteristics. a**, Fluorescent microscopy of H*rr. lacusprofundi* only (R1S1) as a control and *Hrr. lacusprofundi* incubated with *Ca.* Nha. antarcticus (R1S1+Nha) sampled at 0, 3, 6, 9, 12, and 24 h. The *Hrr. lacusprofundi* cells were stained with MitoTracker Orange (represented green) and *Ca*. Nha. antarcticus were stained with MitoTracker DeepRed (represented magenta). Scale bars, 2 µm. Images were taken on a Delta Vision Elite microscope and then deconvolved by softWoRx and shown at single focal plane. **b**, Cell area analysis of *Hrr. lacusprofundi* only and mixed culture at 0, 6, 12, and 24 h. The values of cell area value at each time point are displayed as mean with SEM [Hlac only: 0 h (n=933), 6 h (n=971), 12 h (n=1194), 24 h (1189); Hlac + Nha: 0 h (n= 945), 6 h (n=723), 12 h (n=823), 24 h (n=1218). Statistical significance of cell area between 0 h and 24 h was calculated using the two-tailed unpaired Mann–Whitney U-test (**** : P < 0.0001; ns : not significant).

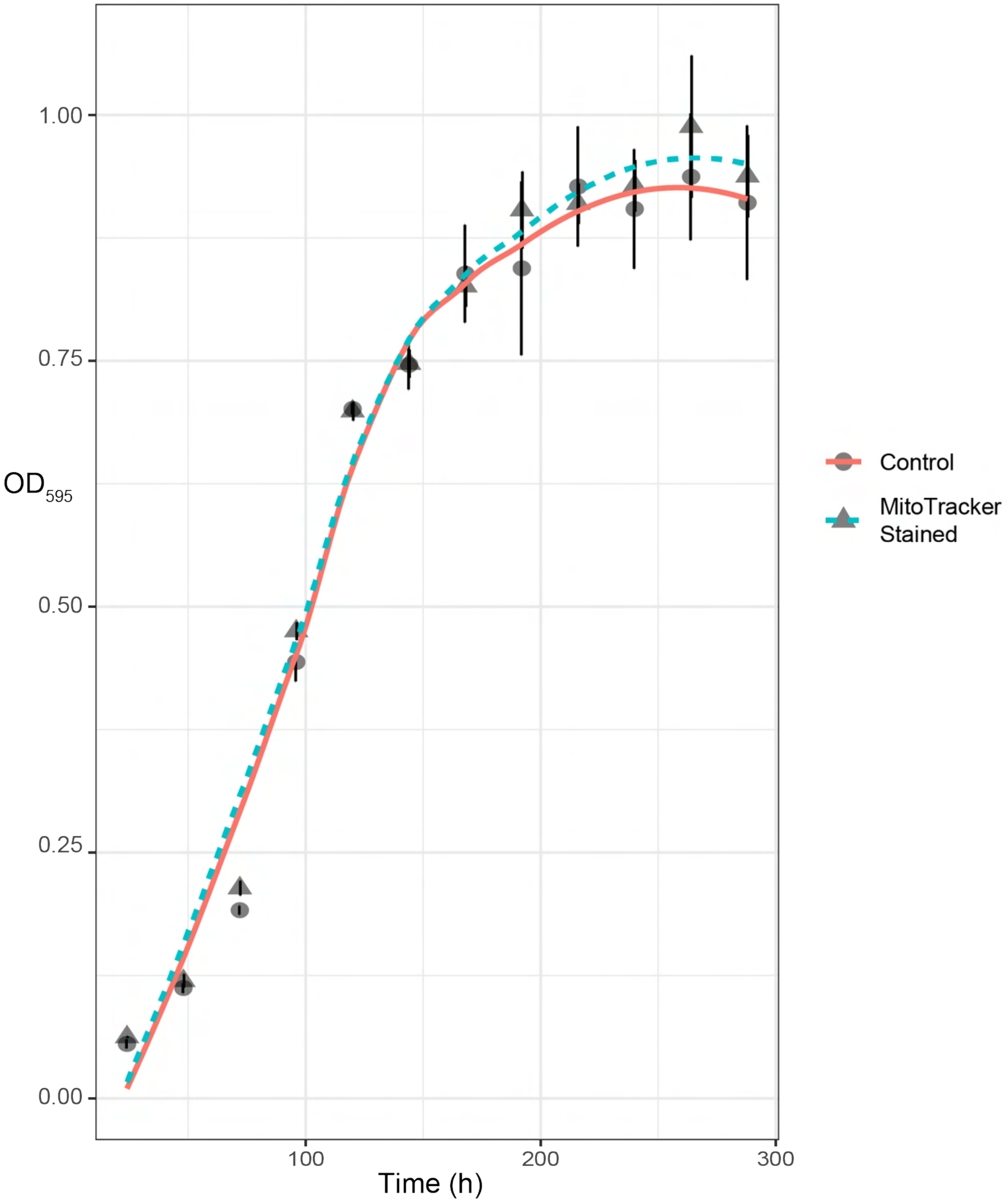


**Extended Data Fig. 11. Growth curve of MitoTracker stained *Hrr. lacusprofundi*.** Growth characteristics of *Hrr. lacusprofundi* stained with MitoTracker Orange compared with unstained *Hrr. lacusprofundi*. MitoTracker stained *Hrr. lacusprofundi* cultures (Triangle symbols, blue line) displayed no apparent difference in growth characteristics compared to unstained *Hrr. lacusprofundi* cultures (Circle symbols, red dotted line). Data points represent six replicates; error bars represent standard deviation across the six replicates.

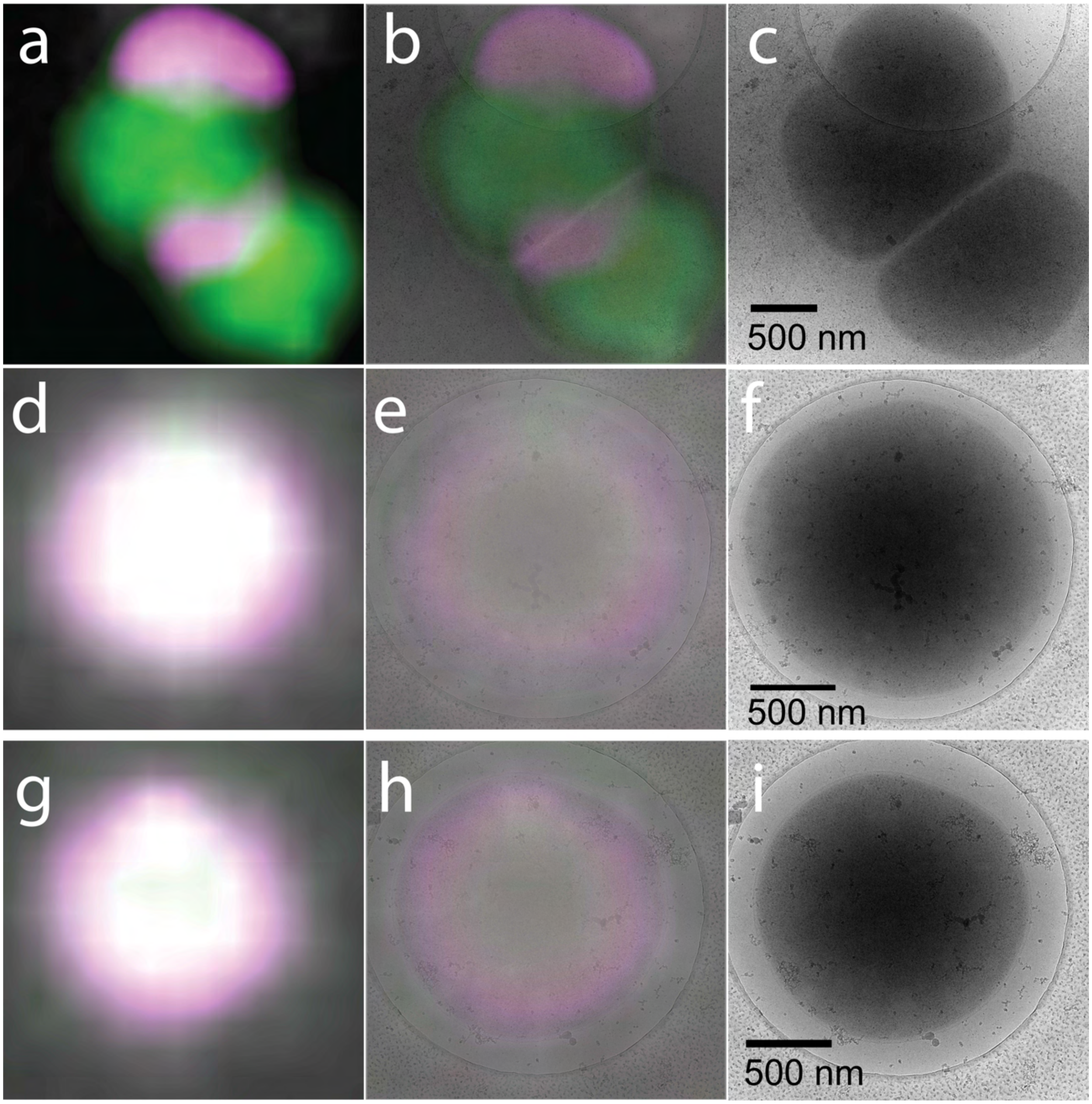


**Extended Data Fig. 12. Cryo-CLEM of internalised nanohaloarchaeal cells.** Cryo-CLEM of *Ca.* Nha. antarcticus (stained with MitoTracker DeepRed, represented magenta) and *Hrr. lacusprofundi* (stained with MitoTracker Orange, represented green) following incubation for 20 h. The colour panels (**a**, **d**, **g**) are confocal fluorescence images, colour panels (**b**, **e**, **h**) are composites of the confocal-fluorescence and cryo-electron microscopy images of the same cell and the grey-scale panels (**c**, **f**, **i**) are the corresponding cryo-TEM micrographs. *Hrr. lacusprofundi* cells that displayed fluorescence signals suggestive of internalised *Ca.* Nha. antarcticus cells did not show any signs of membrane disruption or lysis in the corresponding cryo-TEM micrographs indicating host cells were likely still viable. *Ca.* Nha. antarcticus cells were not visible externally attached, consistent with the nanohaloarchaeal cells having fully internalised within the *Hrr. lacusprofundi* cell. Electron micrographs were acquired on a Talos Arctica at 200 kV.

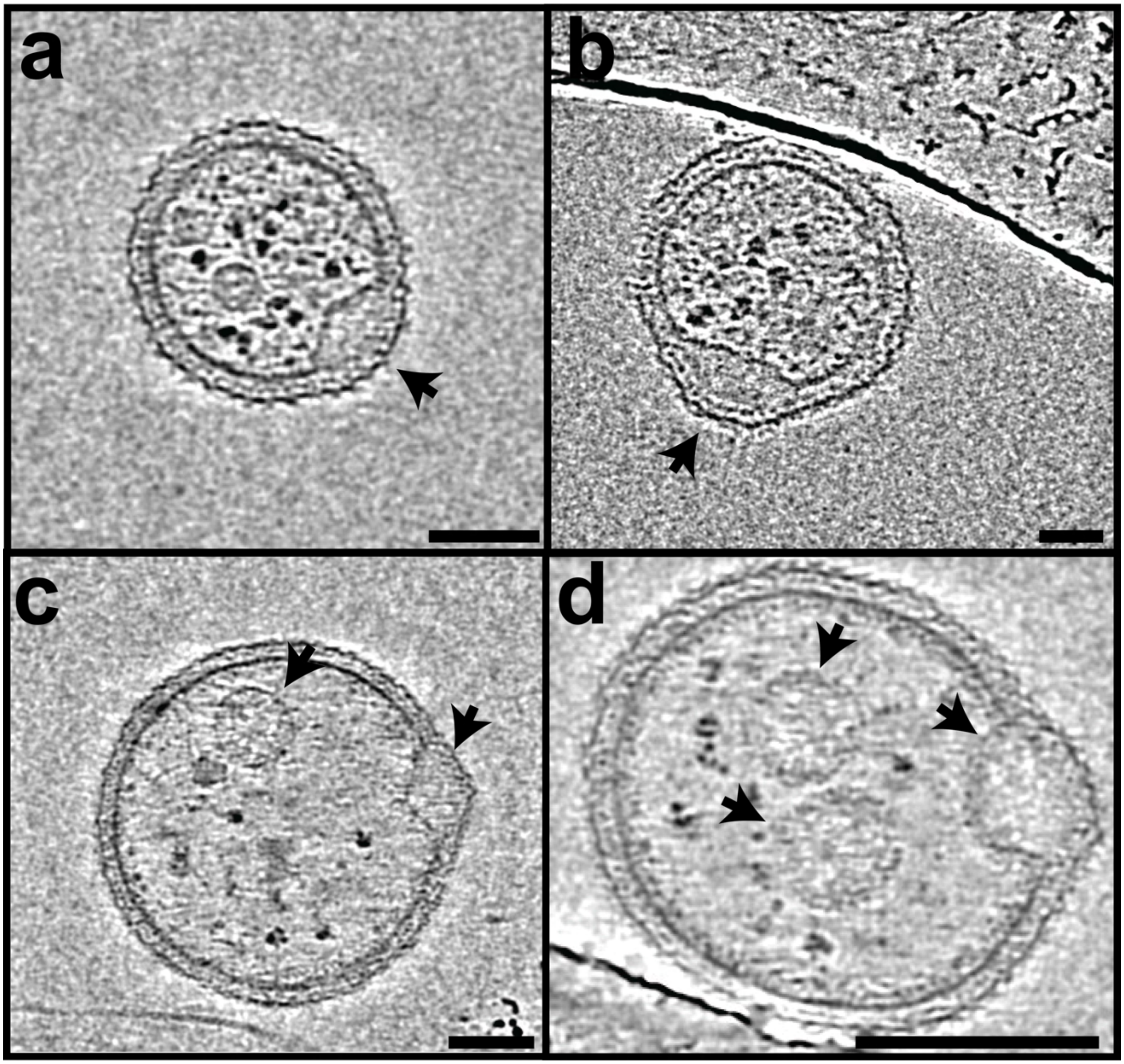


**Extended Data Fig. 13. Cryo-ET of unattached *Ca*. Nha. antarcticus cells. a** – **d**, Z-slices from tomograms (**a**: Supplementary Video 5, **b**: Supplementary Video 6, **c**: Supplementary Video 7, **d**: Supplementary Video 8) of unattached *Ca*. Nha. antarcticus cells showing structural features of nanohaloarchaeal cells. Z-slices reveal that *Ca*. Nha. antarcticus cells frequently possessed lipid droplets embedded within the bilayer that forms the cell membrane (**a** – **d**, arrows). Nanohaloarchaeal cells were also observed with cytoplasmic membrane bound structures likely to be PHA-like granules acquired during interactions with *Hrr. lacusprofundi* (**c** and **d**, arrow). Images acquired using a Titan Krios at 300 kV. Scale bars: **a** - **c** - 50 nm, **d** - 100 nm

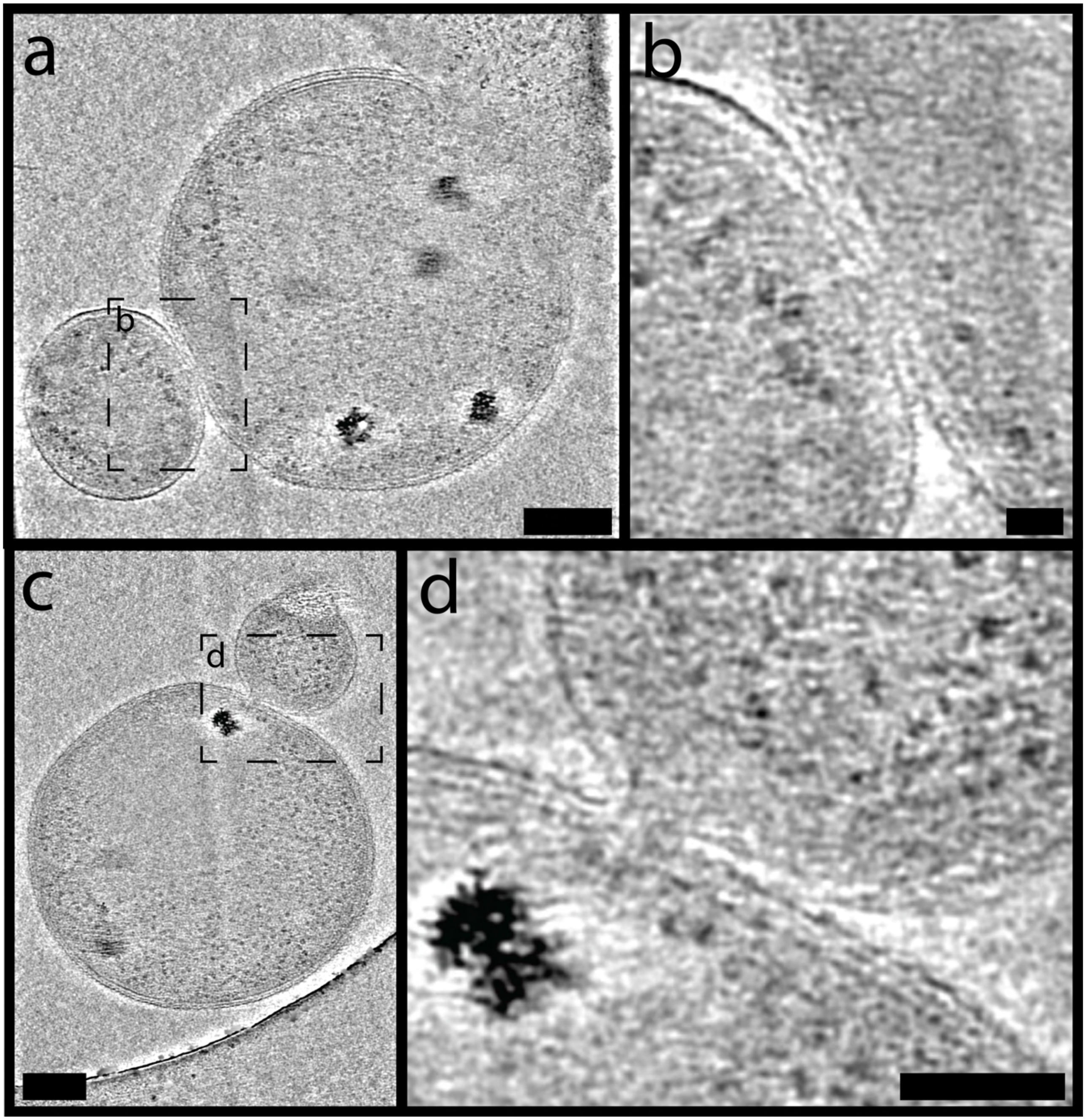


**Extended Data Fig. 14. Cryo-ET of interactions between *Ca.* Nha. antarcticus and *Hrr. lacusprofundi* cells. a** – **d,** Single Z-slices taken from cryo-tomograms (**a** and **b**: Supplementary Video 9, **c** and **d**: Supplementary Video 10) showing interactions between externally attached *Ca.* Nha. antarcticus cells and *Hrr. lacusprofundi* cells. Cell membranes at contact sites show signs of localised deformation and fusion (**b** and **d**) including an apparent direct channel between the cytoplasms of the two organisms (**b**). Scale bars: **a** and **c** - 100 nm, **b** - 20 nm, **d** - 50 nm

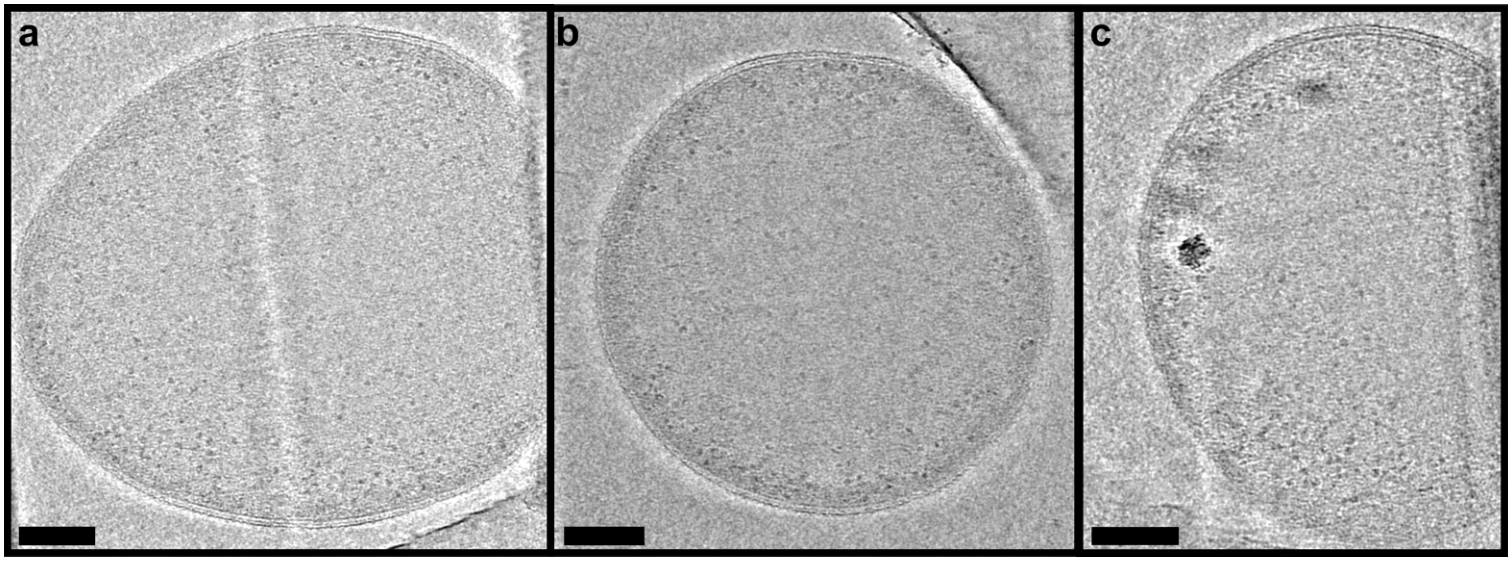


**Extended Data Fig. 15. Cryo-ET of pure *Hrr. lacusprofundi cells*. a** – **c,** Single Z-slices taken from cryo-tomograms (**a**: Supplementary Video 11, **b**: Supplementary Video 12, **c**: Supplementary Video 13) of pure *Hrr. lacusprofundi* cells show no evidence of internal structures resembling the membrane bound structures observed within *Hrr. lacusprofundi* cells incubated with *Ca.* Nha. antarcticus. All scale bars 100 nm

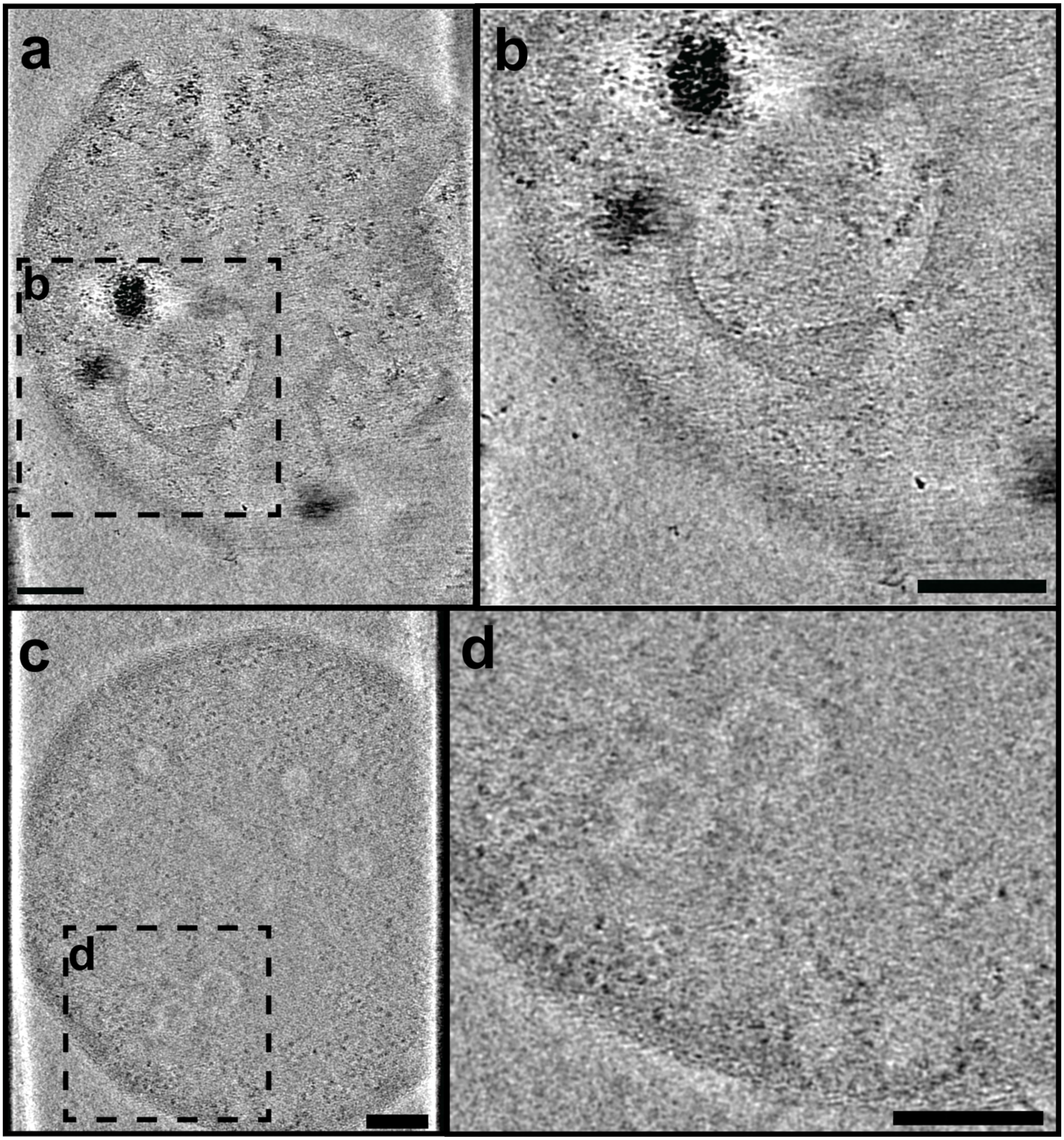


**Extended Data Fig. 16. Cryo-tomography of radiation sensitive internal membrane bound structures. a** – **d**, Z-slices from tomograms (**a** and **b**: Supplementary Video 14, **c** and **d**: Supplementary Video 15) showing internal membrane bound structures within *Hrr. lacusprofundi* cells consistent with *Ca*. Nha. antarcticus that have internalised. Panels show both (**a** and **c**) full field of view and (**b** and **d**) internalised membrane bound structures. **a** and **b,** A Hrr. lacusprofundi cell that has lysed (membrane break indicated by black arrow in panel **a**) that contains an internal membrane bound structure consistent with a Ca. Nha. antarcticus cell (**b**). **c** and **d,** A *Hrr. lacusprofundi* cell that appears intact with internal membrane bound structures. Internalised membrane bound structures displayed higher radiation sensitivity during imaging than other cellular features. Images acquired using a Titan Krios at 300 kV. All scale bars: 100 nm.

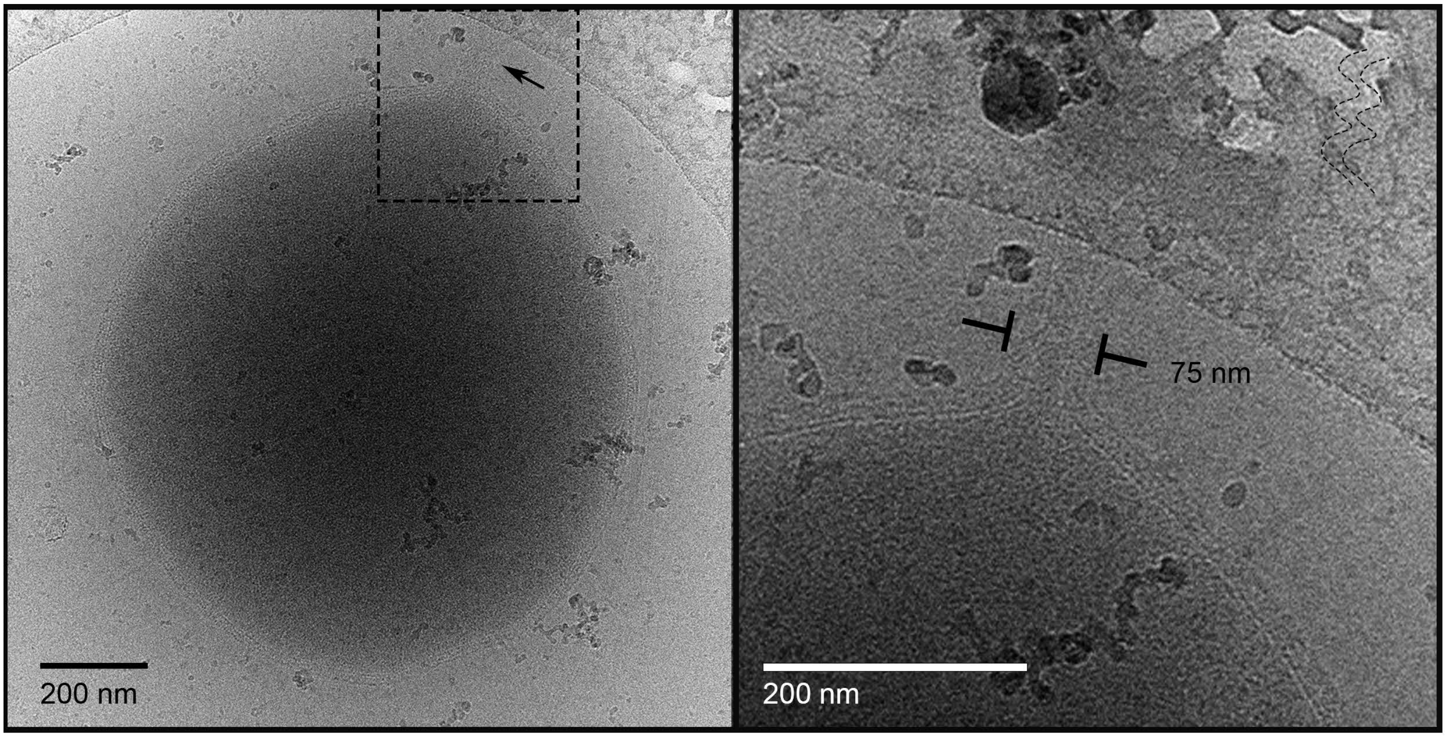


**Extended Data Fig. 17. Cryo-EM of *Ca.* Nha. antarcticus cytoplasmic bridges and internalisation structures. a** – **b**, Cryo-TEM micrograph showing cytoplasmic bridge of a *Ca.* Nha. antarcticus cell. **a**, A cytoplasmic bridge extended out from a nanohaloarchaeal cell (arrow), with the expanded view of the bridge (**b**). There is no visible boundary between the contents of the bridge and the nanohaloarchaeal cytoplasm. Images acquired on a Talos Arctica at 200 kV

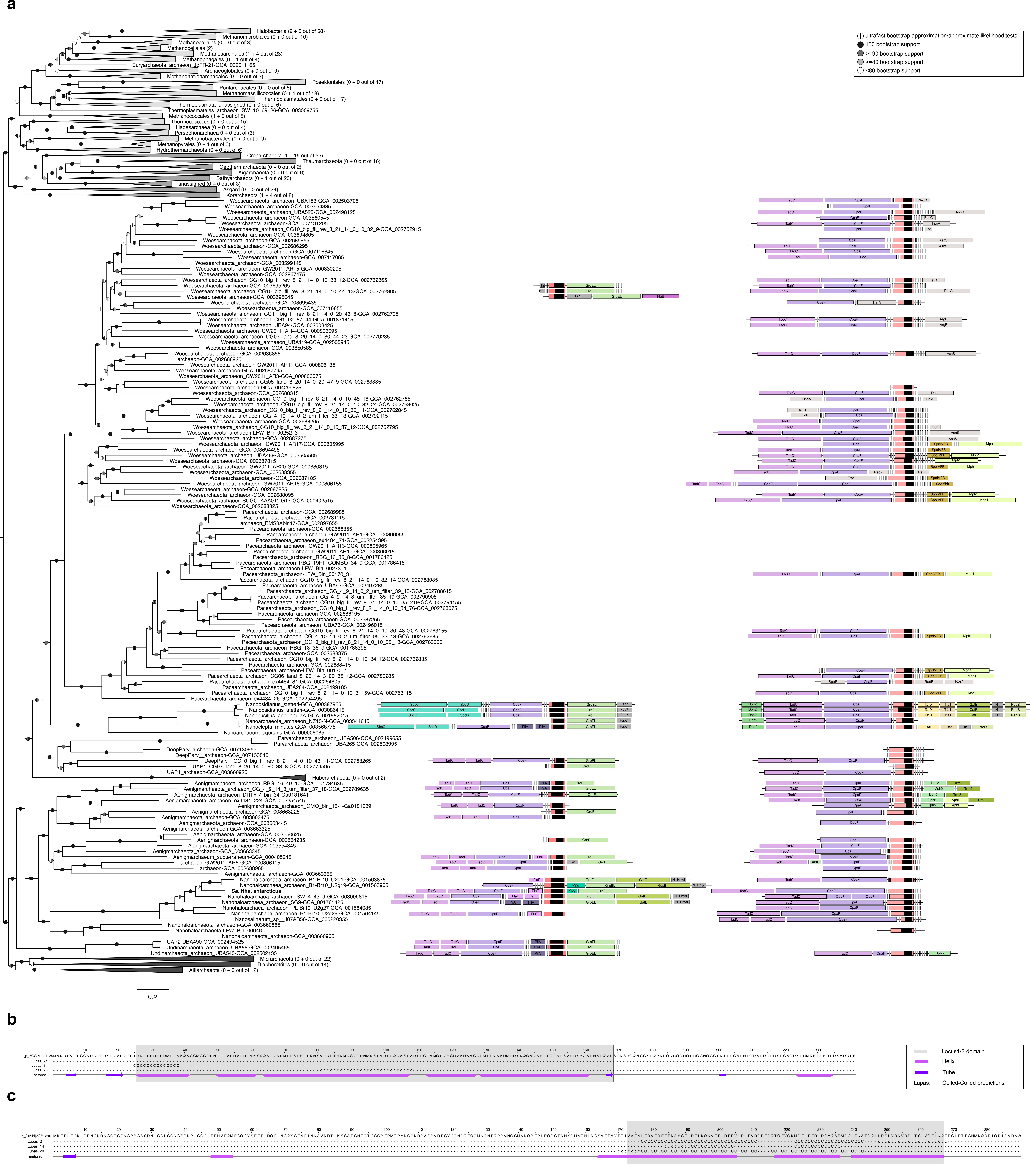


**Extended Data Fig. 18. Conservation of loci encoding CCP genes in Nanohaloarchaeota across Archaea. a**, The maximum-likelihood phylogenetic tree was based on 51 marker proteins and 569 archaeal species. The alignment was trimmed with BMGE (alignment length, 11399 aa). A phylogenetic tree was inferred in IQ-TREE with the LG+C20+F+R model with an ultrafast bootstrap approximation (left half of bootstrap symbol) and SH-like approximate likelihood test (right half of bootstrap symbol), each run with 1000 replicates (see key for shading indicating bootstrap support). The tree was artificially rooted between DPANN and all other Archaea. The number of species represented in each clade is shown in parentheses after the taxonomic name of the clade. Scale bar: Average number of substitutions per site. The two, Cluster 2 DPANN loci are shown aligned to their respective sequences in the phylogenetic tree. **b**, **c**, Secondary structure predictions inferred with JPred4 for the CCP from Locus 1 (**b**) and CCP from Locus 2 (**c**). For further details, see Fig. 4.

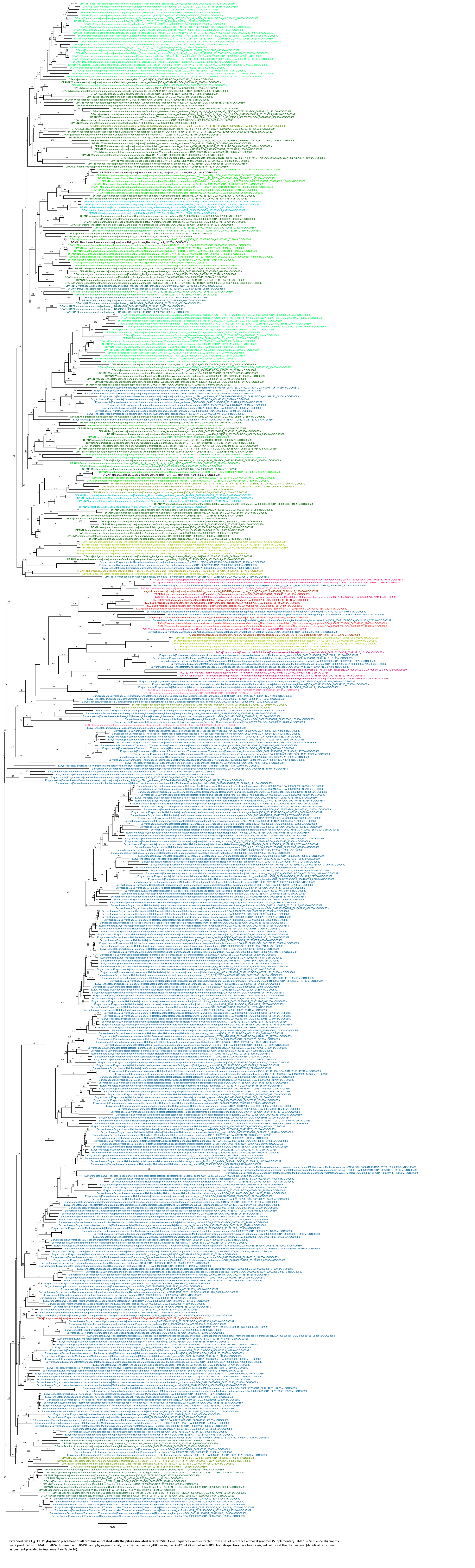

**Extended Data Fig. 19. Phylogenetic placement of all proteins annotated with the pilus associated arCOG00589.** Gene sequences were extracted from a set of reference archaeal genomes (Supplementary Table 13). Sequence alignments were produced with MAFFT L-INS-i, trimmed with BMGE, and phylogenetic analysis carried out with IQ-TREE using the LG+C10+F+R model with 1000 bootstraps. Taxa have been assigned colours at the phylum level (details of taxonomic assignment provided in Supplementary Table 16).

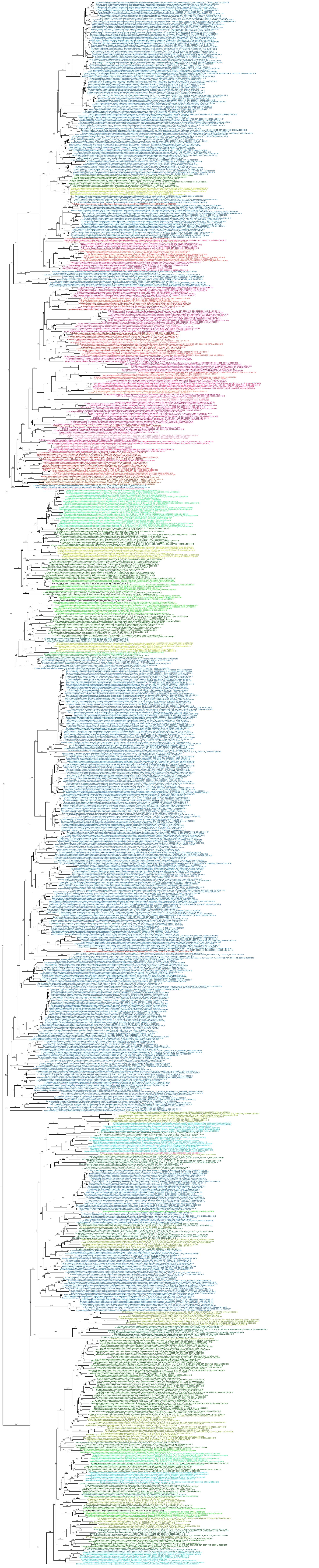

**Extended Data Fig. 20. Phylogenetic placement of all proteins annotated with the pilus associated arCOG01819.** Gene sequences were extracted from a set of reference archaeal genomes (Supplementary Table 16). Sequence alignments were produced with MAFFT L-INS-i, trimmed with BMGE, and phylogenetic analysis carried out with IQ-TREE using the LG+C10+F+R model with 1000 bootstraps. Taxa have been assigned colours at the phylum level (details of taxonomic assignment provided in Supplementary Table 16).

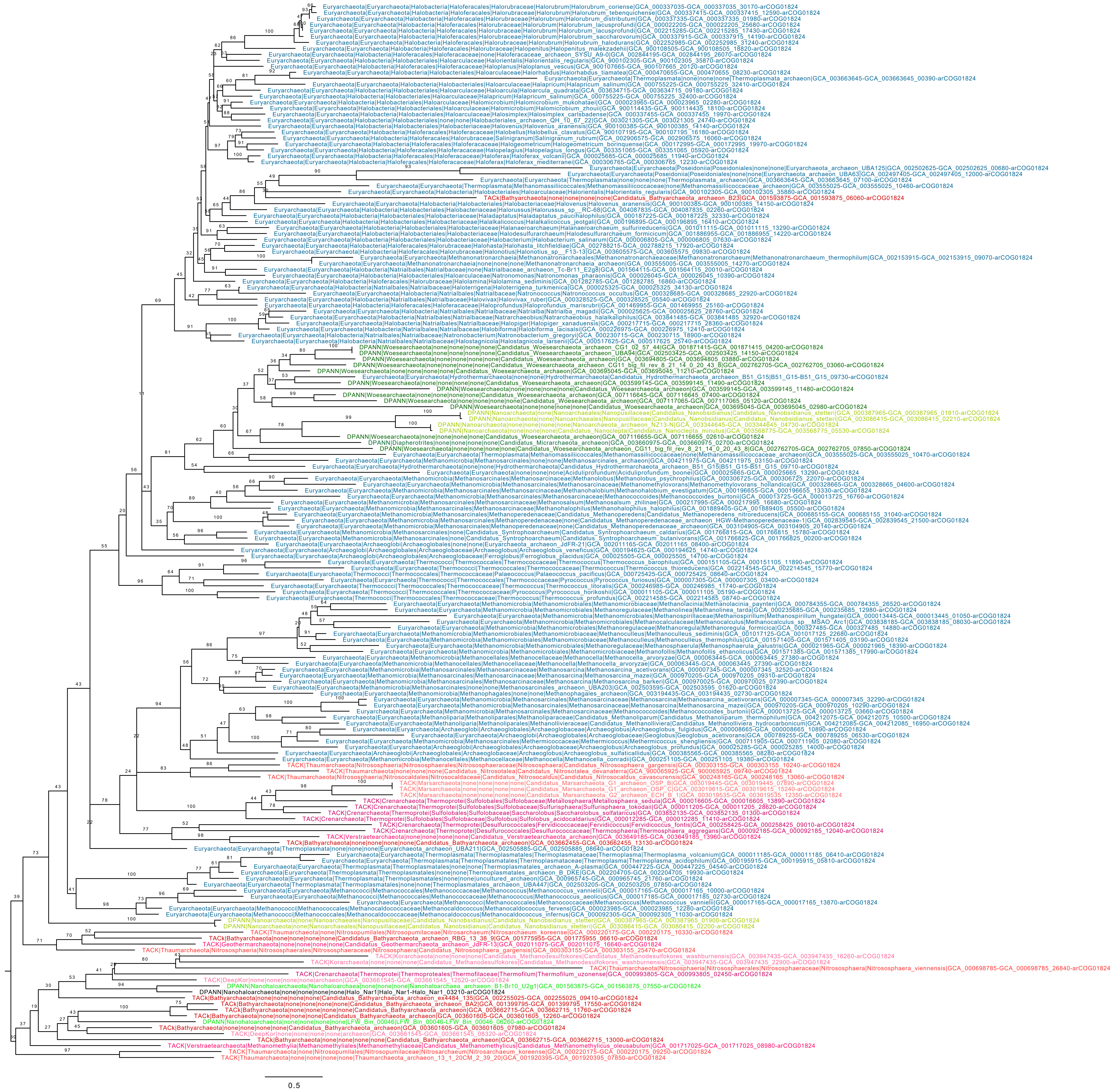

**Extended Data Fig. 21. Phylogenetic placement of all proteins annotated with the pilus associated arCOG001824.** Gene sequences were extracted from a set of reference archaeal genomes (Supplementary Table 16). Sequence alignments were produced with MAFFT L-INS-i, trimmed with BMGE, and phylogenetic analysis carried out with IQ-TREE using the LG+C10+F+R model with 1000 bootstraps. Taxa have been assigned colours at the phylum level (details of taxonomic assignment provided in Supplementary Table 16).

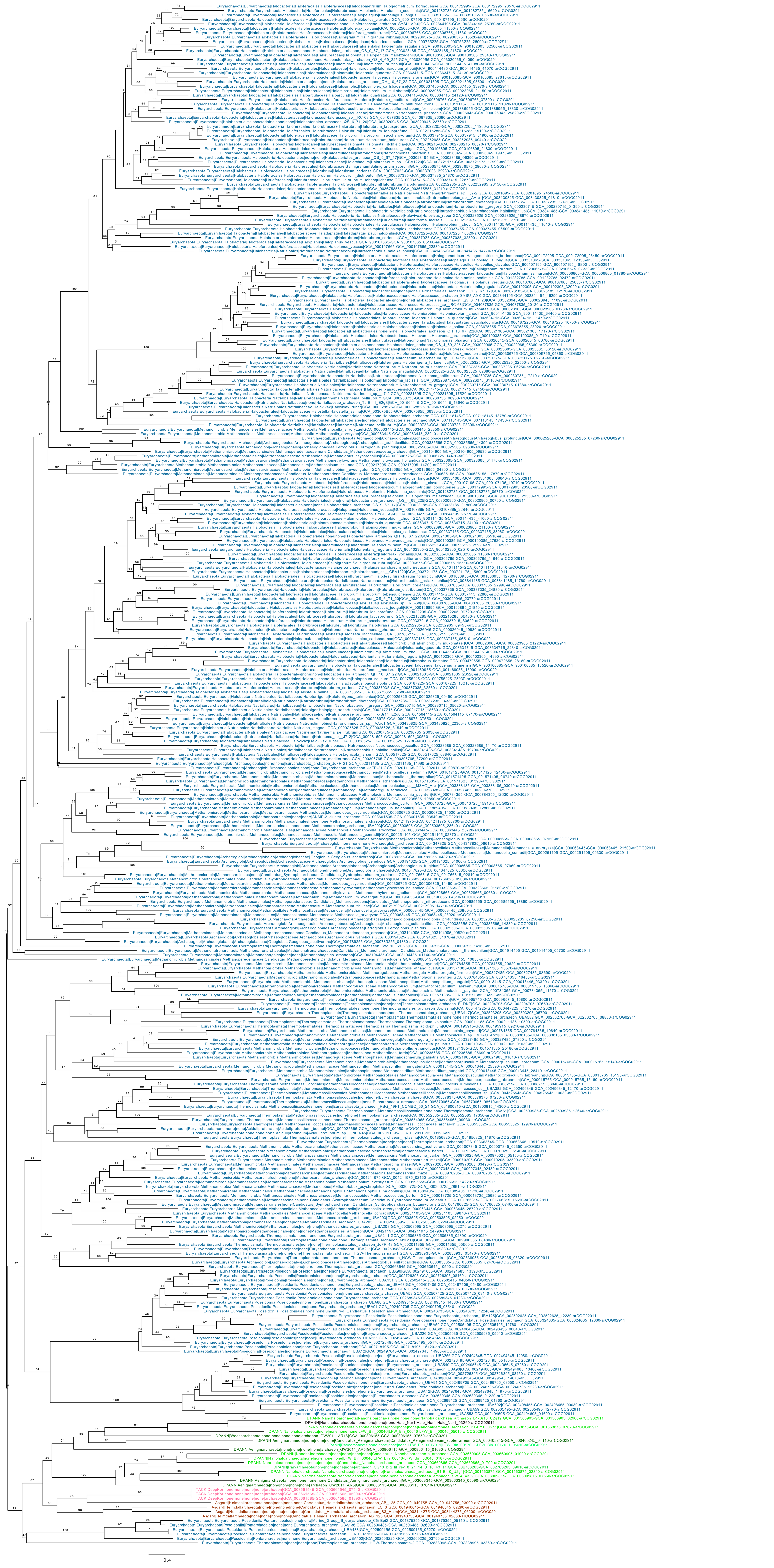

**Extended Data Fig. 22. Phylogenetic placement of all proteins annotated with the pilus associated arCOG02911.** Gene sequences were extracted from a set of reference archaeal genomes (Supplementary Table 16). Sequence alignments were produced with MAFFT L-INS-i, trimmed with BMGE, and phylogenetic analysis carried out with IQ-TREE using the LG+C10+F+R model with 1000 bootstraps. Taxa have been assigned colours at the phylum level (details of taxonomic assignment provided in Supplementary Table 16).

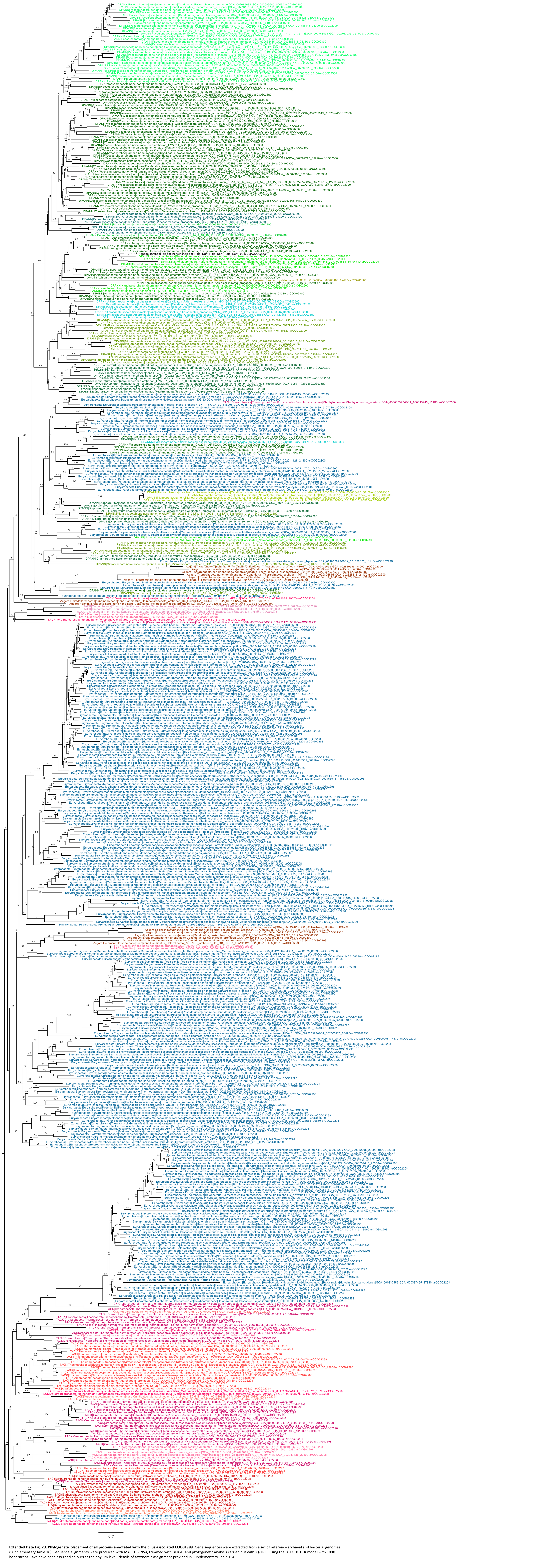


**Extended Data Fig. 23. Phylogenetic placement of all proteins annotated with the pilus associated COG01989.** Gene sequences were extracted from a set of reference archaeal and bacterial genomes (Supplementary Table 16). Sequence alignments were produced with MAFFT L-INS-i, trimmed with BMGE, and phylogenetic analysis carried out with IQ-TREE using the LG+C10+F+R model with 1000 boot-straps. Taxa have been assigned colours at the phylum level (details of taxonomic assignment provided in Supplementary Table 16).

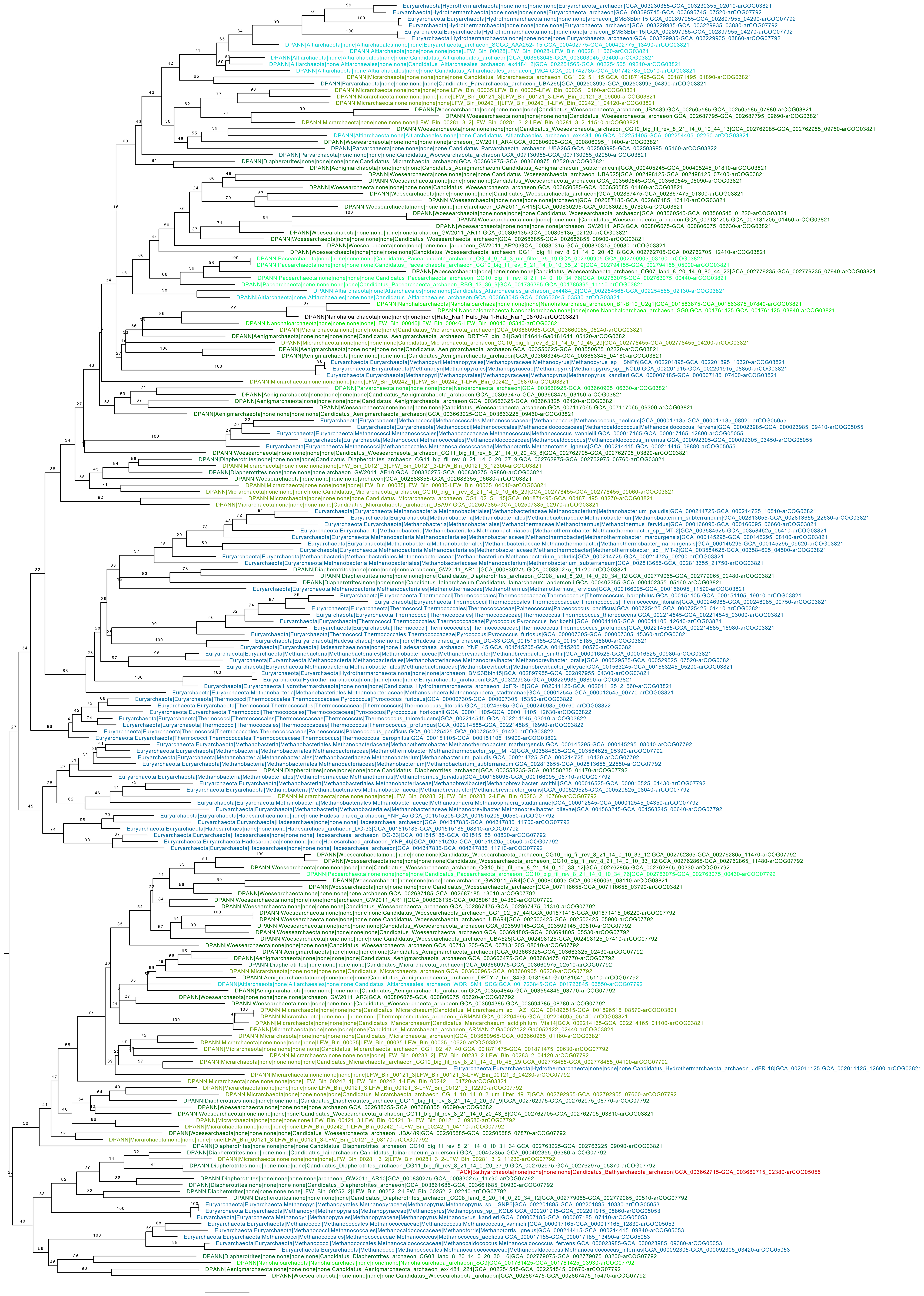

**Extended Data Fig. 24. Phylogenetic placement of all proteins annotated with the pilus associated COG01991b.** Gene sequences were extracted from a set of reference archaeal and bacterial genomes (Supplementary Table 16). Sequence alignments were produced with MAFFT L-INS-i, trimmed with BMGE, and phylogenetic analysis carried out with IQ-TREE using the LG+C10+F+R model with 1000 bootstraps. Taxa have been assigned colours at the phylum level (details of taxonomic assignment provided in Supplementary Table 16).

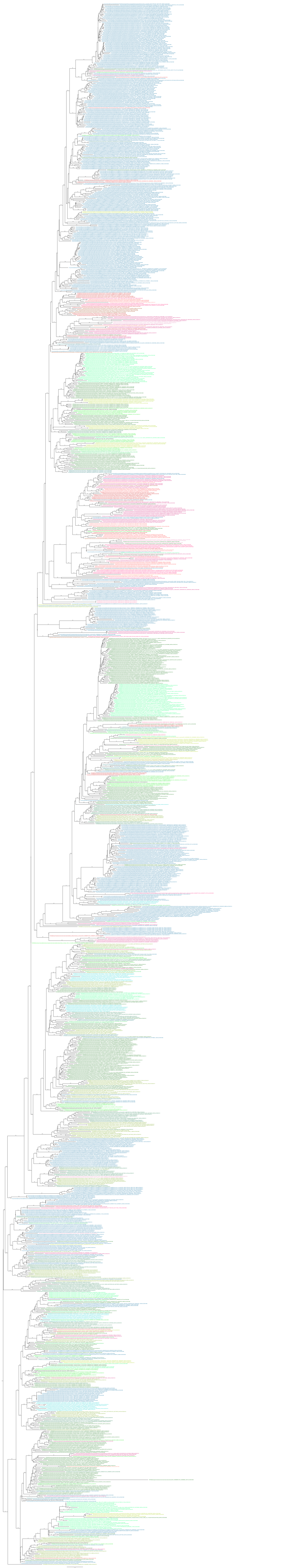

**Extended Data Fig. 25. Phylogenetic placement of all proteins annotated with the pilus associated COG02064b.** Gene sequences were extracted from a set of reference archaeal and bacterial genomes (Supplementary Table 16). Sequence alignments were produced with MAFFT L-INS-i, trimmed with BMGE, and phylogenetic analysis carried out with IQ-TREE using the LG+C10+F+R model with 1000 bootstraps. Taxa have been assigned colours at the phylum level (details of taxonomic assignment provided in Supplementary Table 16).

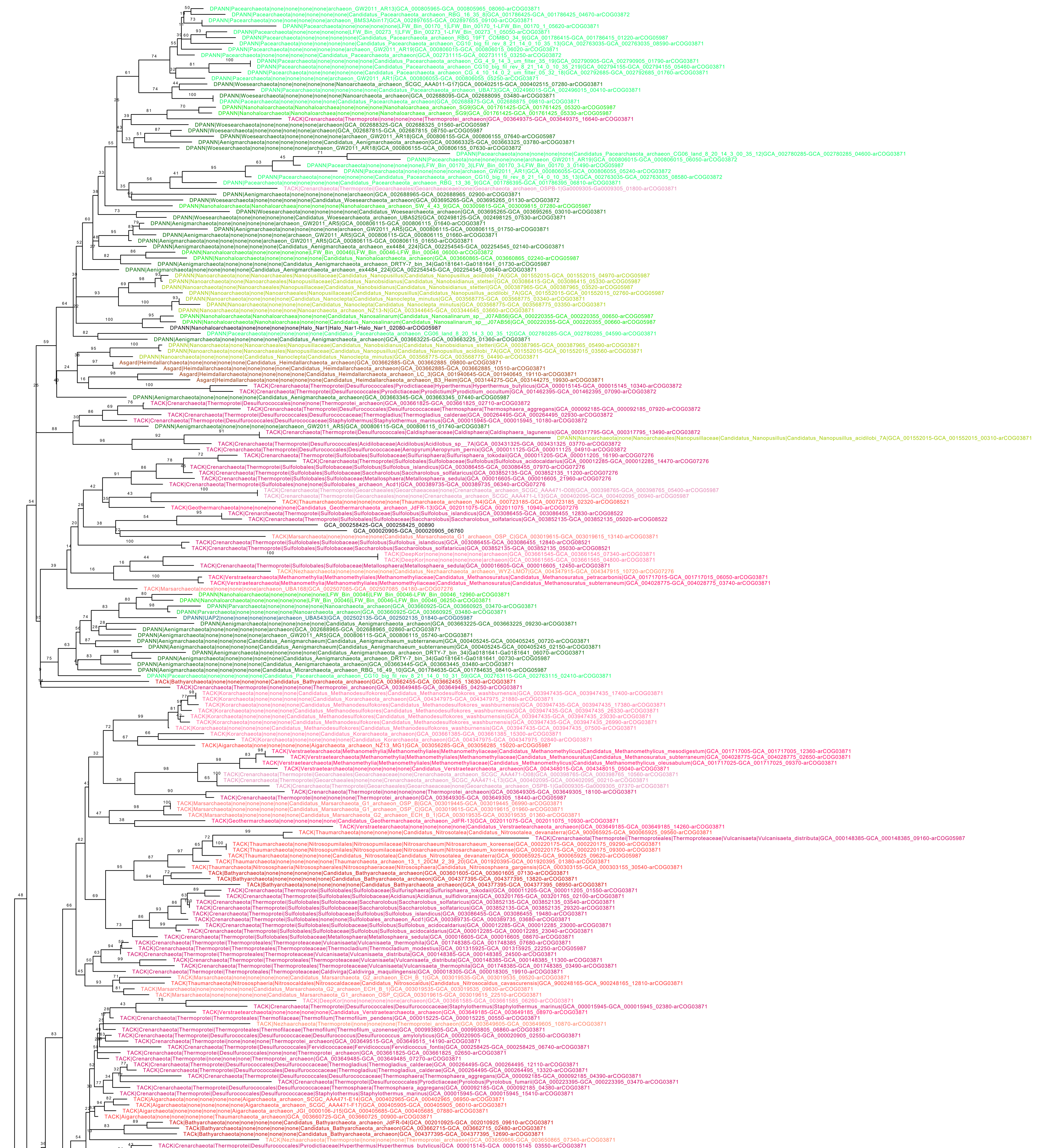

**Extended Data Fig. 26. Phylogenetic placement of all proteins annotated with the pilus associated COG03430.** Gene sequences were extracted from a set of reference archaeal and bacterial genomes (Supplementary Table 16). Sequence alignments were produced with MAFFT L-INS-i, trimmed with BMGE, and phylogenetic analysis carried out with IQ-TREE using the LG+C10+F+R model with 1000 bootstraps. Taxa have been assigned colours at the phylum level (details of taxonomic assignment provided in Supplementary Table 16).

**Supplementary Video 1. Live cell imaging of microfluidics cultivation experiments.** Video shows phase-contrast time lapse of a *Hrr. lacusprofundi* and *Ca.* Nha. antarcticus co-culture (left panel) and a pure *Hrr. lacusprofundi* culture (right panel) grown in microfluidics chambers at 30 °C. Images were acquired on a Nikon TiE2 inverted microscope fitted with a 100× oil-immersion phase-contrast NA 1.45 objective once per hour for a period of 21 hours.

**Supplementary Video 2. Cryo-ET of *Hrr. lacusprofundi* and *Ca.* Nha antarcticus co-culture.** Video shows a cryo-electron tomogram of a *Hrr. lacusprofundi* cell with internal structures consistent with *Ca.* Nha. antarcticus cells visible in the cytoplasm (Z-slice from this tomogram is shown in Fig. 3a and b). Tilt series was acquired on a Titan Krios at 300 kV.

**Supplementary Video 3. Cryo-ET of *Hrr. lacusprofundi* and *Ca.* Nha antarcticus co-culture.** Video shows a cryo-electron tomogram of a *Hrr. lacusprofundi* cell with an internal structure consistent with a *Ca.* Nha. antarcticus cell visible in the cytoplasm (Z-slice from this tomogram is shown in Fig. 3c). Tilt series was acquired on a Titan Krios at 300 kV.

**Supplementary Video 4. Cryo-ET of *Hrr. lacusprofundi* and *Ca.* Nha antarcticus co-culture.** Video shows a cryo-electron tomogram of a *Hrr. lacusprofundi* cell with internal structures consistent with *Ca.* Nha. antarcticus cells visible in the cytoplasm. Tomogram also shows a *Ca.* Nha. antarcticus cell that appears to be in the process of internalising within the *Hrr. lacusprofundi* cell (Z-slice from this tomogram is shown in Fig. 3d - f). Tilt series was acquired on a Titan Krios at 300 kV.

**Supplementary Video 5. Cryo-ET of a pure *Ca.* Nha antarcticus sample.** Video shows a cryo-electron tomogram of a purified *Ca.* Nha. antarcticus cell used for co-culture experiments. Tomogram shows presence of internal structures resembling membrane embedded lipid droplets and PHA-like granules (Z-slice from this tomogram is shown in Extended Data Fig. 13a). Tilt series was acquired on a Titan Krios at 300 kV.

**Supplementary Video 6. Cryo-ET of a pure *Ca.* Nha antarcticus sample.** Video shows a cryo-electron tomogram of a purified *Ca.* Nha. antarcticus cell used for co-culture experiments. Tomogram shows presence of internal structures resembling membrane embedded lipid droplets and PHA-like granules (Z-slice from this tomogram is shown in Extended Data Fig. 13b). Tilt series was acquired on a Titan Krios at 300 kV.

**Supplementary Video 7. Cryo-ET of a pure *Ca.* Nha antarcticus sample.** Video shows a cryo-electron tomogram of a purified *Ca.* Nha. antarcticus cell used for co-culture experiments. Tomogram shows presence of internal structures resembling membrane embedded lipid droplets and PHA-like granules (Z-slice from this tomogram is shown in Extended Data Fig. 13c). Tilt series was acquired on a Titan Krios at 300 kV.

**Supplementary Video 8. Cryo-ET of a pure *Ca.* Nha antarcticus sample.** Video shows a cryo-electron tomogram of a purified *Ca.* Nha. antarcticus cell used for co-culture experiments. Tomogram shows presence of internal structures resembling membrane embedded lipid droplets and PHA-like granules (Z-slice from this tomogram is shown in Extended Data Fig. 13d). Tilt series was acquired on a Titan Krios at 300 kV.

**Supplementary Video 9. Cryo-ET of *Hrr. lacusprofundi* and *Ca.* Nha antarcticus co-culture.** Video shows a cryo-electron tomogram of a *Hrr. lacusprofundi* cell interacting with a *Ca.* Nha. antarcticus cell showing membrane fusion at the contact site between the two cells (Z-slice from this tomogram is shown in Extended Data Fig. 14a and b). Tilt series was acquired on a Titan Krios at 300 kV.

**Supplementary Video 10. Cryo-ET of *Hrr. lacusprofundi* and *Ca.* Nha antarcticus co-culture.** Video shows a cryo-electron tomogram of a *Hrr. lacusprofundi* cell interacting with a *Ca.* Nha. antarcticus cell showing membrane fusion at the contact site between the two cells (Z-slice from this tomogram is shown in Extended Data Fig. 14c and d). Tilt series was acquired on a Titan Krios at 300 kV.

**Supplementary Video 11. Cryo-ET of a pure *Hrr. lacusprofundi* culture.** Video shows a cryo-electron tomogram of a *Hrr. lacusprofundi* cell in pure culture showing no sign of internal structures similar to those seen in co-cultures (Z-slice from this tomogram is shown in Extended Data Fig. 15a). Tilt series was acquired on a Titan Krios at 300 kV.

**Supplementary Video 12. Cryo-ET of a pure *Hrr. lacusprofundi* culture.** Video shows a cryo-electron tomogram of a *Hrr. lacusprofundi* cell in pure culture showing no sign of internal structures similar to those seen in co-cultures (Z-slice from this tomogram is shown in Extended Data Fig. 15b). Tilt series was acquired on a Titan Krios at 300 kV.

**Supplementary Video 13. Cryo-ET of a pure *Hrr. lacusprofundi* culture.** Video shows a cryo-electron tomogram of a *Hrr. lacusprofundi* cell in pure culture showing no sign of internal structures similar to those seen in co-cultures (Z-slice from this tomogram is shown in Extended Data Fig. 15c). Tilt series was acquired on a Titan Krios at 300 kV.

**Supplementary Video 14. Cryo-ET of *Hrr. lacusprofundi* and *Ca.* Nha antarcticus co-culture.** Video shows a cryo-electron tomogram of a *Hrr. lacusprofundi* cell that has lysed with an internal structure consistent with a *Ca.* Nha. antarcticus cell visible in the cytoplasm (Z-slice from this tomogram is shown in Extended Data Fig. 16a and b). Tilt series was acquired on a Titan Krios at 300 kV.

**Supplementary Video 15. Cryo-ET of *Hrr. lacusprofundi* and *Ca.* Nha antarcticus co-culture.** Video shows a cryo-electron tomogram of a *Hrr. lacusprofundi* cell with internal structures consistent with *Ca.* Nha. antarcticus cells visible in the cytoplasm (Z-slice from this tomogram is shown in Extended Data Fig. 16c and d). Tilt series was acquired on a Titan Krios at 300 kV.
